# Global change drivers and the risk of infectious disease

**DOI:** 10.1101/2022.07.21.501013

**Authors:** Michael B. Mahon, Alexandra Sack, O. Alejandro Aleuy, Carly Barbera, Ethan Brown, Heather Buelow, David J. Civitello, Jeremy M. Cohen, Luz de Wit, Meghan Forstchen, Fletcher W. Halliday, Patrick Heffernan, Sarah A. Knutie, Alexis Korotasz, Joanna G. Larson, Samantha L. Rumschlag, Emily Selland, Alexander Shepack, Nitin Vincent, Jason R. Rohr

**Affiliations:** Department of Biological Sciences, University of Notre Dame; Notre Dame, Indiana, USA; Environmental Change Initiative, University of Notre Dame; Notre Dame, Indiana, USA; Eck Institute of Global Health, University of Notre Dame; Notre Dame, Indiana, USA; Department of Biology, Emory University; Atlanta, Georgia, USA; Department of Ecology and Evolutionary Biology, Yale University, New Haven, CT, USA; Department of Evolutionary Biology and Environmental Studies, University of Zurich; Zurich 8057, Switzerland; Department of Ecology and Evolutionary Biology, Institute for Systems Genomics, University of Connecticut; Storrs, CT USA

## Abstract

Anthropogenic change is contributing to the rise in emerging infectious diseases, but it remains unclear which global change drivers most increase disease and under what contexts. We amassed a dataset from the literature that includes 1,832 observations of infectious disease responses to global change drivers across 1,202 host-parasite combinations. We found that biodiversity loss, climate change, and introduced species were associated with increases in disease-related endpoints or harm (i.e., enemy release for introduced species), whereas urbanization was associated with decreases in disease endpoints. Natural biodiversity gradients, deforestation, forest fragmentation, and most classes of chemical contaminants had non-significant effects on these endpoints. Overall, these results were consistent across human and non-human diseases. Context-dependent effects of the global change drivers on disease were common and are discussed. These findings will help better target disease management and surveillance efforts towards global change drivers that increase disease.

**One-Sentence Summary:** Here we quantify which global change drivers increase infectious diseases the most to better target global disease management and surveillance efforts.

## Main Text

Emerging infectious diseases are on the rise, often originate from wildlife, and are significantly correlated with socio-economic, environmental, and ecological factors (*1*). Consequently, there is concern that anthropogenic global change is contributing to alterations in disease risk. For example, several studies have demonstrated that infectious disease risk is modified by changes to biodiversity (*2-5*), climate change (*6-9*), and chemical pollution (*10-12*). Landscape transformations, such as forest conversion to agriculture or urban centers, also regularly shift disease risk (*13-16*). Additionally, the movement of people, products, and animals around the planet has resulted in pathogen introductions with massive health consequences to humans and wildlife (*17*). Mechanistically, global changes can alter disease by affecting the distribution of epidemiological traits in ecological communities, modulating immune defenses, and altering contact rates among parasites, wildlife, livestock and humans. As an example, the COVID-19 pandemic, which has reshaped the global economic and public health landscape, has been linked to animal trade and global travel, as well as urbanization, climate change, air pollution, and habitat loss (*18*). It has also undoubtedly heightened interest in understanding causes of disease outbreaks and investment in infectious disease control, mitigation, and surveillance.

While there are many individual studies on infectious disease risk and environmental change, as well as syntheses on how individual drivers of ecosystem change affect infectious diseases (*1-17*), no quantitative syntheses exist that examine how infectious disease risk is modified across global change drivers (*19*). This literature gap is critical to fill because resources for infectious disease management will always be limited and could be poorly targeted without knowledge of which global change drivers most affect infectious disease risk. Moreover, risk might only be high for certain types of pathogens or hosts, for wildlife but not human diseases, or for certain ecological conditions. As an example, the emergence of zoonotic diseases of humans tends to be driven more by interactions with particular mammalian and avian taxa than other vertebrate groups (*20*). Thus, understanding these context dependencies will further enhance the efficacious use of limited resources for disease control.

Here, our primary goal is to use a traditional meta-analytical approach to determine the magnitude with which global change drivers are associated with infectious disease risk and whether these associations depend on ecological contexts, such as host or parasite taxon or human versus non-human diseases. To accomplish these goals, we conducted a literature search to identify studies on infectious disease that considered at least one of the five major drivers of global change highlighted by the Millennium Ecosystem Assessment (*21*): biodiversity change and overexploitation, climate change, chemical pollution, habitat loss/change, or introduced species (see Methods). Although we treat all global drivers as independent variables in our analyses, we acknowledge that they can be interdependent. For example, climate change and chemical pollution can cause habitat loss and change, which in turn can cause biodiversity loss and facilitate species introductions.

The database resulting from our literature search includes 579 studies and 1,832 observations of global change drivers on disease or parasitism from 909 parasite taxa, 380 host taxa, and 1,202 host-parasite taxa combinations. Each observation in the database contains information on the associated global change driver and host and parasite taxa and traits (e.g. a human and non-human parasite), and whether it was derived from freshwater, marine, or terrestrial systems and laboratory or field studies (Fig. 1, Fig. S1-S5, Table S2). Additionally, each response variable was classified as a host endpoint, which captures host symptoms or consequences of infection (host disease, survival, growth, and reproduction) or a parasite endpoint, which captures parasite abundance in hosts (parasite prevalence, incidence, abundance, survival, growth, and richness). Hedge’s g effect sizes were calculated from each study, with positive and negative values representing increases and decreases in disease, respectively. The exception was for studies on introduced species, for which decreases and increases in parasites or disease in the native host received negative and positive values, respectively, and decreases and increases in parasites or disease in non-native hosts were given positive and negative values, respectively. Thus, a reduction in disease in non-native species received the same effect size direction as an increase in disease in native hosts because both were deemed potentially detrimental (see Supplement for further discussion). We addressed non-independence among observations by treating study as a random intercept in all analyses (see Methods).

**Fig. 1.**
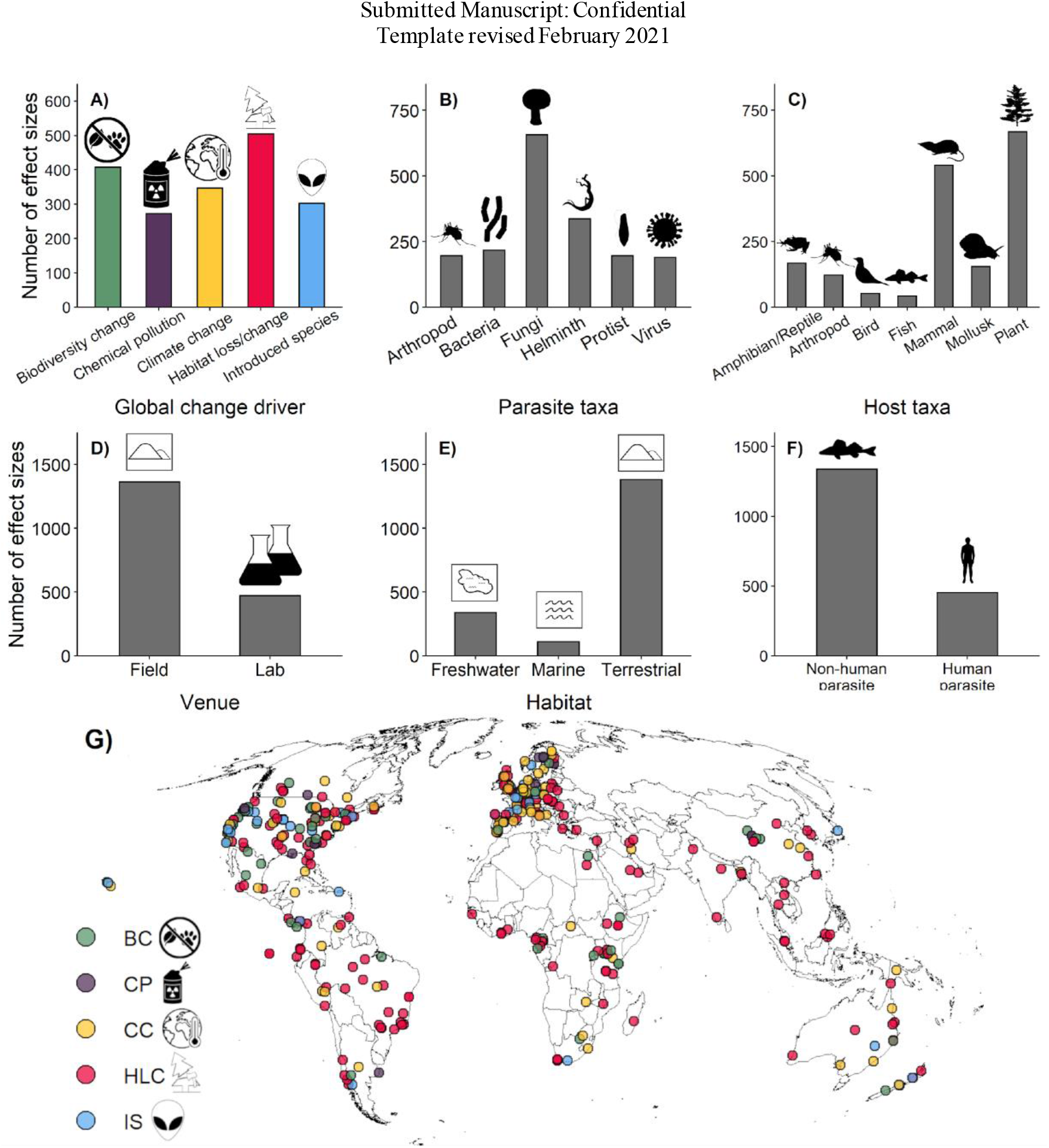
Summary of the number of observations (i.e., effect sizes) in the infectious disease database across ecological contexts. The contexts are (**A**) global change driver, (**B**) parasite taxa, (**C**) host taxa, (**D**) experimental venue, (**E**) habitat of the study, and (**F**) human parasite status. Locations of field studies (**G**) show broad global coverage of studies included in the database. See Fig. S1 and S2 for similar figures but where the response variables are the number of studies and number of parasite taxa, respectively. See Fig. S3-S5 for the number of observations, studies, and parasite taxa in the database partitioned by ecto- and endoparasites, ecto- and endothermic hosts, vectors and non-vectors, vector-borne and non-vector-borne parasites, complex and direct transmission parasites, parasites with and without free-living stages, parasites that do and do not infect humans, and micro- and macroparasites.

Among the global change drivers, habitat loss/change caused marginally non-significant reductions in disease, while biodiversity change, climate change and introduced species increased disease responses or disease-related harm (i.e., fewer enemies for introduced species). Chemical pollution was intermediate to these two groups, with non-significant effects on disease endpoints. Biodiversity change was associated with a 108% greater increase in disease than climate change and a 28% greater increase in disease than introduced species (Fig. 2). Importantly, we found no evidence that effect size patterns among global change drivers could be explained by differences in variances or sample sizes among global change drivers (Fig. S6). Further, forest plots showed that extreme observations were uncommon in our dataset and were typically given low weight in our meta-analytical models, suggesting that these data had minimal impacts on patterns among global change drivers (Fig. S7). Egger’s test indicated no evidence for publication bias (small-study effect; p > 0.05; Fig. S8) when using multi-level meta-regression. Additionally, we documented a time-lag bias (e.g., stronger positive effects earlier in the dataset; *p* < 0.001; Fig. S9), but the effect was likely driven by only positive effects in the first 20 years, which represented <5% of the dataset. Thus, while statistically significant, the time-lag effect seems to be small. Finally, our results were robust to the iterative exclusion of individual studies (Fig. S10). Given the results of these tests of bias and robustness, patterns across global change drivers do not appear to be artefactual.

**Fig. 2.**
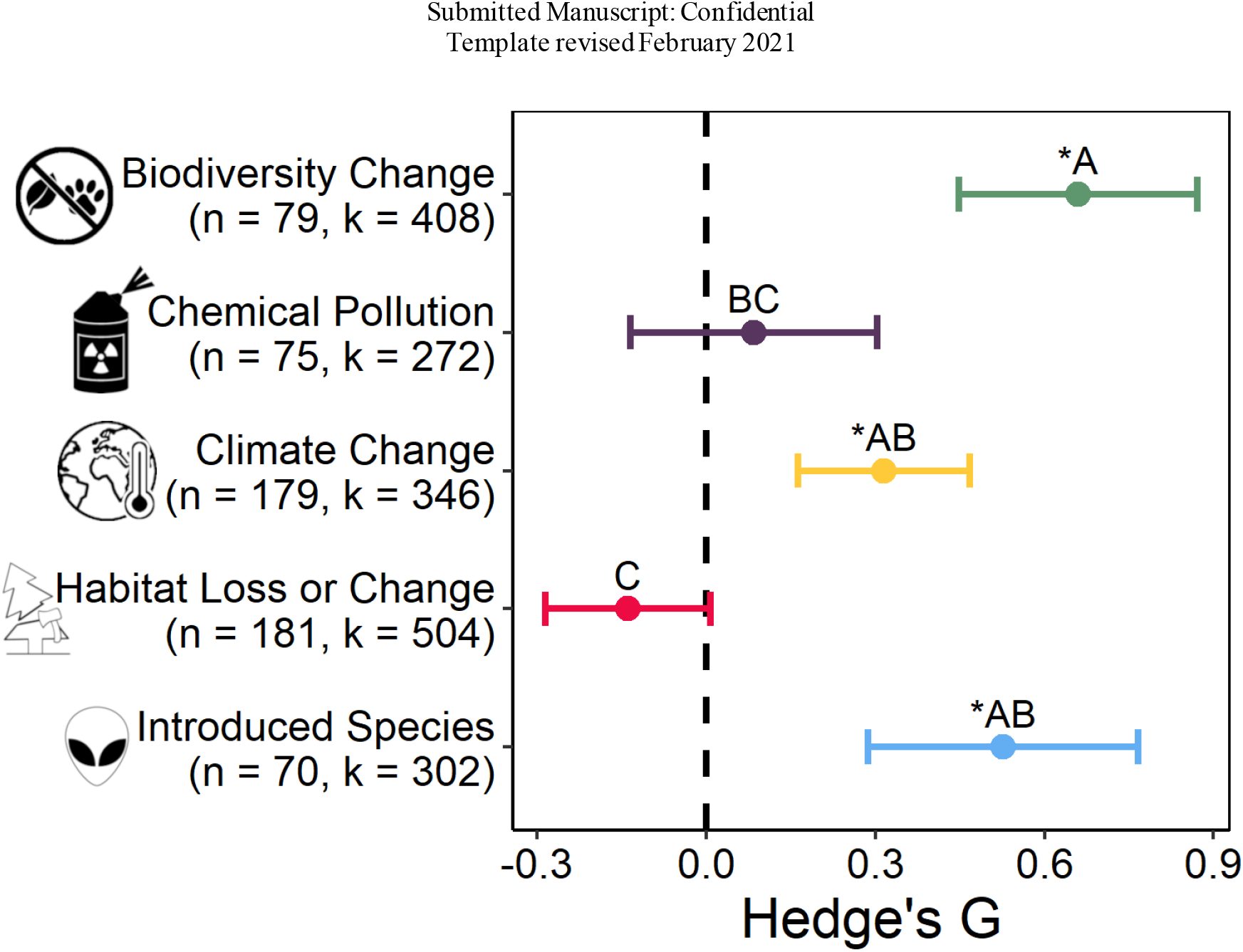
The effects of five common global change drivers on infectious disease responses. Biodiversity change, climate change, and introduced species are associated with increases in disease-related endpoints or harm (i.e., introduced species having less parasites). Habitat loss or change was associated with non-significant decreases in disease endpoints. Chemical pollution was associated with non-significant differences in disease. Shown in parentheses are the number of studies (n) and effect sizes (k) for each driver. The displayed points represent the mean predicted values (with 95% confidence intervals) from a meta-analytical model with separate random intercepts for study. Global change driver effects are significant (indicated by asterisks; p < 0.05) when confidence intervals do not overlap with zero. Points that do not share letters are significantly different from one another based on a Tukey’s posthoc multiple comparison test.

Next, we evaluated global change driver subcategories (Fig. 3). Consistent with previous studies (*3*), the loss of pre-existing biodiversity was associated with significantly greater increases in infectious disease outcomes (81% more) than natural biodiversity gradients (e.g., latitudinal or elevational gradients in species richness) (Fig. 3). Enemy release (i.e., the notion that introduced species leave many of their parasites behind in their native range) reduced infectious diseases in introduced species to a similar magnitude as biodiversity loss increased infectious diseases. Independent studies revealed that mean temperature and carbon dioxide similarly increased disease but had weaker effects than biodiversity loss (55% and 47% weaker, respectively) and enemy release (45% and 35% weaker, respectively). Urbanization decreased infectious diseases, perhaps because urban development is associated with improved water, sanitation, and hygiene for humans and habitat loss for many parasites and their non-human hosts (*16*). All other subcategories had non-significant effects on disease (Fig. 3). Given the limited funds for infectious disease management, these results suggest that controlling or mitigating biodiversity loss, climate change, and introduced species might be particularly important for infectious disease management.

**Fig. 3.**
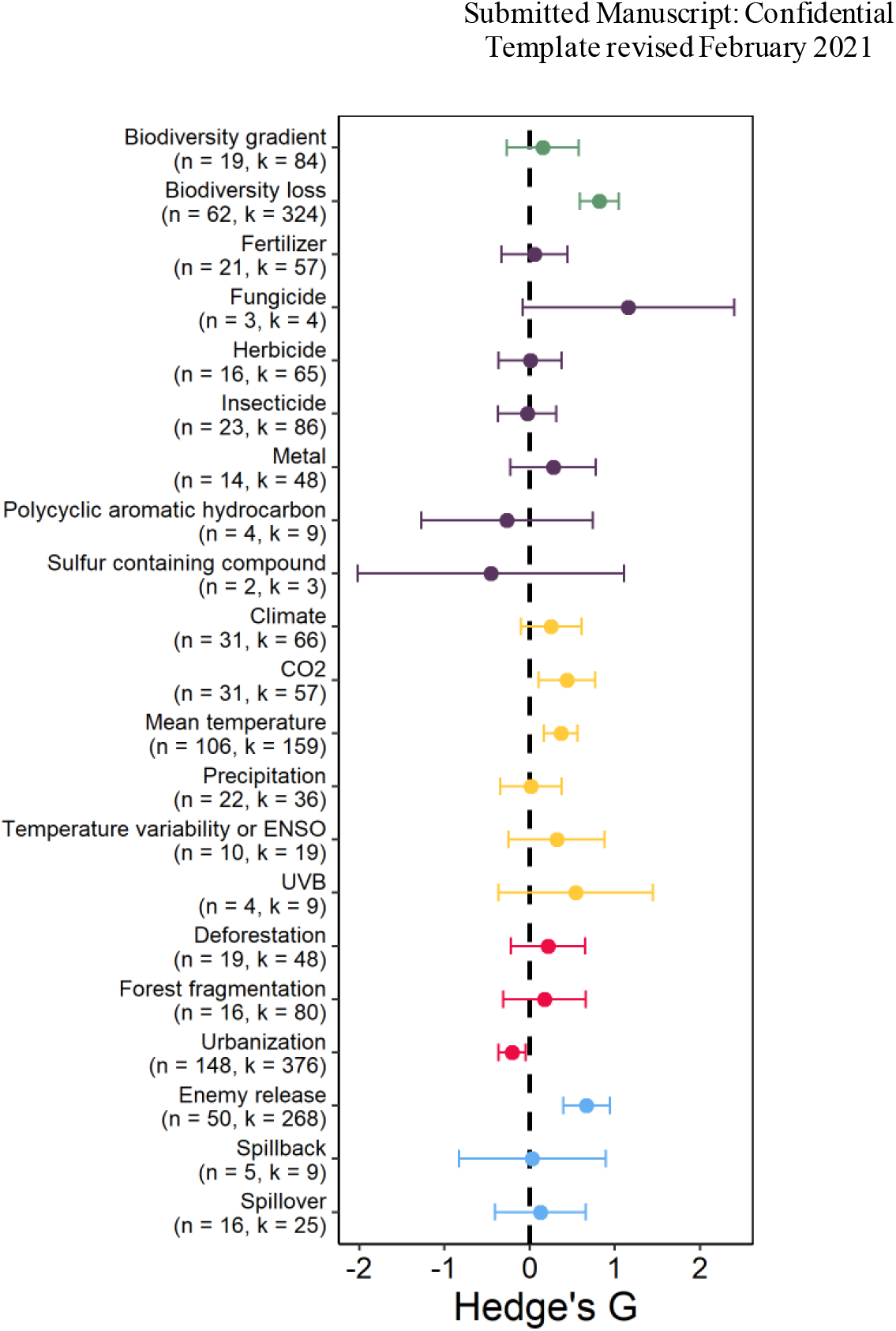
The effects of subcategories within five common global change drivers on mean infectious disease responses in the literature. Shown in parentheses are the number of studies (n) and effect sizes (k) of each subcategory. The displayed points represent the mean predicted values (with 95% confidence intervals) from a meta-analytical model with study as a random intercept. Confidence intervals that do not overlap with zero are generally significant (*p*<0.05), but see text for details. Spillover is defined as spread of a parasite from an introduced to native host or from a native to introduced host (*17*). Spillback occurs when an introduced host is a competent host for a native parasite and amplifies infections in the native host (*17*). Enemy release is defined as cases when an introduced host has fewer parasites in its introduced range than native range or than native species in its introduced range (*17, 24, 26*). Forest fragmentation compares different sizes of forest patches whereas deforestation compares forests to the absence of forests (i.e. two different habitats). Biodiversity gradient covers natural variation in biodiversity (e.g. across latitude or elevation), whereas biodiversity loss is a loss of biodiversity usually associated with an anthropogenic factor (*3*). UVB=ultraviolet radiation B, ENSO=El Niño-Southern Oscillation.

Understanding context dependencies is also crucial for properly targeting limited resources for disease control. Although the ideal approach would have been to compare global change drivers in a single model-selection analysis that considered the correlations among all independent variables and their interactions, this approach was not possible due to missing cells. For instance, all habitat loss/change studies were conducted in freshwater and terrestrial systems (i.e., no lab or marine studies), virtually all fungi were ectoparasitic, and some host taxa were too infrequently tested under certain global change drivers (see Table S2). We circumvented these statistical limitations using two approaches. First, we tested for two-way interactions between each global change driver and host and parasite taxa and various traits of hosts, parasites, and studies. Second, to account for the covariances among predictors and to identify the most parsimonious combinations of predictors, we fit models with all possible combinations of main effects of host, parasite, and study factors for each driver separately (see Methods and Supplemental Materials RMarkdown file). We then qualitatively assessed the consistency in the results between these statistical approaches.

Importantly, these analyses can reveal both when there are and are not context dependencies. For instance, there were many consistent patterns across global change drivers. As an example, for every global change driver, parasite versus host endpoint was an important moderator in either the two-way interaction (Fig. 4A) or model selection analyses (Fig. 4B), and in each case, parasite endpoints were more sensitive to the global change drivers (many times significantly so) than host endpoints (see Fig. S11). This is likely because host endpoints are capturing host symptoms or consequences of infection, whereas parasite endpoints are capturing parasite abundance in hosts. Given that the abundance of parasites can change profoundly without changes in symptoms or disease, especially for hosts that employ tolerance (i.e., ameliorating the damage that infection causes) rather than resistance (i.e., “fighting” the parasite directly) defense strategies (*22, 23*), it is unsurprising that parasite endpoints are more sensitive to global change factors than are host endpoints.

**Fig. 4.**
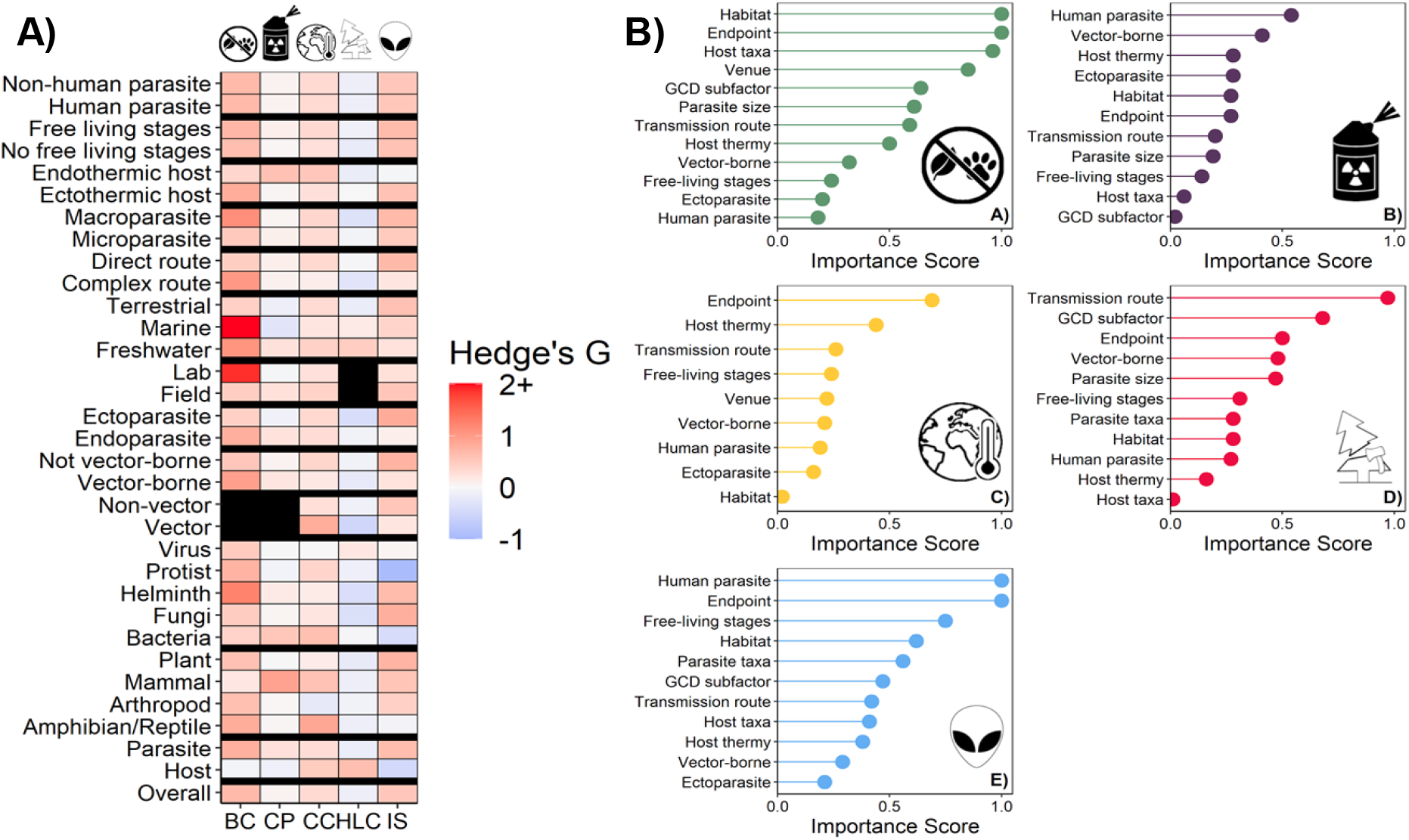
Results of analyses testing for context dependent effects of global change drivers on infectious disease responses. (**A**) Coefficients from separate tests of two-way interactions between each global change driver and host and parasite taxa and various traits of hosts, parasites, and studies. Black sections of the heat map could not be tested because of missing data. (**B**) Relative importance scores from model selection examining the effects of five common global change drivers on mean infectious disease responses. Unlike the two-way interaction analyses, the model selection analyses account for the covariances among predictors and identify the most parsimonious combinations of predictors. See RMarkdown file for the coefficients from these models and Figures 1 or 2 for definitions of the cartoon drawings. Endpoint: host or parasite; Free-living stage: free-living stage or not; GCD subfactor: see Fig. 3; Habitat: freshwater, marine, terrestrial; Host taxa: see Fig. 1; Host thermy: ecto- or endotherm; Human parasite: human parasite or not; Parasite size: macro- or microparasite; Parasite taxa: see Fig. 1; Transmission route: complex or direct; Vector-borne: vector- borne or not; Venue: lab or field; Ectoparasite: ecto- or endo-parasite.

The effects of global change drivers on infectious disease outcomes also did not significantly depend on host taxon or whether the parasite infected humans or not (Fig. 4). These results indicate that global change drivers are having consistent effects on infectious disease risk across broad host taxa, including humans. The one exception was for introduced species, as non-human parasites, on average, responded more strongly to introduced species and enemy release than human parasites (Fig. 4). This is likely because human parasites in the meta-analysis must be generalist parasites (because humans are never introduced hosts in these studies because of their global distribution and thus the parasites must infect human and non-human hosts), which do not suffer as strongly from enemy release as host specialists (*24*). Finally, no clear context dependencies emerged from studies on chemical pollution (no significant two-way interactions and most relative importance scores <0.5; Fig.4). This result was likely because chemical pollution had the fewest effect sizes (Fig. 1) and enormous diversity in the pollutants tested, making it challenging to uncover consistent patterns on infectious disease and highlighting the need for further infectious disease research on this global change driver, especially given that many contaminants can be immunosuppressive (*12*).

In contrast to these consistent patterns across global change drivers, numerous context dependencies were also detected. For example, when compared to viruses, fungi responded more positively to introduced species, and helminths responded more negatively to habitat loss/change. Helminths also responded more positively to biodiversity loss than all other parasite taxa (Fig. S12). Relative to non-vectors, vectors increased with climate change and decreased with habitat loss/change (Fig. S13), results consistent with studies highlighting that climate change typically increases vector-borne diseases (*7, 9*) and that urbanization often decreases vector-borne diseases, especially for those diseases that are not associated with container-breeding vectors (*16, 25*). Similarly, relative to parasites with simple (i.e., direct) life cycles, parasites with complex life cycles, such as vectored parasites and macroparasites, experienced greater decreases and increases when exposed to habitat loss/change and biodiversity loss, respectively, results that were generally similar across the two-way interaction and model selection analyses (Fig. S14-16, Fig. 4). For parasites with complex life cycles, only one of the host species needs to be sensitive to habitat loss/change to disrupt transmission; thus, it is not surprising that habitat loss/change is associated with greater declines of parasites with complex than direct life cycles. Moreover, the biodiversity change results are consistent with a meta-analysis highlighting that biodiversity loss increases parasites more if they have complex (such as vector-borne parasites) than simple life cycles (*2*). Biodiversity loss also increased disease more in laboratory than field studies (Fig. S17), in aquatic than terrestrial systems (Fig. S18), and in ectothermic than endothermic hosts (Fig. S19). Additionally, ectoparasites decreased more than endoparasites with habitat loss/change (Fig. S20), likely because ectoparasites are less protected than endoparasites from the changes in abiotic conditions associated with habitat loss/change.

Here we revealed that biodiversity change, climate change and introduced species increased disease responses or disease-related harm, while habitat loss/change caused marginally non-significant reductions in disease and chemical pollution had non-significant effects on disease endpoints. Within these broad global change drivers, the subcategories of biodiversity loss, temperature, CO_2_, and enemy release generally increased infectious disease endpoints or disease-related harm (i.e., less enemies for introduced species), whereas natural biodiversity gradients, deforestation, fragmentation, other categories of climate change, and all classes of chemical contaminants had non-significant effects on these endpoints. Conversely, urbanization reduced disease endpoints. All of these results were generally consistent across human and non-human diseases. Context dependencies were somewhat common. Endpoints from parasites with complex life cycles, such as macroparasites and vector-borne pathogens, decreased more with habitat loss/change and increased more with biodiversity loss and climate change than parasites with simple life cycles, and ectoparasites declined more with habitat loss/change than endoparasites. Future studies should more thoroughly explore the interdependencies and relative contributions of global change factors to disease risk. Our analyses should facilitate disease control, mitigation, and surveillance efforts globally, ultimately improving wildlife and human health.

## Acknowledgments

We thank Charles Mitchell for contributing data on enemy release, and Lauren Albert and Brin Shayhorn for assisting with data collection, and Jessica Gurevitch, Marc Lajeunesse, and Gavin Stewart for providing comments on an earlier version of this manuscript.

## Funding

National Science Foundation DEB 2109293 (JRR)

National Science Foundation DEB-2017785 (JRR)

National Science Foundation DEB-1518681 (JRR)

National Science Foundation IOS-1754868 (JRR)

National Institutes of Health R01TW010286 (JRR)

US Department of Agriculture 2021-38420-34065 (JRR)

USGS Powell Grant (JRR, SLR)

University of Connecticut Start-up funds (SAK)

National Science Foundation IOS-1755002 (DJC)

National Institutes of Allergy and Infectious Diseases NIH R01 AI150774 (DJC)

## Author contributions

Conceptualization: JRR

Methodology: all authors

Investigation: all authors

Visualization: MBM

Funding acquisition: JRR, SAK, SLR

Project administration: JRR

Supervision: JRR

Writing – original draft: JRR, MBM

Writing – review & editing: all authors

## Data and materials availability

All data and code for all analyses are available in the main text or the supplementary materials.

## Supplementary Materials

Materials and Methods

Figs. S1 to S17

Tables S1 to S2

Data S1 to S2

