## Supplementary material for "Global change drivers and the risk of infectious disease": Fig. S

### *Science* Supplementary Materials Template Instructions

This is the *Science* template for presenting and formatting your supplementary materials. To organize your supplementary materials section, please follow the instructions below. Once formatted, you should delete this first page of instructions.

**Overview:**
Supplementary materials present additional information in support of the conclusions of your paper, such as a description of the materials and methods, controls, or tabulated data presented in Tables or Figures. It will generally consist of one integrated PDF file. Audio or movie files or large data Tables can be presented as separate files. Further information is available at: http://www.sciencemag.org/authors/instructions-preparing-initial-manuscript#format-supplemental.

This section should not be used for additional discussion, analysis, or interpretations. It is not to be used as a forum to critique other publications.

References can be cited in the supplementary text section. These should be cited in order following the references in the main text as per *Science* style (i.e., italicized number in parentheses). One full reference list should be provided for all cited references at the end of the main paper.

**Using the Template**
Paste the title, author list, and corresponding author email address(es) from the main text file onto the cover page. On the cover page, complete the relevant description of the SM and delete text that does not apply.

Copy and paste relevant text into each appropriate section of the template.

Each Figure or Table should be on a separate page and can be placed above each caption. To add additional captions, simply copy and paste (repeatedly) the last caption template. Large Tables that extend beyond the width of the page should be provided as separate files in an appropriate spreadsheet format (.xlsx or similar).

Large amounts of text can be grouped by subheads. To repeat subheads, simply copy the subhead and repeat/rename.

### Supplementary Materials for

Global change drivers and the risk of infectious disease

Michael B. Mahon, Alexandra Sack, O. Alejandro Aleuy, Carly Barbera, Ethan Brown, Heather Buelow, David J. Civitello, Jeremy M. Cohen, Luz de Wit, Meghan Forstchen, Fletcher W. Halliday, Patrick Heffernan, Sarah A. Knutie, Alexis Korotasz, Joanna G. Larson, Samantha L. Rumschlag, Emily Selland, Alexander Shepack, Nitin Vincent, Oscar Aleuy Young, Jason R. Rohr

**This PDF file includes:**

Materials and Methods
Supplemental Discussion
Figs. S1 to S17
Tables S1 to S2
Data S1 to S2
Caption for Data S1: Data and metadata associated with this publication,
Caption for Data S2: RMarkdown code and output associated with this publication

**Other Supplementary Materials for this manuscript include the following:**

Data S1 to S2 [Data and metadata associated with this publication, RMarkdown code and output associated with this publication]

**Materials and Methods**

We conducted literature searches in Web of Science on each of the five global change drivers and infectious disease (see Table S1 for dates of searches and search terms). We screened papers to determine whether they drew clear conclusions about the impact of the global change driver on a parasite (e.g. parasite growth, prevalence, abundance, intensity, virulence) or host endpoint (e.g. disease, growth rate, survival) via experiments or field studies (see Supplementary Text). Each of the five global change drivers were further categorized into subcategories, which are provided in Fig. 3. From each study, we extracted data on the effect of the global change driver on each infectious disease endpoint, the subcategory of global change driver, the host and parasite species, and various traits of the study, hosts, and parasites.

The list of studies associated with biodiversity change was based Halliday *et al.* (*3*), which combined studies from four meta-analyses (see Table S1 and Halliday *et al.* (*3*) for details). Each meta-analysis included only studies that reported a measure of host biodiversity as the independent variable (e.g., host richness, Shannon diversity, Simpson diversity). For chemical pollution, contaminants were assigned to one of 11 contaminant classes provided by Newman (*27*), and we excluded papers that evaluated the development of a treatment for a parasitic infection or evaluated how naturally occurring nutrients influence disease development. Studies on introduced species focused on enemy release, spillover of parasites from native to invasive or invasive to native hosts, dilution effects, and amplification effects via parasite spillback (see Fig. 3 for definitions of these terms). Initial study list and related information were compiled by DC, JC, FH, SK, SR, and JR.

*Data extraction and effect size calculations*

We extracted mean values with associated sample sizes and dispersion (e.g., variance, standard deviation, standard error). Data extraction was performed by OY (2.4%), AK (2.3%), AS (3.4%), CB (4.3%), ES (5.9%), EB (5.7%), HB (3.4%), LdW (3.6%), LS (12.5%), MF (6.9%), MM (19.8%), NV (4.3%), and PH (3.3%) and all data were checked for accuracy by MM and AS. All data in BC was taken from Halliday *et al.* (*3*). Data presented in text or tables were directly extracted. Data from figures were digitized using *WebPlotDigitizer (28).* When available, raw data were used to calculate mean, dispersion, and sample size. When studies presented statistics other than mean values (i.e., odds ratio; regression coefficient; correlation coefficient; t-, z-, $\chi^2$-, and f-statistics), we extracted these values and their subsequent dispersion in place of mean values. Additionally, for data where disease endpoint was measured through time, a natural cubic spline relationship was generated between time and disease endpoint for each treatment and area under the curve (AUC) was then calculated for each natural cubic spline. The resulting AUC and associated error were used for the mean and dispersion to calculate an effect size.

We defined our effect size using Hedge's g

$$g = \frac{\underline{y}_1 - \underline{y}_2}{s_p} \quad \text{(equation 1)},$$

where $\underline{y}_1$, $\underline{y}_2$, and $s_p$ are the mean of sample 1, the mean of sample 2, and the pooled standard deviation, respectively. The pooled standard deviation is

$$s_p = \sqrt{\frac{(n_1-1)s_1^2+(n_2-1)s_2^2}{(n_1-1)+(n_2-1)}} \quad \text{(equation 2)},$$

where $s_1$ and $n_1$ are the standard deviation and number of observations for sample 1, respectively, and $s_2$ and $n_2$ are the standard deviation and number of observations for sample 2, respectively. Variance for this effect size was derived by taking the square of the pooled standard deviation. When observations were statistics rather than mean values, we converted the presented statistic to Hedge's g using standard conversion equations within the *esc* R package (*29, 30*).

*Meta-analyses*

All analyses were conducted in R 4.0. (*31*). All analyses were conducted with meta-analytic multilevel mixed-effects models using the 'rma.mv' function in the *metafor* R package (*32*). Our data had multiple effect sizes from the same studies, so all meta-analytic models were fit with a study-level and observation-level random effect, to account for the non-independence of observations from the same study. Test statistics and confidence intervals for fixed effects were computed using *t* distributions. Statistical significance was assumed when 95% confidence intervals were not overlapping zero.

*Moderator variables*

First, we estimated the overall grand mean and the total heterogeneity explained by the random effect terms. Second, to test for the effects of broad global change drivers on disease, we conducted a meta-analytical model with global change driver as the moderator. Third, to test whether global change driver subfactors differentially affect disease, we conducted a meta-analytical model with the subfactors of global change drivers as the moderator. Fourth, to test for generalities in the effects of global change drivers on disease, we conducted meta-analytical models with the main and interactive moderators of global change drivers and various host, parasite, and study moderators. The various study moderators taken from each study included host taxa, parasite taxa, vector status (vector/non-vector), vector-borne status (vector-borne/non-vector-borne), parasite type (endo-/ecto-parasite), human parasite (human/non-human), transmission route (complex/direct), free living stages (yes/no), macro- vs microparasite, host thermy (endo-/ecto-thermic), experimental venue (field/lab), response variable endpoint (host or parasite focused), and habitat (freshwater/marine/terrestrial); each of these moderators were tested separately. Differences in the main and interactive effects of these moderators was assessed using the *emmeans* R package (*33*). Finally, to determine which moderators best explain disease response to specific global change drivers, we performed model selection based on AICc in which we fit all possible combinations of the main effects of the global change subfactors and various host, parasite, and study moderators using the 'dredge' function in the *MuMIn* R package (*34*). Model selection was conducted on each global change driver separately; because of different numbers of observations across global change drivers and missing cell issues within global change drivers, the replicates varied for each global change driver (biodiversity gradient $k$ = 401; climate change $k$ = 284; habitat loss/change $k$ = 400; introduced species $k$ = 260; chemical pollution $k$ = 253). Model weights and relative importance values for each predictor variable were calculated from models with a ΔAICc ≤ 4. All data, R scripts, and RMarkdown files accompany this manuscript.

*Publication bias and sensitivity analysis*

Publication bias is the selective publishing of certain research findings, such as significant or favorable results. Common publication biases include small study effects (correlation between observed effects sizes and standard errors) and time lag biases (positive results being published before negative results) (*35*). To assess these potential biases, first, we used funnel plots to visually inspect the relationship between model (intercept only) and standard error, but it is important to note that funnel plots assume minimal heterogeneity in data and, thus, should be used as a visual tool only (*36*). Second, we performed multilevel meta regressions using sampling error or sampling variance as moderators to clarify small study effects (Egger's Test; (*36*)). Third, we investigated the time-lag bias in our data using a multilevel meta-regression with publication year as the moderator (*36*). Finally, to assess the robustness of our results, we performed leave-one-out-analysis on the meta-analytical grand mean (intercept only) model. From this analysis, we determined whether the removal of a single study greatly shifted the grand mean estimate (*36*).

Given that effect size is a function of variance and sample size, differences in the distributions of effect sizes on disease endpoints among global change drivers and across contexts might be the product of these factors (*37*). To test whether differences in effect sizes were driven by differences in sample sizes and/or variances, we tested for differences in sample sizes and variances among global change drivers. We applied generalized linear mixed effects models (GLMMs; "glmer" function, *lme4* package (*38*)) with 'study' as a random intercept to compare variances and samples sizes among global change drivers, using Gaussian errors for variance models and Poisson errors for sample size models.

**Supplemental Discussion on Directionality of 'Enemy Release' Studies**

The ecological valuation of non-human species based upon their native-status is a conundrum. Discussions surrounding the human valuation of native and non-native organisms have been going on for decades, with a conservationist bias against non-native species possibly rooted in xenophobia (*39*). Moreover, given that past natural systems are undergoing transformations caused by anthropogenic global change drivers that will forever alter their landscapes, some scientists argue that the field of ecology needs to move on from the dichotomy between native and non-native species (*40*). Nonetheless, the invasion of non-native species presents a threat to native biodiversity, ecosystems as a whole, and human well-being (*41*). We recognize the potential ethical problems of valuing native species over non-native species as we have done here, but our treatment of a reduction in disease in non-native species being given the same effect size direction as an increase in disease in native hosts was based solely on the likelihood of non-native species negatively affecting ecosystems and biodiversity (*42*).

The value judgment applied to non-native species is not unlike the value judgment applied to infectious diseases and parasites. Parasitism is the most common consumer strategy among organisms. Hence, parasites consume other organisms like every heterotrophic organism on the planet. However, unlike herbivores and many other consumers, parasites and the infectious diseases that they cause are generally regarded as bad or problematic by humans. This is a value judgment just like treating non-native species as adverse.

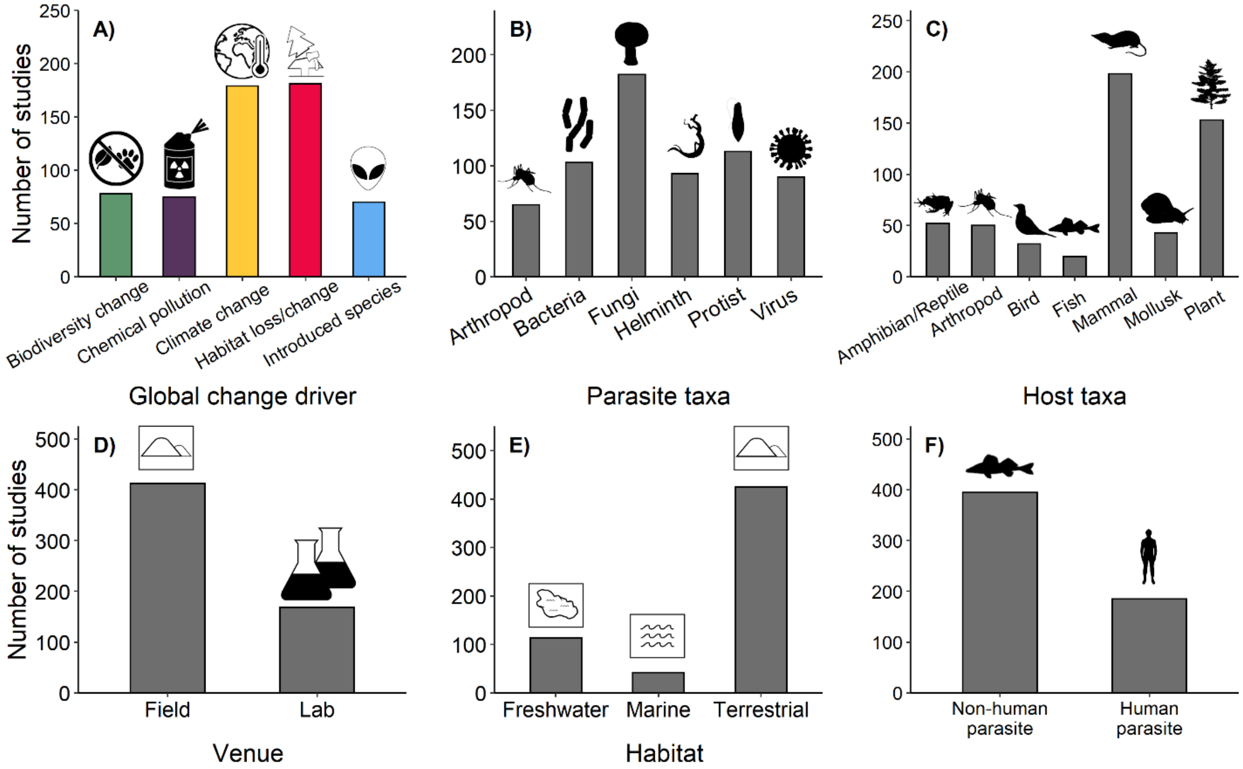


**Fig. S1. Summary of the number of studies in the infectious disease database across ecological contexts.** The contexts are **A)** global change driver, **B)** parasite taxa, **C)** host taxa, **D)** experimental venue, **E)** study habitat, and **F)** human parasite status.

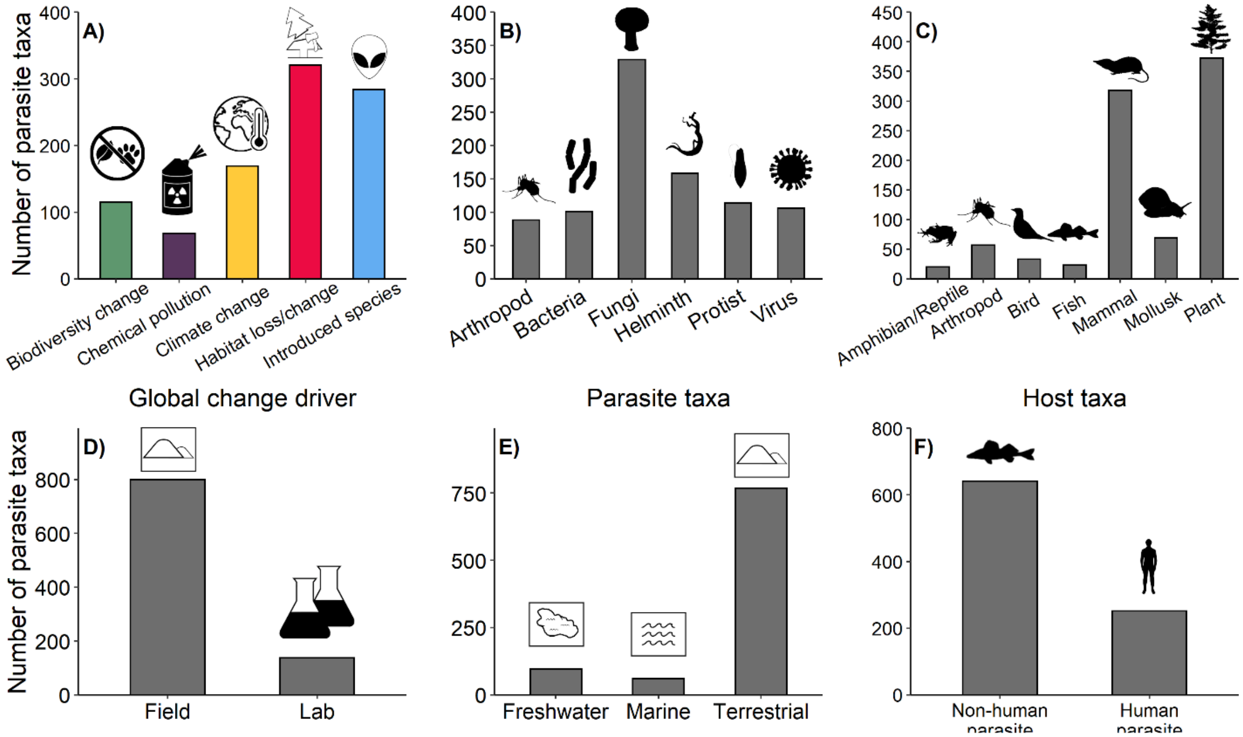


**Fig. S2. Summary of the number of parasite taxa in the infectious disease database across ecological contexts.** The contexts are **A)** global change driver, **B)** parasite taxa, **C)** host taxa, **D)** experimental venue, **E)** study habitat, and **F)** human parasite status.

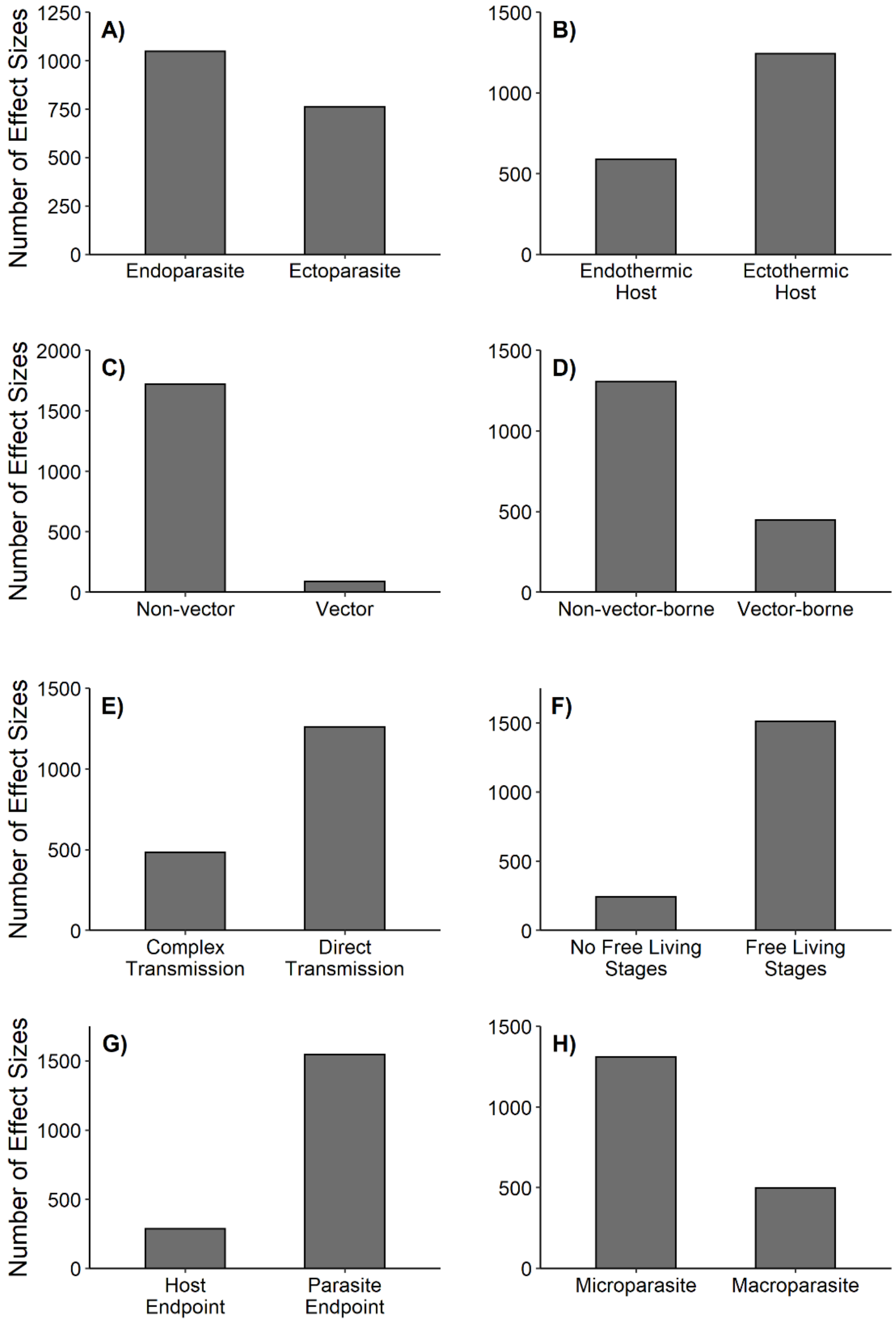


**Fig. S3. Summary of the number of effect sizes in the infectious disease database for various parasite and host contexts.** Shown are **A)** parasite type, **B)** host thermy, **C)** vector status, **D)** vector-borne status, **E)** parasite transmission, **F)** free living stages, **G)** host (e.g. disease, host growth, host survival) or parasite (e.g. parasite abundance, prevalence, fecundity) endpoint, and **H)** micro- vs macrosparasite.

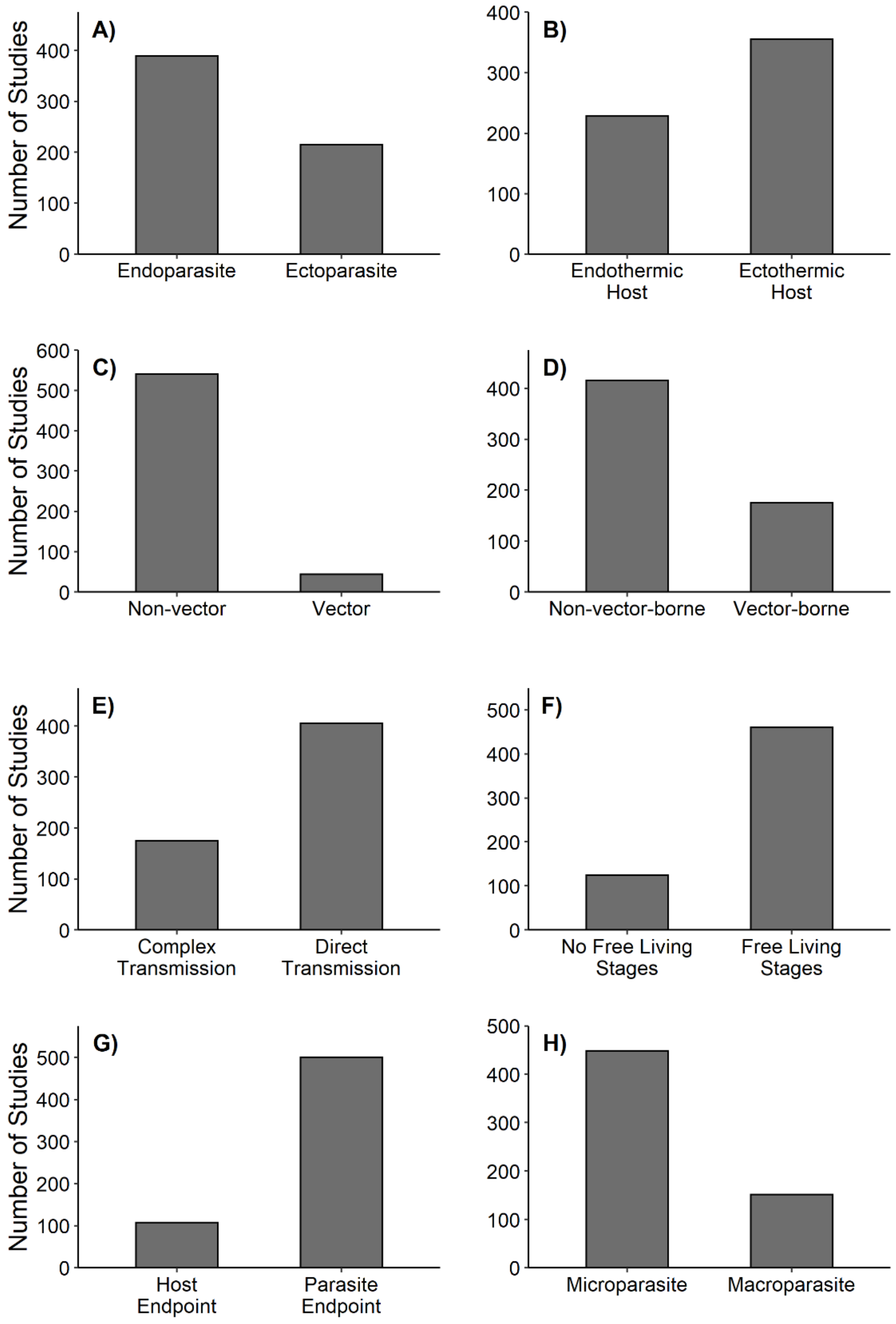


**Fig. S4. Summary of the number of studies in the infectious disease database for various parasite and host contexts.** Shown are **A)** parasite type, **B)** host thermy, **C)** vector status, **D)** vector-borne status, **E)** parasite transmission, **F)** free living stages, **G)** host (e.g. disease, host growth, host survival) or parasite (e.g. parasite abundance, prevalence, fecundity) endpoint, and **H)** micro- vs macrosparasite.

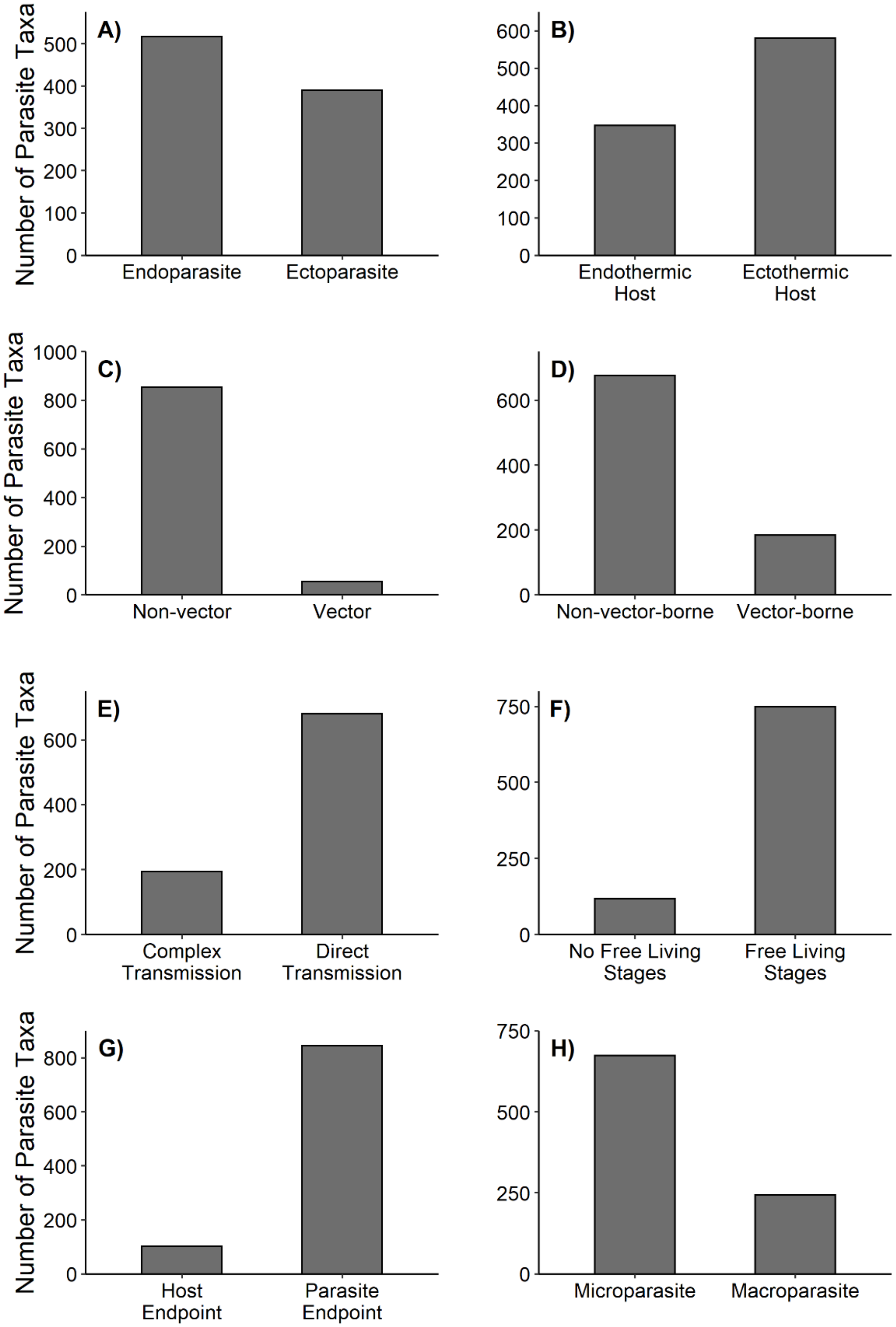


**Fig. S5. Summary of the number of parasite taxa in the infectious disease database for various parasite and host contexts.** Shown are **A)** parasite type, **B)** host thermy, **C)** vector status, **D)** vector-borne status, **E)** parasite transmission, **F)** free living stages, **G)** host (e.g. disease, host growth, host survival) or parasite (e.g. parasite abundance, prevalence, fecundity) endpoint, and **H)** micro- vs macrosparasite.

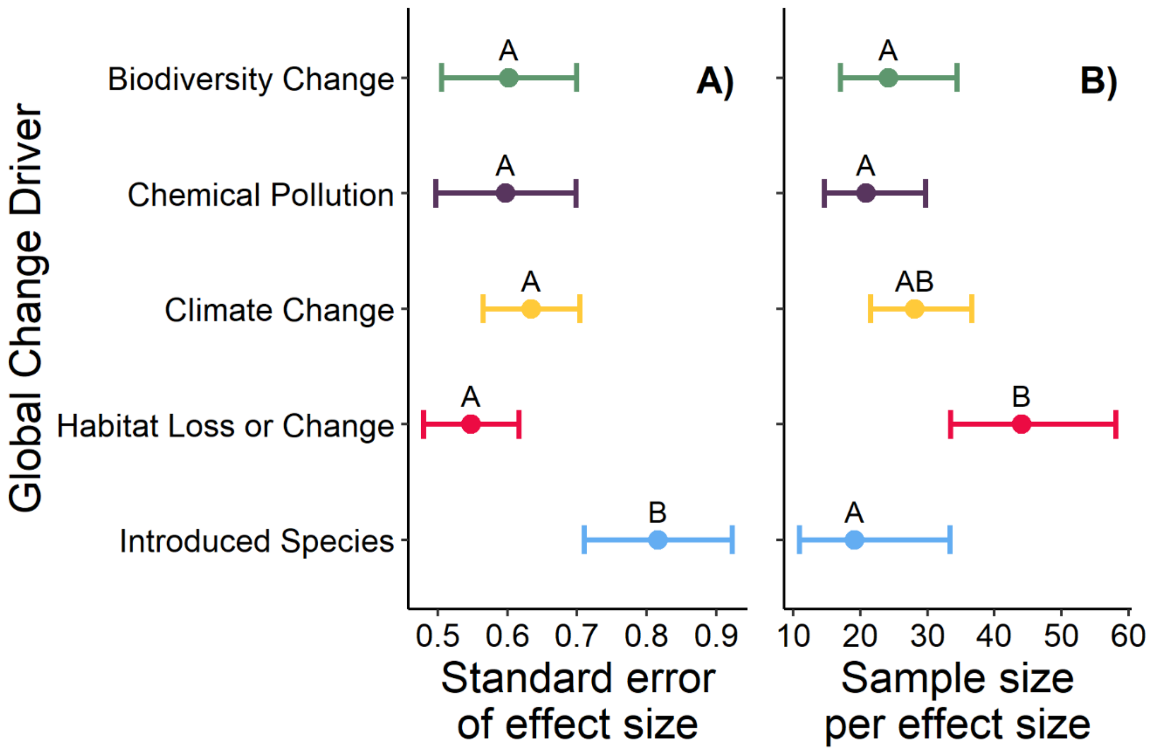


**Fig. S6. Average standard errors of the effect sizes (A) and sample sizes per effect size (B) for each of the five global change drivers.** The displayed points represent the mean predicted values (with 95% confidence intervals) from the generalized linear mixed effects models with separate random intercepts for study (Gaussian distribution for standard error model; Poisson distribution for sample size model). Points that do not share letters are significantly different from one another based on a Tukey's posthoc multiple comparison test.

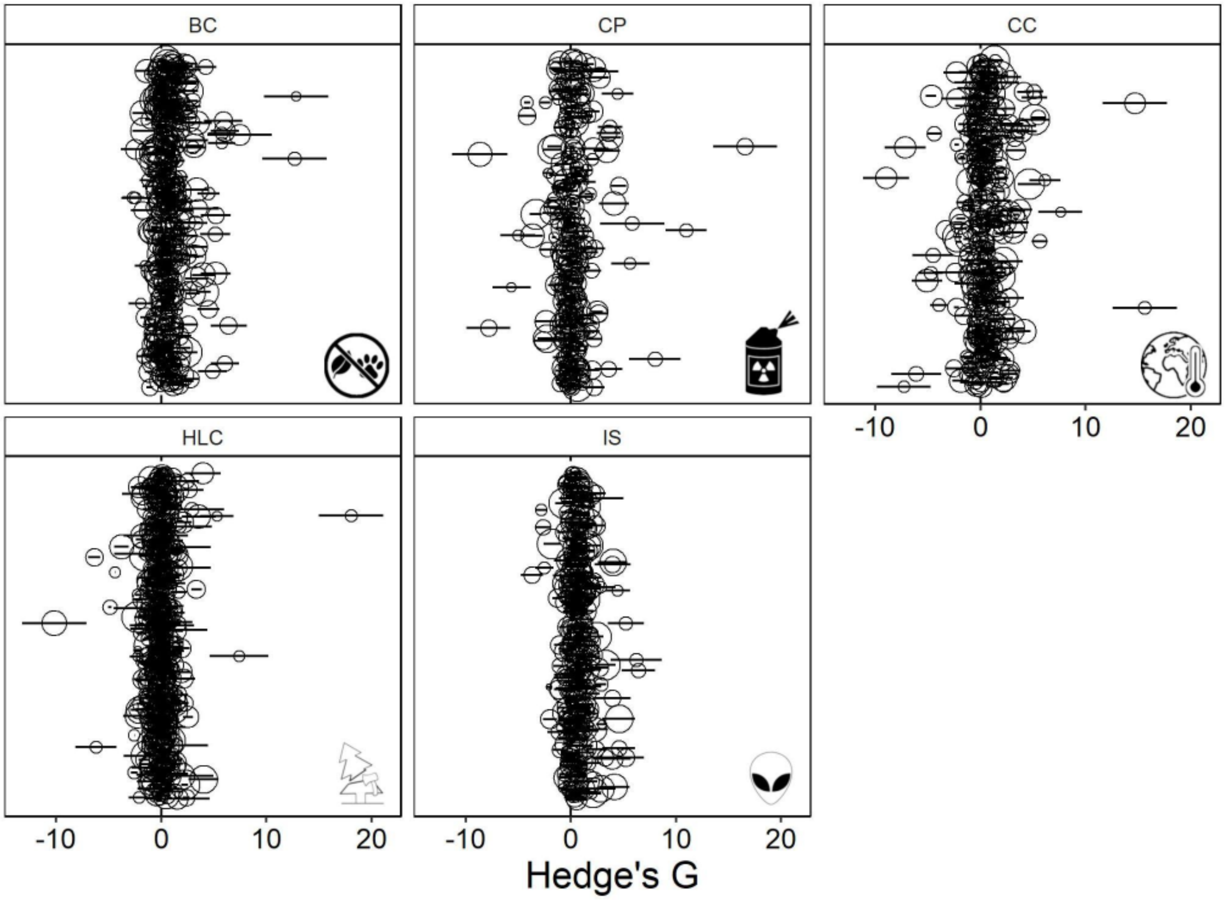


**Fig. S7. Forest plot of effect sizes, associated variances, and relative weights in multilevel metaanalysis models of observations in database.** Error bars are standard errors of the effect size and size of the points is the relative weight of the observation in the model, with larger points representing observations with higher weight in the model. Effect sizes were plotted in a random order.

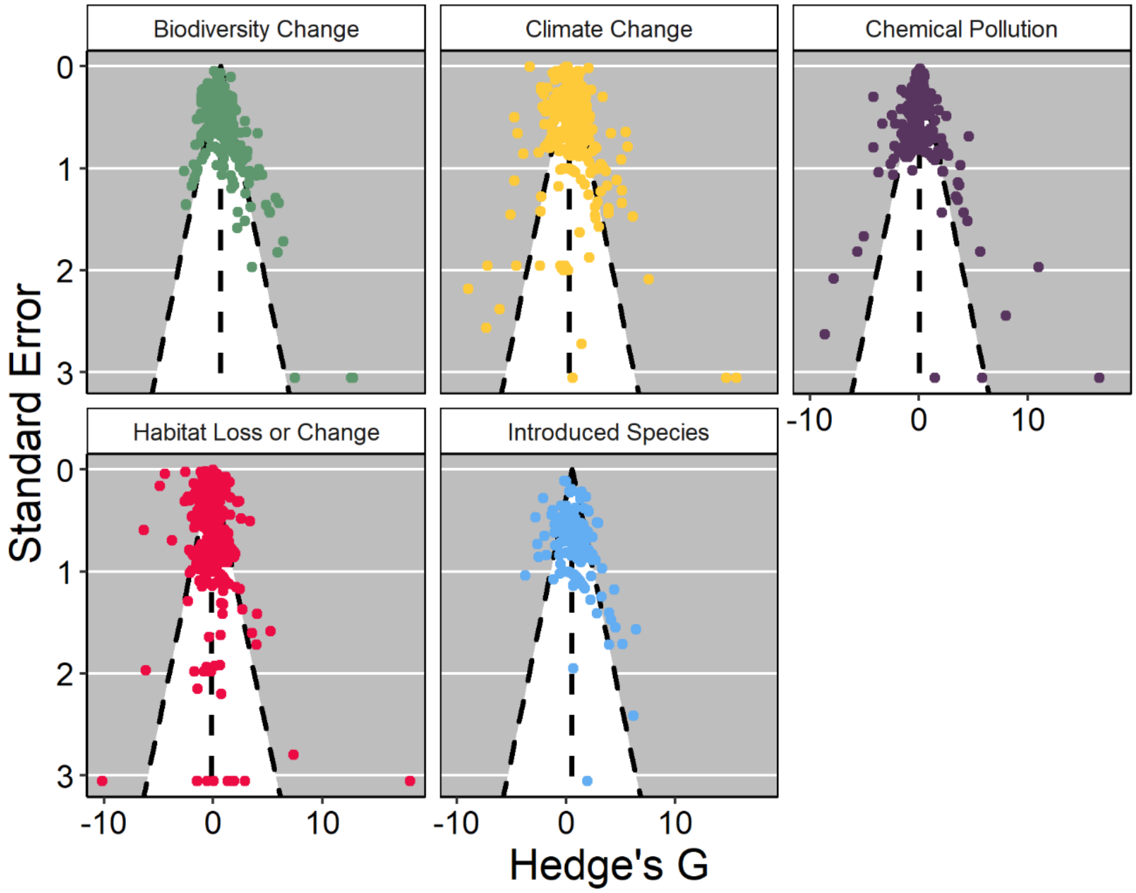


**Fig. S8. Funnel plots for each of the five global change drivers.** Egger's tests indicated no significant asymmetry suggesting negligible publication bias in reported disease responses across the five global change drivers.

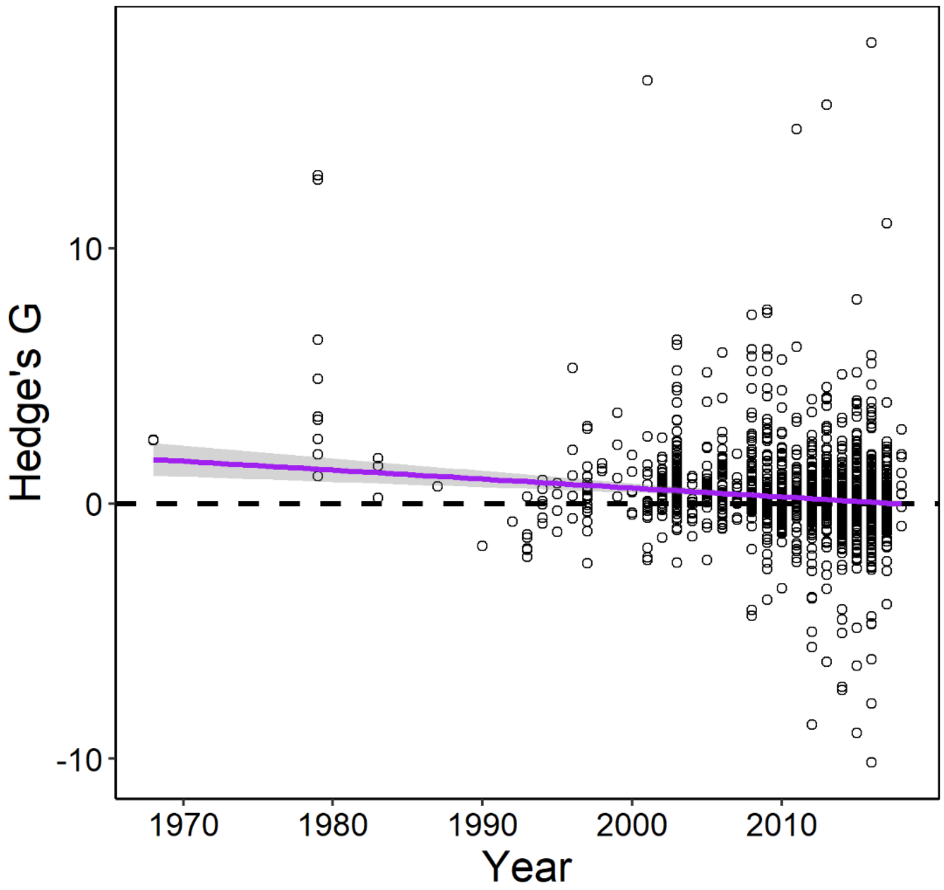


**Fig. S9. Scatter plot showing potential time-lag bias.** Meta-regression of year published as the only moderator indicated a decline effect through time ($p < 0.001$), suggesting a minor time lag bias may be present. However, time lag bias is not necessarily indicative of an overall reduction in published strength of effect; rather, the relationship is likely driven by first 20 years of dataset being solely positive effects. Further, the relationship is weak ($\beta_{Time}$= -0.03), which by the end of the dataset suggests an effect size that is not different from zero, so the bias is towards strong positive effects early on in dataset, during which there are relatively few observations (96% of the dataset was published since 2000). A subsequent re-analysis using only data from 2000 onwards indicated no time-lag bias. Thus, while statistically significant, the time-lag bias is of minor concern.

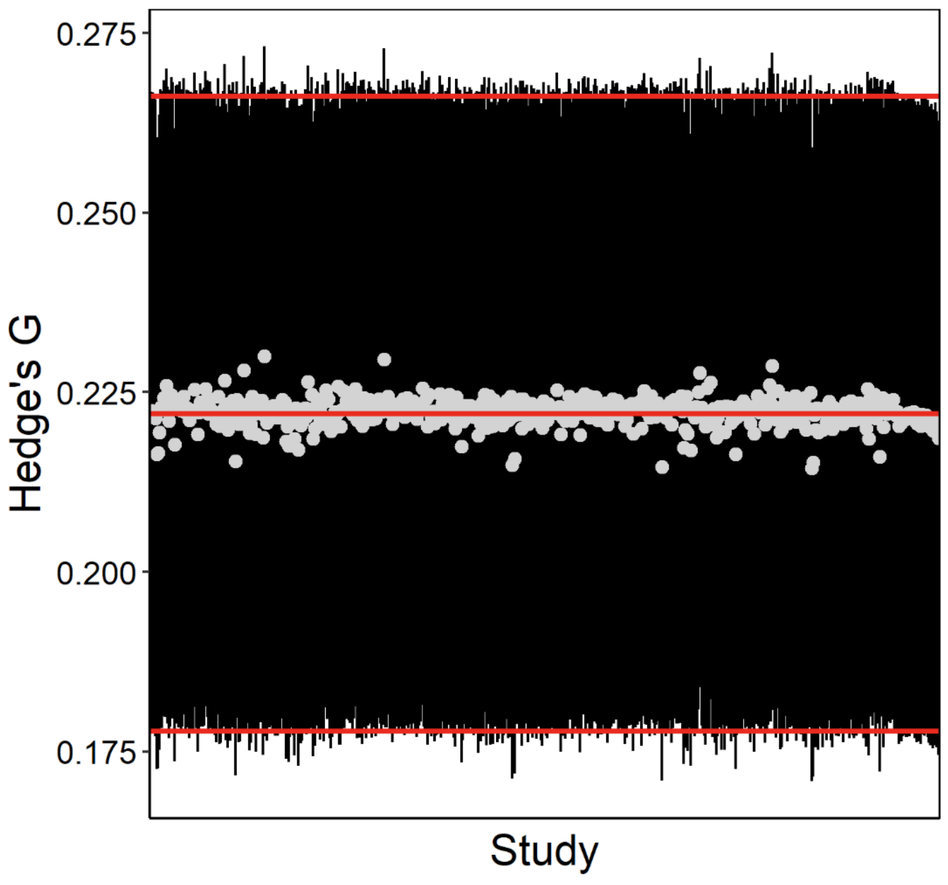


**Figure S10. Results of leave-one-out sensitivity analysis.** The horizontal red lines denote the grand mean and SE of Hedge's g and (g = 0.2087, SE = 0.0438). Grey points and errorbars indicate the Hedge's g and SEs, respectively, using the leave-one-out method (grand mean is recalculated after a given study is removed from dataset). While the remove of certain studies resulted in values that differed from the grand mean, all estimated Hedge's g values fell well within the SE of the grand mean. This sensitivity analysis indicates that our results were robust to the iterative exclusion of individual studies.

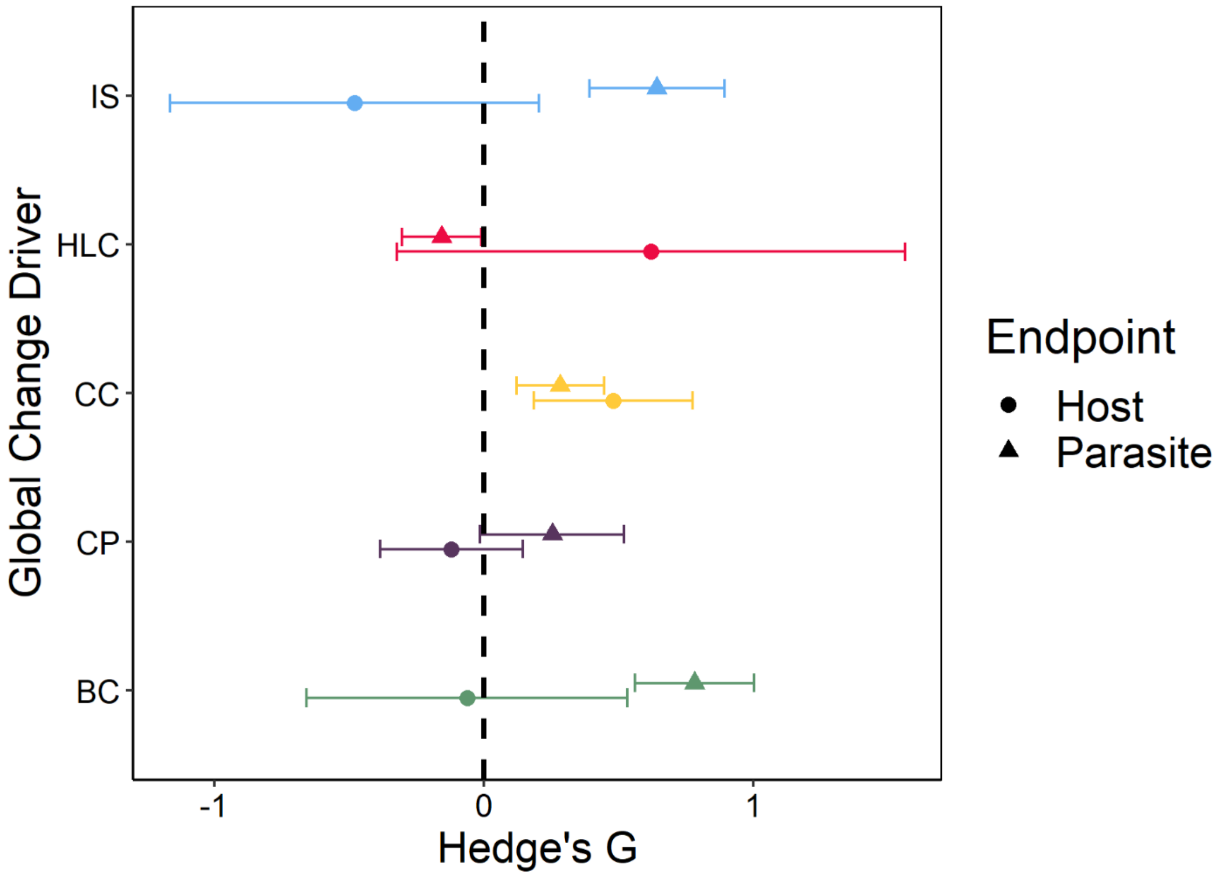


**Fig. S11. The effects of five common global change drivers on mean infectious disease responses in the literature occasionally depend on whether the study endpoint was the host or parasite.** Parasite endpoints responded more positively than host endpoints to both biodiversity change (BC; $Z$=-2.605, $p$=0.0092), chemical pollution (CP; Z = -2.311, p = 0.0208), and introduced species (IS; $Z$=-3.077, $p$=0.0021). There were no significant differences between host and parasite endpoints ($p$>0.05) for climate change (CC) or habitat loss and change (HLC). The displayed points represent the mean predicted values (with 95% confidence intervals) from a *metafor* model where the response variable was a Hedge's G (representing the effect on an infectious disease endpoint relative to control), study was treated as a random effect, and the independent variables included the main effects and an interaction between global change driver and study endpoint. Host endpoints generally included host disease, survival, growth, and reproduction, whereas parasite endpoints generally included parasite prevalence, incidence, abundance, survival, growth, and richness.

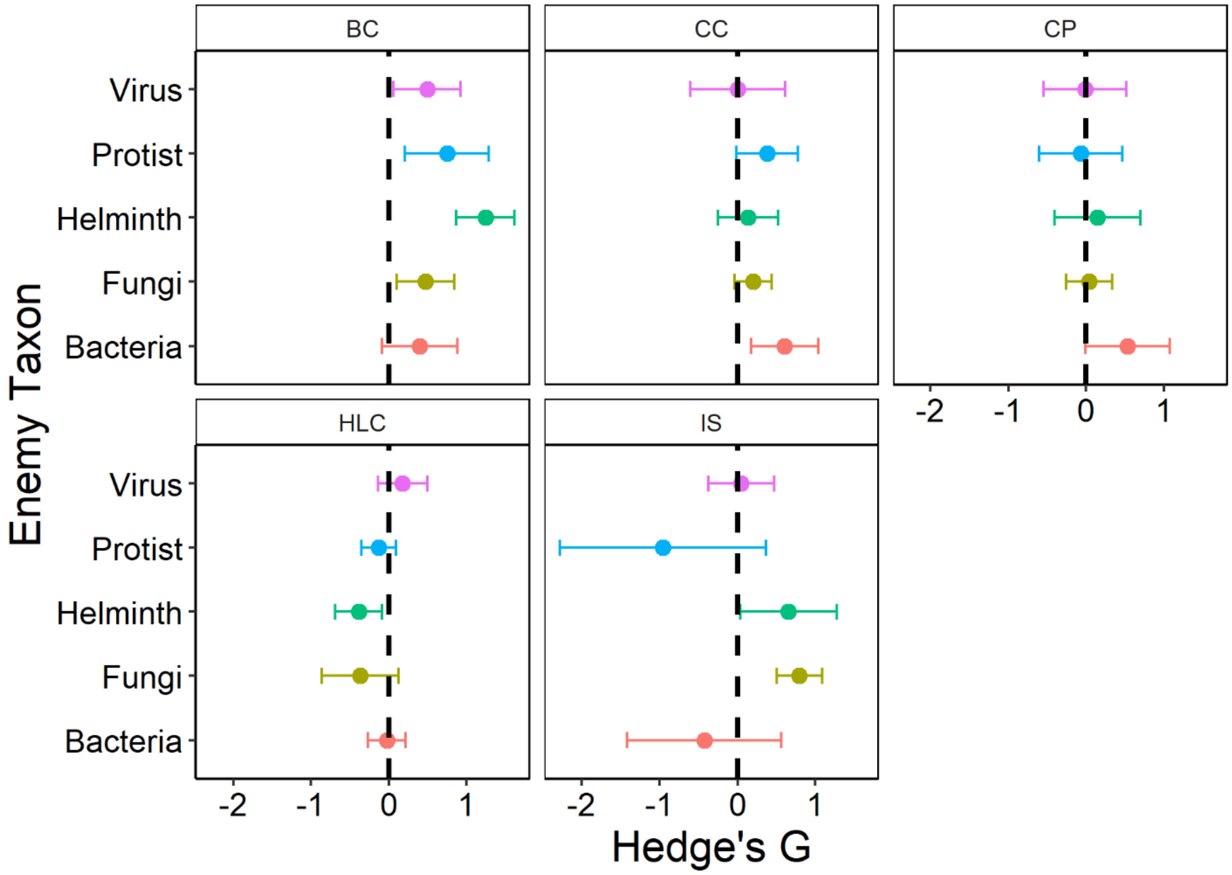


**Fig. S12. The effects of five common global change drivers on mean infectious disease responses in the literature only occasionally depend on parasite taxon.** Helminths responded more positively to biodiversity loss (BC) than all other parasite taxa ($Z$>2.847, $p$<0.0357) except protists ($Z$=1.752, $p$>0.05). Fungi responded more positively to introduced species (IS) than did viruses ($Z$=3.883, $p$=0.0010). There was considerable overlap among the 95% confidence intervals for each parasite taxon for the effects of chemical pollution (CP) and climate change (CC). The displayed points represent the mean predicted values (with 95% confidence intervals) from a *metafor* model where the response variable was a Hedge's G (representing the effect on an infectious disease endpoint relative to control), study was treated as a random effect, and the independent variables included the main effects and an interaction between global change driver and parasite taxon.

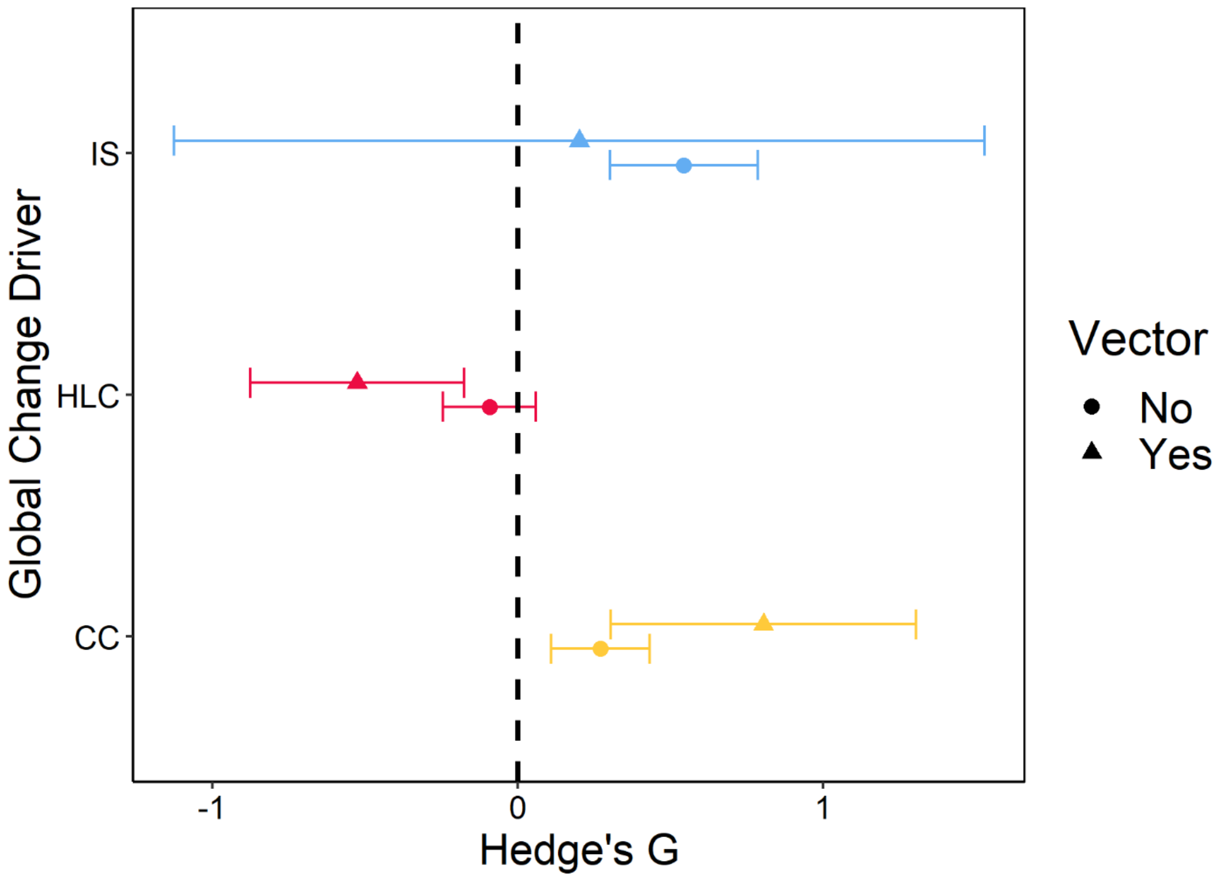


**Fig. S13. The effects of common global change drivers on mean infectious disease responses in the literature depends on whether the parasite is a vector or not.** Vector parasites responded more positively and negatively than non-vector parasites to climate change (CC; $z = 2.013$, $p < 0.05$) and habitat loss and change (HLC; $z = -2.398$, $p < 0.05$); respectively. There was no significant difference for introduced species (IS; $p > 0.10$). The displayed points represent the mean predicted values (with 95% confidence intervals) from a *metafor* model where the response variable was a Hedge's G (representing the effect on an infectious disease endpoint relative to control), study was treated as a random effect, and the independent variables included the main effects and an interaction between global change driver and whether the parasite is a vector or not. Biodiversity change (BC) and chemical pollution (CP) were not included in this set of analyses, because of poor replication for these drivers (2 observations of vectors in CP and no observations of vectors in BC).

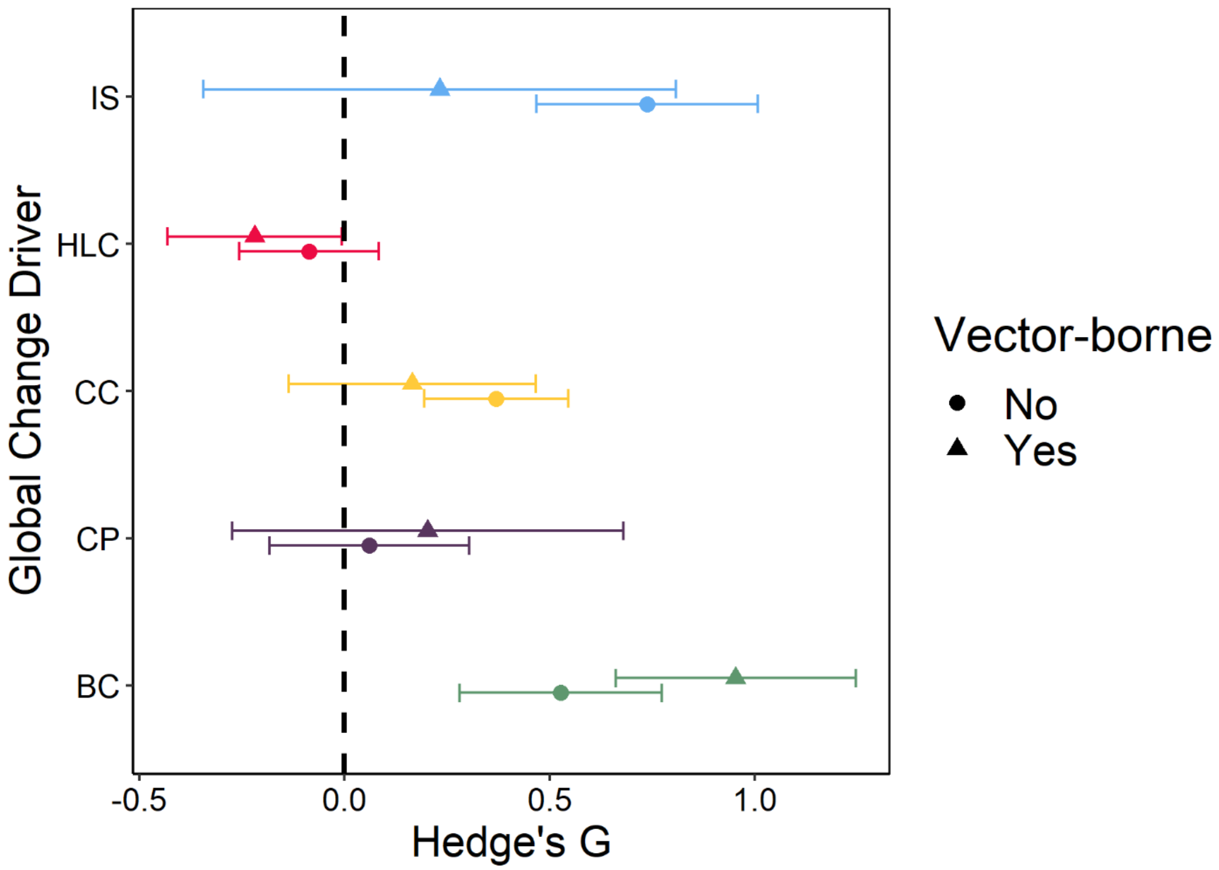


**Fig. S14. The effects of five common global change drivers on mean infectious disease responses in the literature depends on whether the parasite is vector-borne.** Vector-borne parasites responded more positively than non-vector-borne parasites to biodiversity loss (BC; $Z$=2.510, $p$=0.0121). There were no significant differences ($p$>0.10) for climate change (CC), chemical pollution (CP), habitat loss and change (HLC), or introduced species (IS). The displayed points represent the mean predicted values (with 95% confidence intervals) from a *metafor* model where the response variable was a Hedge's G (representing the effect on an infectious disease endpoint relative to control), study was treated as a random effect, and the independent variables included the main effects and an interaction between global change driver and whether the parasite is vector-borne.

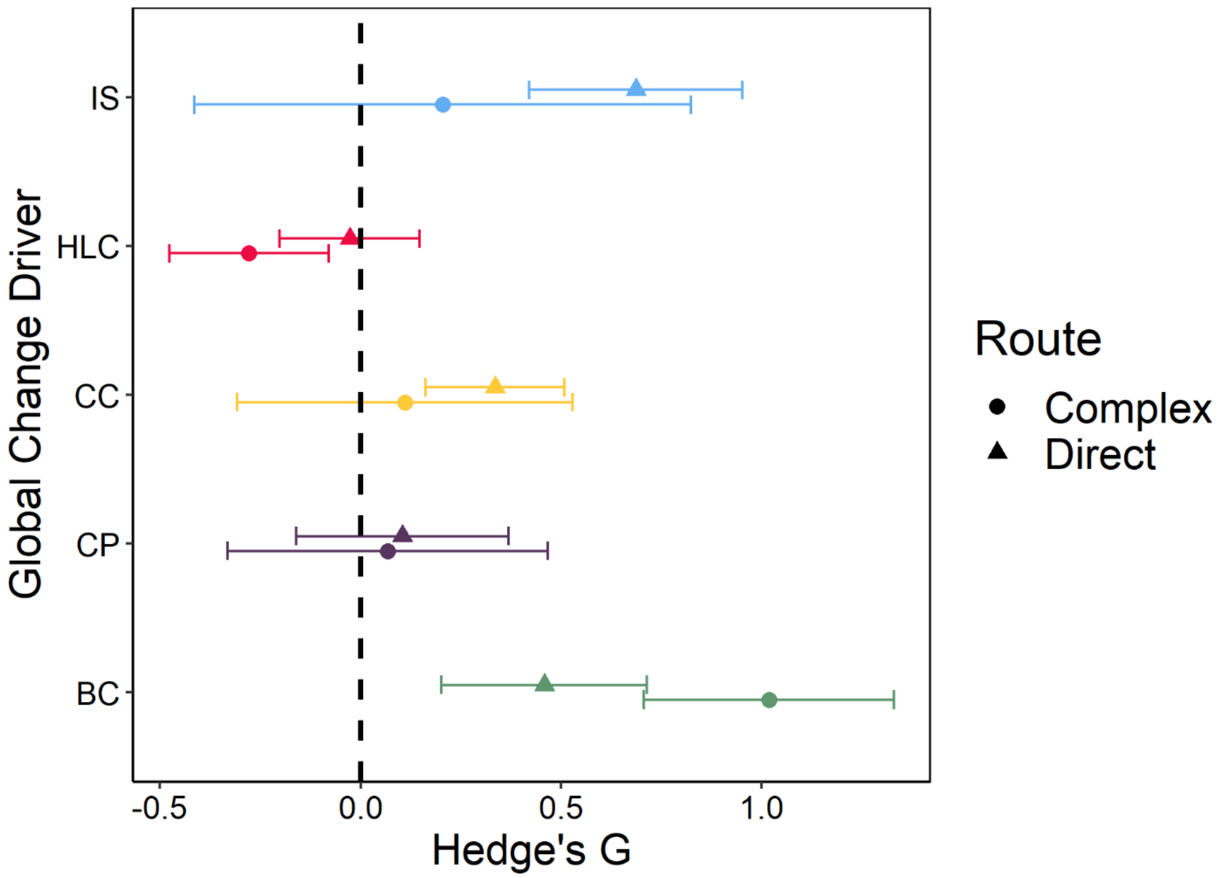


**Fig. S15. The effects of five common global change drivers on mean infectious disease responses in the literature occasionally depend on whether the parasite has a complex or direct life cycle.** Parasites with complex life cycles responded more positively and negatively than parasites with direct life cycles to biodiversity loss (BC; $Z$=2.930, $p$=0.0034) and habitat loss and change (HLC; $Z$=-2.241, $p$=0.025); respectively. There were no significant differences ($p$>0.05) for climate change (CC), chemical pollution (CP), or introduced species (IS). The displayed points represent the mean predicted values (with 95% confidence intervals) from a *metafor* model where the response variable was a Hedge's G (representing the effect on an infectious disease endpoint relative to control), study was treated as a random effect, and the independent variables included the main effects and an interaction between global change driver and type of parasite life cycle.

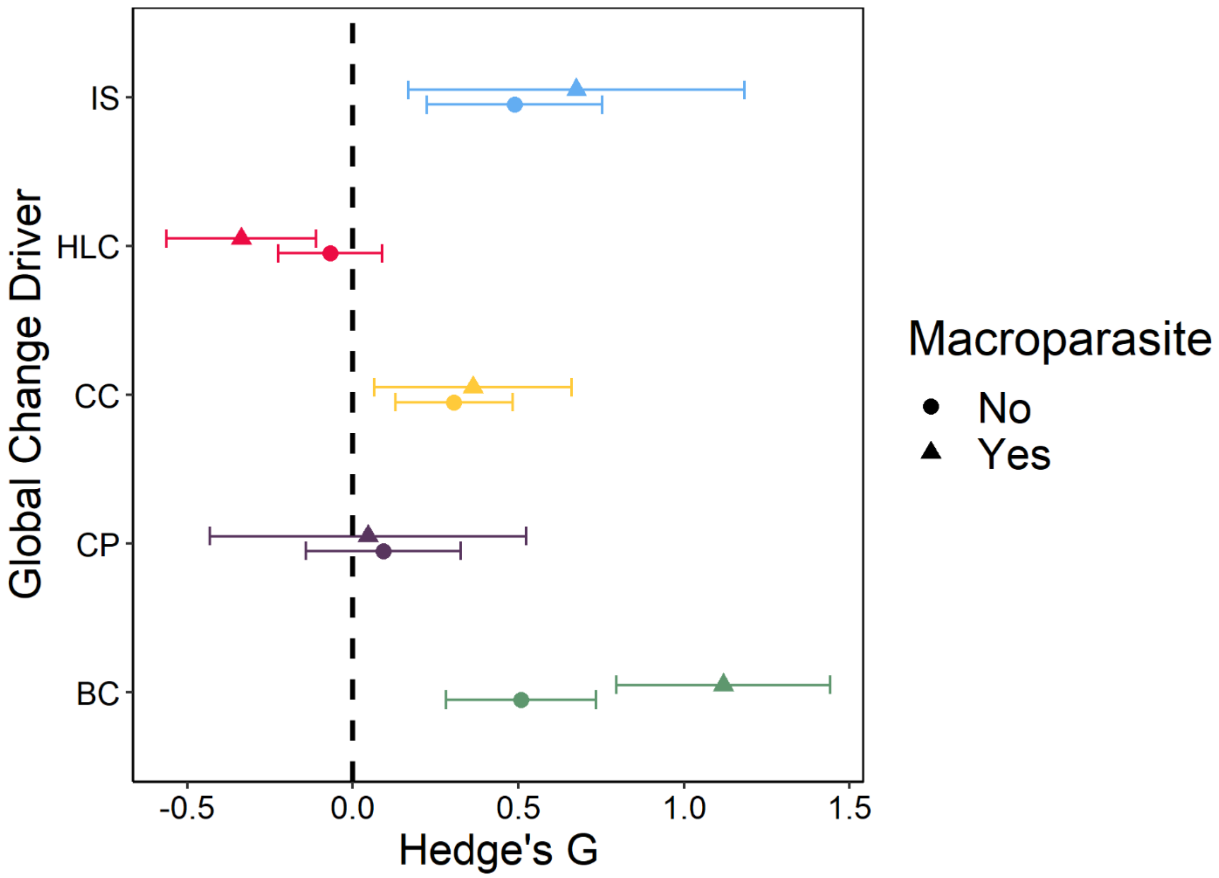


**Fig. S16. The effects of five common global change drivers on mean infectious disease responses in the literature depends on whether the parasite is a micro- or macroparasite.** Macroparasites responded more positively and negatively than microparasites to biodiversity loss (BC; $Z$=3.485, $p$=0.0005) and to habitat loss and change (HLC; $Z$=2.254, $p$=0.0242). There were no significant differences ($p$>0.05) for climate change (CC), chemical pollution (CP), habitat loss and change (HLC), or introduced species (IS). The displayed points represent the mean predicted values (with 95% confidence intervals) from a *metafor* model where the response variable was a Hedge's G (representing the effect on an infectious disease endpoint relative to control), study was treated as a random effect, and the independent variables included the main effects and an interaction between global change driver and whether the parasite is a micro- or macroparasite.

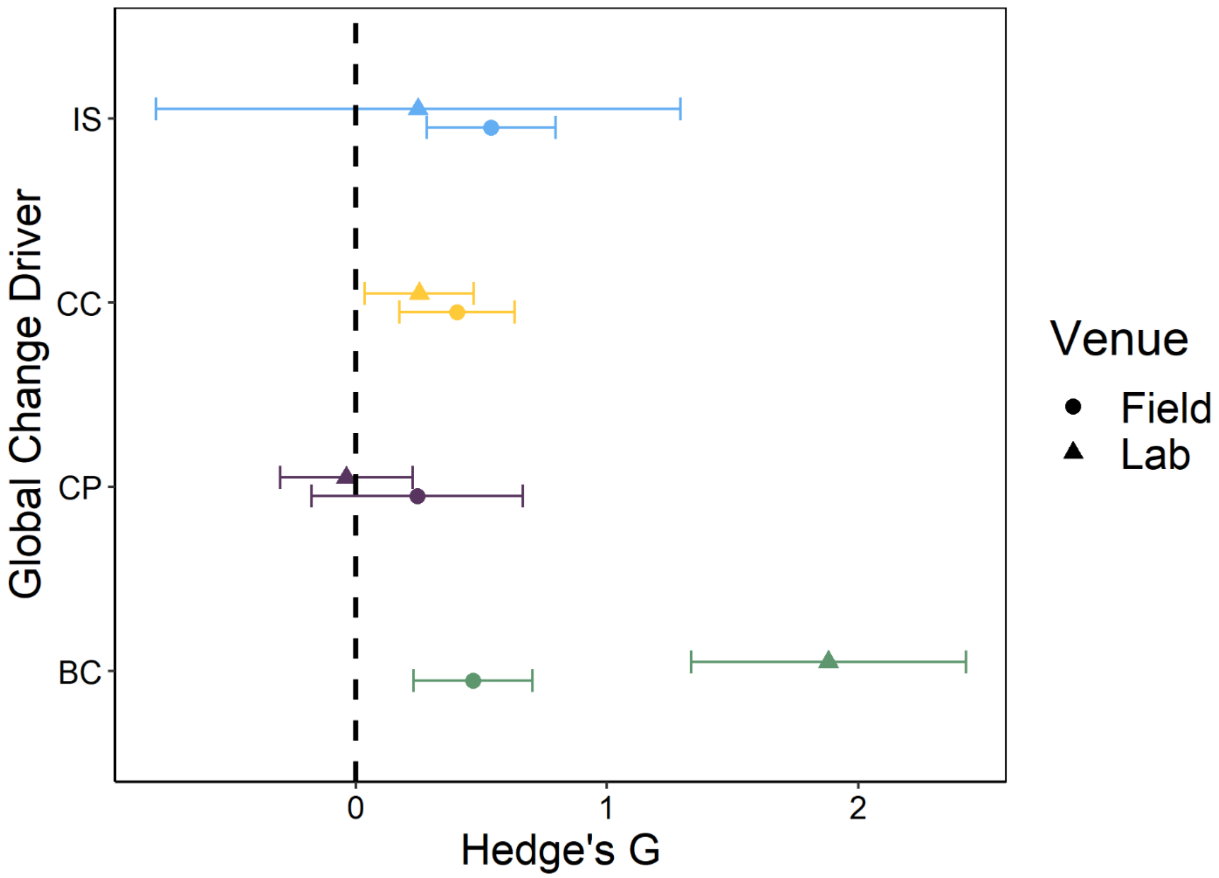


**Fig. S17. The effects of five common global change drivers on mean infectious disease responses in the literature occasionally depend on whether the study was conducted in the laboratory or the field.** Laboratory studies detect significantly stronger positive effects of biodiversity change (BC) on infectious disease endpoints than field studies ($Z$=4.660, $p$<0.0001), whereas chemical pollution (CP), climate change (CC), and introduced species (IS) show stronger positive responses in the field than laboratory, although all were non-significant ($p$>0.05). The habitat loss and change global change driver is not displayed because all of the studies were conducted in the field. The displayed points represent the mean predicted values (with 95% confidence intervals) from a *metafor* model where the response variable was a Hedge's G (representing the effect on an infectious disease endpoint relative to control), study was treated as a random effect, and the independent variables included the main effects and an interaction between global change driver and study venue.

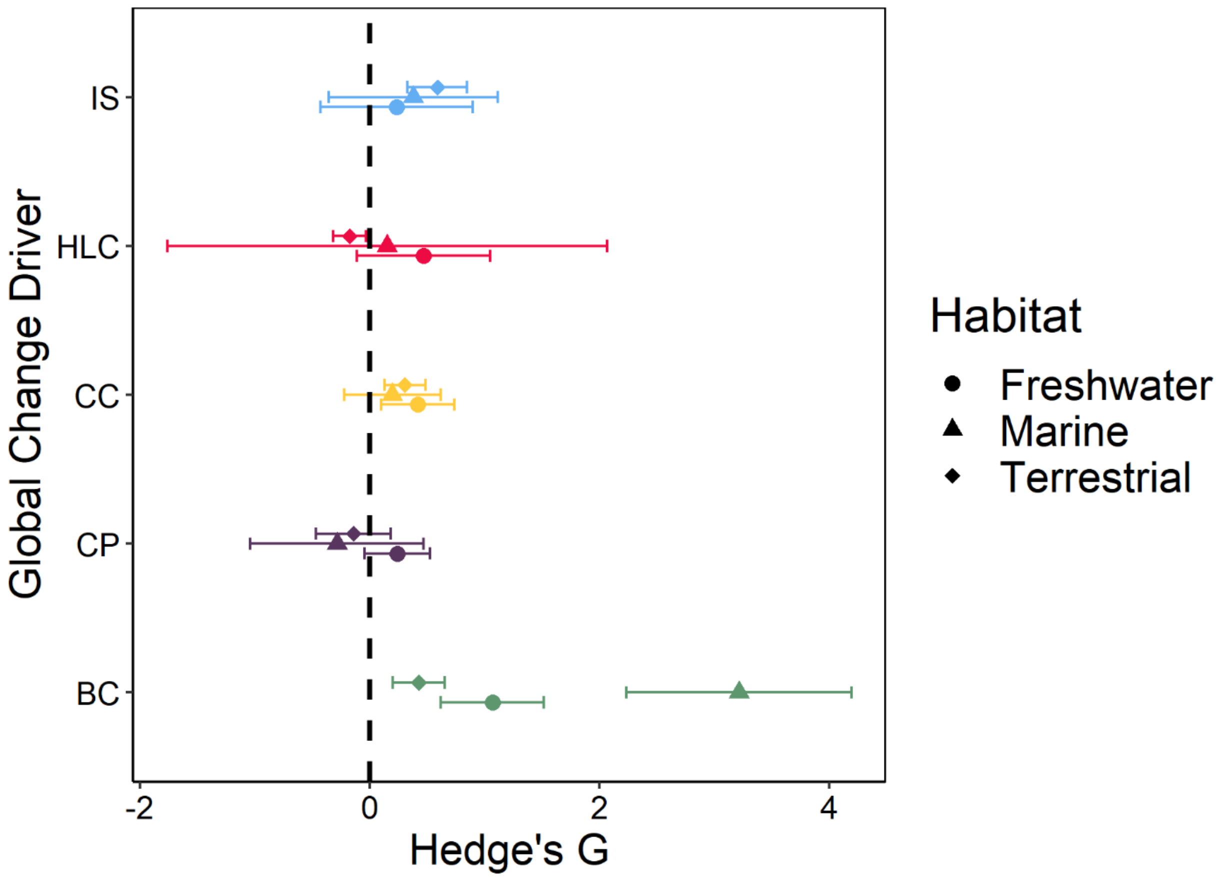


**Fig. S18. Infectious disease endpoints vary across habitat types in biodiversity change studies but not in studies on other global change drivers.** The three habitat types differed from one another for biodiversity change (BC, $Z>2.495$, $p<0.0337$), whereas there were no significant differences ($p>0.05$) among the habitat types for climate change (CC), chemical pollution (CP), habitat loss and change (HLC), or introduced species (IS). The displayed points represent the mean predicted values (with 95% confidence intervals) from a *metafor* model where the response variable was a Hedge's G (representing the effect on an infectious disease endpoint relative to control), study was treated as a random effect, and the independent variables included the main effects and an interaction between global change driver and habitat.

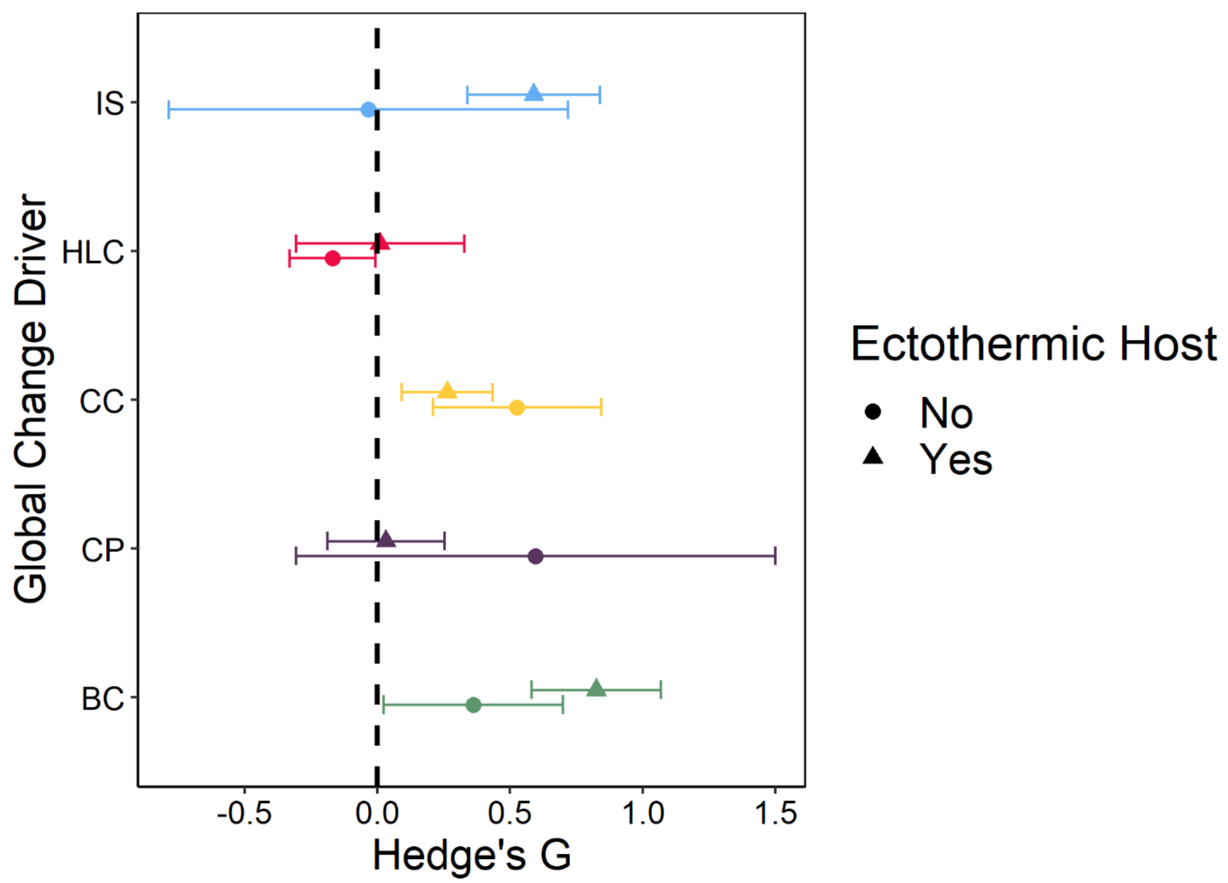


**Fig. S19. The effects of five common global change drivers on mean infectious disease responses in the literature depend on whether the host is ecto- or endothermic.** Ectothermic hosts show stronger positive responses than endothermic hosts to biodiversity change (CC; $Z$=2.289, $p$=0.0221) There were no significant differences ($p$>0.05) for climate change (CC), chemical pollution (CP), habitat loss and change (HLC) or introduced species (IS). The displayed points represent the mean predicted values (with 95% confidence intervals) from a *metafor* model where the response variable was a Hedge's G (representing the effect on an infectious disease endpoint relative to control), study was treated as a random effect, and the independent variables included the main effects and an interaction between global change driver and whether the host was ecto- or endothermic.

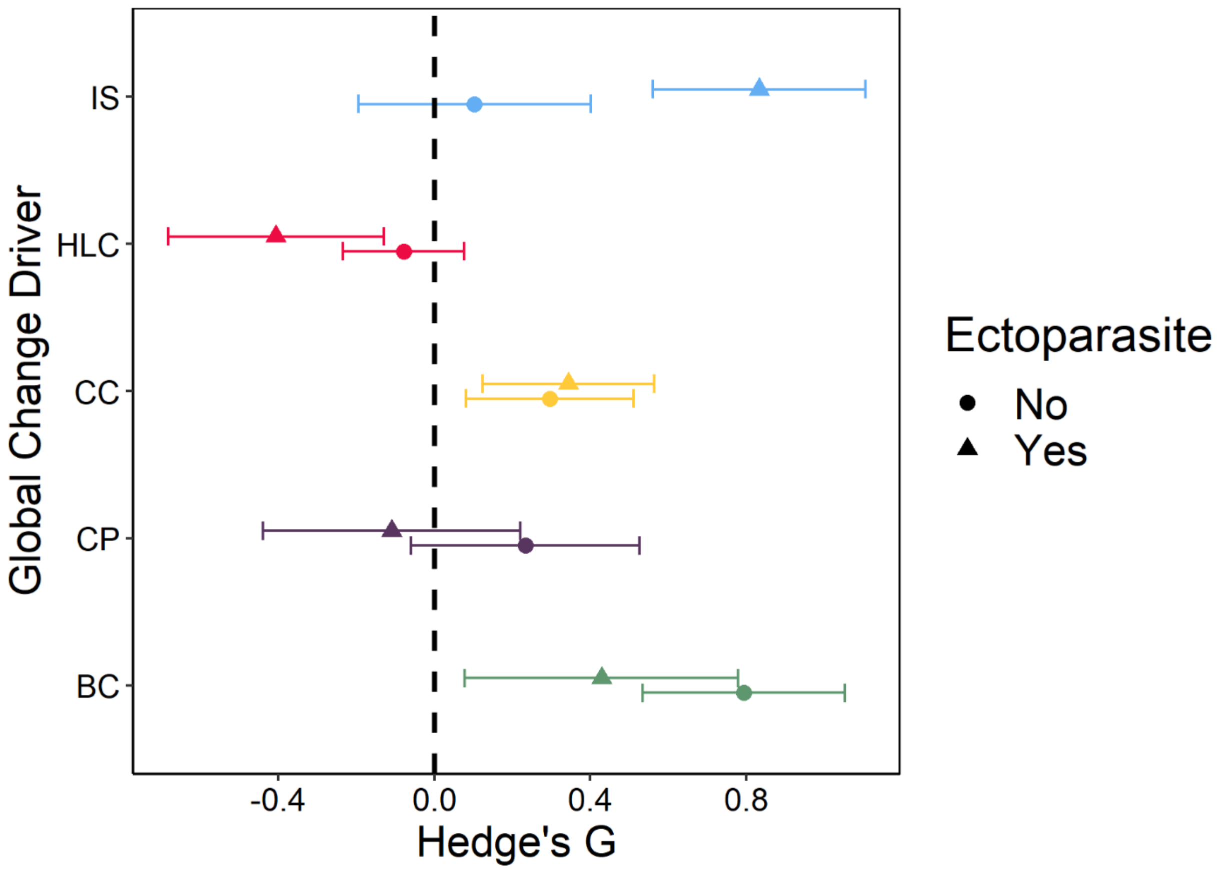


**Fig. S20. The effects of five common global change drivers on mean infectious disease responses in the literature depend on whether the parasite in ecto- or endoparasitic.** Ectoparasites show stronger positive and negative responses than endoparasites to introduced species (IS; $Z$=4.683, $p$<0.0001) and habitat loss and change (HLC; $Z$=-2.259, $p$=0.0239), respectively. Virtually all fungi in the database are ectoparasites, which is likely driving the patterns for IS. There were no significant differences ($p$>0.05) for climate change (CC), chemical pollution (CP), or introduced species (IS). The displayed points represent the mean predicted values (with 95% confidence intervals) from a *metafor* model where the response variable was a Hedge's G (representing the effect on an infectious disease endpoint relative to control), study was treated as a random effect, and the independent variables included the main effects and an interaction between global change driver and whether the parasite was ecto- or endoparasitic.

**Table S1.** Literature search dates, search terms, and refinements used to generate the database on the five global change dirvers and infectious disease. See Methods for details on inclusion criteria and data extracted from each study.

| Global change driver | Date of Web of Science Search | Search terms | Refinements |
|---|---|---|---|
| Biodiversity change | 2018 (see details in Methods) | Civitello et al. (2015), Halliday and Rohr (2019), and Magnusson et al. (2020): used a combination of parasite, pathogen, diversity, richness, evenness, dilution effect, and decoy effect, and identified additional papers by searching the literature cited sections of these and related articles; Liu et al. (2020): [(biodivers* OR divers* OR mixed specie* OR mixed-specie* OR species divers* OR species rich* OR polyculture* OR diversity-disease*) AND (disease* OR pathogen* OR infect* OR epidemic*) AND (plant* OR tree* OR forest* OR wood* OR grass* OR herb*) AND (inciden* OR prevalen* OR load* OR severity OR occur* OR abundance)], identified additional papers by examining the references of previous meta-analyses and literature reviews, and also searched the China National Knowledge Infrastructure (<www.cnki.net>) with the terms (diversity AND plant AND disease) for Chinese peer-reviewed studies | None |
| Climate change | Sept. 1, 2017 | "climate change" AND disease AND (parasit* OR pathogen) | None |
| Chemical pollution | Sept. 6, 2017 | disease, parasite, pathogen, metal, metalloid, organometal, nitrogen oxide, sulfur oxide, acid rain, nitrogen fertilizer, phosphorous fertilizer, fertiliz*, sulfate effluent, sulfuric acid, acid mine drainage, acid rock drainage, hydrochlorofluorocarbon, chlorofluorocarbon, organochlorine alkene, degreas*, dense nonaqueous phase liquid, polycyclic aromatic hydrocarbon, polynuclear aromatic hydrocarbon, polyhalogenated benzene, polyhalogenated phenol, polyhalogenated biphenyl, polychlorinated benzene, polychlorinated phenol, polychlorinated biphenyl, polychlorinated terphenyl, polybrominated biphenyl, polychlorinated naphthalene, polychlorinated dibenzodioxin, polychlorinated dibenzofuran, pesticide, herbicide, insecticide, fungicide | None |
| Habitat loss and change | Oct. 8, 2017 | disease* AND (parasit* OR pathoge*) AND (defores* OR urban* OR "habitat loss" OR "habitat fragmentation") | excluding the following Web of Science categories: (CRITICAL CARE MEDICINE OR CHEMISTRY MEDICINAL OR PLANNING DEVELOPMENT OR HEALTH POLICY SERVICES OR SOCIAL SCIENCES BIOMEDICAL OR METEOROLOGY ATMOSPHERIC SCIENCES OR HEMATOLOGY OR OBSTETRICS GYNECOLOGY OR ECONOMICS OR CONSTRUCTION BUILDING TECHNOLOGY OR CHEMISTRY MULTIDISCIPLINARY OR SOCIAL SCIENCES INTERDISCIPLINARY OR HEALTH CARE SCIENCES SERVICES OR REPRODUCTIVE BIOLOGY OR OTORHINOLARYNGOLOGY OR EMERGENCY MEDICINE OR NURSING OR PSYCHIATRY OR INTERNATIONAL RELATIONS OR HISTORY OR ONCOLOGY OR DENTISTRY ORAL SURGERY MEDICINE OR IMAGING SCIENCE PHOTOGRAPHIC TECHNOLOGY OR HISTORY PHILOSOPHY OF SCIENCE OR OPHTHALMOLOGY OR ENGINEERING BIOMEDICAL OR NEUROSCIENCES OR COMPUTER SCIENCE THEORY METHODS OR COMPUTER SCIENCE INTERDISCIPLINARY APPLICATIONS OR BIOCHEMICAL RESEARCH METHODS OR COMPUTER SCIENCE INFORMATION SYSTEMS OR CHEMISTRY ANALYTICAL OR SOCIOLOGY OR BIOPHYSICS), and the following Web of Science document types: (REVIEW OR EDITORIAL MATERIAL OR RETRACTED PUBLICATION OR MEETING ABSTRACT). |
| Introduced species | Sept. 8, 2017 | disease* AND (parasit* OR pathoge*) AND ("invasive species" OR "introduced species" OR "non-native species" OR "enemy release" OR "enemy-release" OR "spillover" OR "spillback") | excluding the following Web of Science categories: (MATHEMATICS INTERDISCIPLINARY APPLICATIONS OR IMAGING SCIENCE PHOTOGRAPHIC TECHNOLOGY OR GASTROENTEROLOGY HEPATOLOGY OR COMPUTER SCIENCE INTERDISCIPLINARY APPLICATIONS OR CELL BIOLOGY OR UROLOGY NEPHROLOGY OR URBAN STUDIES OR PERIPHERAL VASCULAR DISEASE OR SOCIAL SCIENCES MATHEMATICAL METHODS OR SOCIAL SCIENCES BIOMEDICAL OR ENDOCRINOLOGY METABOLISM OR PHYSICS FLUIDS PLASMAS OR PATHOLOGY OR PHARMACOLOGY PHARMACY OR OPERATIONS RESEARCH MANAGEMENT SCIENCE OR MEDICINE GENERAL INTERNAL OR ONCOLOGY OR METEOROLOGY ATMOSPHERIC SCIENCES OR MATHEMATICS OR ECONOMICS OR INTEGRATIVE COMPLEMENTARY MEDICINE OR HEMATOLOGY OR HEALTH CARE SCIENCES SERVICES OR GREEN SUSTAINABLE SCIENCE TECHNOLOGY OR EDUCATION SCIENTIFIC DISCIPLINES OR ENGINEERING ENVIRONMENTAL OR CRITICAL CARE MEDICINE OR CARDIAC CARDIOVASCULAR SYSTEMS OR CHEMISTRY MULTIDISCIPLINARY OR BIOPHYSICS OR BIOCHEMICAL RESEARCH METHODS OR PHYSICS MATHEMATICAL OR BEHAVIORAL SCIENCES OR NUTRITION DIETETICS OR AGRICULTURE DAIRY ANIMAL SCIENCE), and the following Web of Science document types: (REVIEW OR EDITORIAL MATERIAL) |

**Table S2. Summary of the number of studies and effect sizes in the infectious disease database across ecological contexts and global change drivers.**

| Study, Host, and Parasite Factors | | Global Change Driver | Number of Studies | Number of Effect Sizes |
|---|---|---|---|---|
| Parasite Taxa | Arthropod | BC | 9 | 71 |
| | | CC | 23 | 36 |
| | | CP | 1 | 1 |
| | | HLC | 26 | 78 |
| | | IS | 6 | 10 |
| | Bacteria | BC | 12 | 26 |
| | | CC | 23 | 37 |
| | | CP | 10 | 24 |
| | | HLC | 56 | 126 |
| | | IS | 2 | 4 |
| | Fungi | BC | 21 | 172 |
| | | CC | 69 | 138 |
| | | CP | 39 | 127 |
| | | HLC | 11 | 22 |
| | | IS | 46 | 198 |
| | Helminth | BC | 17 | 86 |
| | | CC | 29 | 59 |
| | | CP | 10 | 59 |
| | | HLC | 27 | 88 |
| | | IS | 10 | 44 |
| | Protist | BC | 5 | 14 |
| | | CC | 27 | 47 |
| | | CP | 11 | 23 |
| | | HLC | 68 | 109 |
| | | IS | 2 | 3 |
| | Virus | BC | 19 | 37 |
| | | CC | 12 | 19 |
| | | CP | 10 | 22 |
| | | HLC | 28 | 73 |
| | | IS | 21 | 39 |
| Native Host Taxa | Amphibian/Reptile | BC | 5 | 26 |
| | Amphibian/Reptile | CC | 16 | 29 |
| | Amphibian/Reptile | CP | 21 | 96 |
| | Amphibian/Reptile | HLC | 6 | 12 |
| | Amphibian/Reptile | IS | 4 | 5 |
| | Arthropod | BC | 4 | 6 |
| | Arthropod | CC | 8 | 14 |
| | Arthropod | CP | 15 | 38 |
| | Arthropod | HLC | 16 | 55 |
| | Arthropod | IS | 7 | 10 |

| | | | | |
|---|---|---|---|---|
| | Bird | BC | 2 | 2 |
| | Bird | CC | 5 | 11 |
| | Bird | CP | 1 | 2 |
| | Bird | HLC | 20 | 31 |
| | Bird | IS | 4 | 6 |
| | Fish | BC | 1 | 2 |
| | Fish | CC | 9 | 17 |
| | Fish | CP | 5 | 13 |
| | Fish | HLC | 1 | 1 |
| | Fish | IS | 4 | 10 |
| | Mammal | BC | 28 | 88 |
| | Mammal | CC | 36 | 74 |
| | Mammal | CP | 3 | 13 |
| | Mammal | HLC | 127 | 360 |
| | Mammal | IS | 4 | 5 |
| | Mollusk | BC | 11 | 55 |
| | Mollusk | CC | 14 | 20 |
| | Mollusk | CP | 12 | 44 |
| | Mollusk | HLC | 1 | 1 |
| | Mollusk | IS | 5 | 35 |
| | Plant | BC | 29 | 229 |
| | Plant | CC | 64 | 136 |
| | Plant | CP | 19 | 64 |
| | Plant | HLC | 3 | 8 |
| | Plant | IS | 42 | 231 |
| Human parasite? | No | BC | 50 | 329 |
| | No | CC | 159 | 313 |
| | No | CP | 73 | 264 |
| | No | HLC | 48 | 129 |
| | No | IS | 69 | 301 |
| | Yes | BC | 28 | 79 |
| | Yes | CC | 20 | 33 |
| | Yes | CP | 2 | 8 |
| | Yes | HLC | 134 | 331 |
| | Yes | IS | 1 | 1 |
| Host Thermy | Endotherm | BC | 30 | 90 |
| | Endotherm | CC | 41 | 85 |
| | Endotherm | CP | 4 | 15 |
| | Endotherm | HLC | 145 | 388 |
| | Endotherm | IS | 8 | 11 |
| | Ectotherm | BC | 50 | 318 |
| | Ectotherm | CC | 138 | 261 |
| | Ectotherm | CP | 71 | 257 |
| | Ectotherm | HLC | 38 | 116 |

| | | | | |
|---|---|---|---|---|
| | Ectotherm | IS | 62 | 291 |
| Parasite Size | Microparasite | BC | 59 | 279 |
| | Microparasite | CC | 126 | 243 |
| | Microparasite | CP | 63 | 196 |
| | Microparasite | HLC | 149 | 339 |
| | Microparasite | IS | 56 | 250 |
| | Macroparasite | BC | 21 | 129 |
| | Macroparasite | CC | 51 | 93 |
| | Macroparasite | CP | 13 | 64 |
| | Macroparasite | HLC | 49 | 164 |
| | Macroparasite | IS | 17 | 52 |
| Free Living Stages? | No | BC | 17 | 48 |
| | No | CC | 22 | 49 |
| | No | CP | 7 | 13 |
| | No | HLC | 72 | 126 |
| | No | IS | 7 | 9 |
| | Yes | BC | 63 | 359 |
| | Yes | CC | 152 | 283 |
| | Yes | CP | 64 | 245 |
| | Yes | HLC | 122 | 370 |
| | Yes | IS | 64 | 256 |
| Endpoint | Host | BC | 8 | 58 |
| | Host | CC | 41 | 62 |
| | Host | CP | 44 | 144 |
| | Host | HLC | 5 | 12 |
| | Host | IS | 10 | 11 |
| | Parasite | BC | 70 | 350 |
| | Parasite | CC | 153 | 284 |
| | Parasite | CP | 41 | 128 |
| | Parasite | HLC | 176 | 492 |
| | Parasite | IS | 62 | 291 |
| Transmission Route | Complex | BC | 32 | 131 |
| | Complex | CC | 26 | 55 |
| | Complex | CP | 21 | 102 |
| | Complex | HLC | 86 | 154 |
| | Complex | IS | 10 | 42 |
| | Direct | BC | 48 | 277 |
| | Direct | CC | 139 | 267 |
| | Direct | CP | 52 | 155 |
| | Direct | HLC | 109 | 338 |
| | Direct | IS | 61 | 225 |
| Vector-borne? | No | BC | 47 | 280 |
| | No | CC | 135 | 257 |
| | No | CP | 60 | 191 |

| | | | | |
|---|---|---|---|---|
| | No | HLC | 118 | 355 |
| | No | IS | 59 | 223 |
| | Yes | BC | 32 | 127 |
| | Yes | CC | 42 | 75 |
| | Yes | CP | 14 | 69 |
| | Yes | HLC | 76 | 134 |
| | Yes | IS | 12 | 42 |
| Parasite Type | Endoparasite | BC | 54 | 215 |
| | Endoparasite | CC | 91 | 177 |
| | Endoparasite | CP | 43 | 147 |
| | Endoparasite | HLC | 157 | 407 |
| | Endoparasite | IS | 44 | 102 |
| | Ectoparasite | BC | 24 | 193 |
| | Ectoparasite | CC | 85 | 159 |
| | Ectoparasite | CP | 29 | 111 |
| | Ectoparasite | HLC | 34 | 97 |
| | Ectoparasite | IS | 47 | 200 |

**Data S1. (separate file)**
Caption for Data S1: Data and metadata associated with this publication,

**Data S1. (separate file)**
Caption for Data S2: RMarkdown code and output associated with this publication
