## Supplementary material for "Global change drivers and the risk of infectious disease": Data S2: GCD_Analyses_Final.html

#### 6/23/2022

### Setup R environment

Read in R packages, make a ggplot theme to be used throughout the
code and for all figures.

```
library(metafor)
library(MuMIn)
library(tidyverse)
library(multcomp)
library(emmeans)
library(ggeffects)
library(forcats)
library(stringr)

##Make ggplot theme to use throughout
theme_JR <- function (base_size = 12, base_family = "") 
{
    theme(
    panel.background = element_rect(fill=NA),
    panel.grid = element_blank(),
    panel.border = element_rect(color = "black", fill = NA),
    axis.line = element_line(color="black"),
    legend.title = element_text(size=18,color="black"),
    legend.text = element_text(size=16,color="black"),
    legend.key = element_rect(fill=NA,color=NA),
    axis.text = element_text(size=12,color="black"),
    axis.title = element_text(size=16,color="black"),
    strip.background =element_rect(fill=NA, color = "black")
    )
}
```

#### Grand Mean effect of GCDs on disease

Analysis of grand mean indicates that overall, global change will
increase disease.

```
mgrand <- rma.mv(yi, vi2,
             random = list(~1|Citation.number, ~1|ind_id),
             data = fulldat2)
summary(mgrand)
```

```
## 
## Multivariate Meta-Analysis Model (k = 1832; method: REML)
## 
##     logLik    Deviance         AIC         BIC        AICc 
## -2817.4911   5634.9821   5640.9821   5657.5200   5640.9953   
## 
## Variance Components:
## 
##             estim    sqrt  nlvls  fixed           factor 
## sigma^2.1  0.7568  0.8699    579     no  Citation.number 
## sigma^2.2  0.3172  0.5632   1832     no           ind_id 
## 
## Test for Heterogeneity:
## Q(df = 1831) = 1454471.2057, p-val < .0001
## 
## Model Results:
## 
## estimate      se    zval    pval   ci.lb   ci.ub 
##   0.2222  0.0442  5.0231  <.0001  0.1355  0.3089  *** 
## 
## ---
## Signif. codes:  0 '***' 0.001 '**' 0.01 '*' 0.05 '.' 0.1 ' ' 1
```

```
##I2 (heterogeneity statistic) calculation
W <- diag(1/mgrand$vi)
X <- mgrand$X
P <- W - W %*% X %*% solve(t(X) %*% W %*% X) %*% t(X) %*% W
100 * sum(mgrand$sigma2) / (sum(mgrand$sigma2) + (mgrand$k-mgrand$p)/sum(diag(P)))
```

```
## [1] 99.87599
```

```
##I2 = 0.9987599

##70.37616 is due to between study variation; 29.49983 is due to within study variation
100 * mgrand$sigma2 / (sum(mgrand$sigma2) + (mgrand$k-mgrand$p)/sum(diag(P)))
```

```
## [1] 70.37616 29.49983
```

Funnel plot to investigate publication biases in dataset.

Conduct an Egger’s test to test for asymmetry in funnels plot. It can
identify small-study effects, but not tell directly whether publication
bias exists. To evaluate the funnel asymmetry, we inspect the size of
the intercept when the response is the effect size predictor is either
SE or variance (meta-regression, so weighted by inverse of SE), and if
it differs significantly from zero. When this is the case, Egger’s test
indicates funnel plot asymmetry. Additionally, test for time lag
effect.

```
###Publication bias analysis

##Time lag and Egger's test for multilevel models

##Egger's test
mpbias1 <- rma.mv(yi, vi2, mods = ~ sqrt(vi2),
                 random = list(~1|Citation.number, ~1|ind_id),
                 data = fulldat2)
summary(mpbias1)
```

```
## 
## Multivariate Meta-Analysis Model (k = 1832; method: REML)
## 
##     logLik    Deviance         AIC         BIC        AICc 
## -2764.3830   5528.7661   5536.7661   5558.8144   5536.7880   
## 
## Variance Components:
## 
##             estim    sqrt  nlvls  fixed           factor 
## sigma^2.1  0.6508  0.8068    579     no  Citation.number 
## sigma^2.2  0.3156  0.5618   1832     no           ind_id 
## 
## Test for Residual Heterogeneity:
## QE(df = 1830) = 1451582.9093, p-val < .0001
## 
## Test of Moderators (coefficient 2):
## QM(df = 1) = 105.6184, p-val < .0001
## 
## Model Results:
## 
##            estimate      se     zval    pval    ci.lb    ci.ub 
## intrcpt     -0.2370  0.0611  -3.8792  0.0001  -0.3567  -0.1172  *** 
## sqrt(vi2)    0.9419  0.0917  10.2771  <.0001   0.7623   1.1215  *** 
## 
## ---
## Signif. codes:  0 '***' 0.001 '**' 0.01 '*' 0.05 '.' 0.1 ' ' 1
```

```
##note that, per Nakagawa 2022, if slope of sqrt(vi2) [SE] is significant,
##use variance (it is significant, so use model below)

mpbias2 <- rma.mv(yi, vi2, mods = ~ vi2,
                 random = list(~1|Citation.number, ~1|ind_id),
                 data = fulldat2)
summary(mpbias2)
```

```
## 
## Multivariate Meta-Analysis Model (k = 1832; method: REML)
## 
##     logLik    Deviance         AIC         BIC        AICc 
## -2765.0048   5530.0095   5538.0095   5560.0578   5538.0315   
## 
## Variance Components:
## 
##             estim    sqrt  nlvls  fixed           factor 
## sigma^2.1  0.6743  0.8212    579     no  Citation.number 
## sigma^2.2  0.3119  0.5585   1832     no           ind_id 
## 
## Test for Residual Heterogeneity:
## QE(df = 1830) = 1453106.6393, p-val < .0001
## 
## Test of Moderators (coefficient 2):
## QM(df = 1) = 102.9261, p-val < .0001
## 
## Model Results:
## 
##          estimate      se     zval    pval    ci.lb   ci.ub 
## intrcpt    0.0453  0.0458   0.9909  0.3217  -0.0443  0.1350      
## vi2        0.4760  0.0469  10.1453  <.0001   0.3841  0.5680  *** 
## 
## ---
## Signif. codes:  0 '***' 0.001 '**' 0.01 '*' 0.05 '.' 0.1 ' ' 1
```

```
##Vi2 is still significant, but that's alright.
##intercept is NOT significantly different from zero, suggesting that 
##there is not asymmetry in the funnel plot, which means there is no small-study
##publication bias present

##time lag analysis
x_numbers <- regmatches(fulldat2$Citation, gregexpr("[[:digit:]]+", fulldat2$Citation))  # Apply gregexpr & regmatches

x_numhold <- list()
for(i in 1:length(x_numbers)){
  if(is_empty(x_numbers[[i]])){
    x_numhold[[i]] = "None"
  } else {
    x_numhold[[i]] = grep("20|19", names(which(sapply(x_numbers[[i]], nchar) == 4)), value = T)
  }
}

x_numhold[[28]] = 2012
x_numhold[[29]] = 2012
x_numhold[[139]] = 2014
x_numhold[[172]] = 2015
x_numhold[[177]] = 2017
x_numhold[[204]] = 2016
x_numhold[[205]] = 2016
x_numhold[[206]] = 2016
x_numhold[[282]] = 2006
x_numhold[[545]] = 2008
x_numhold[[661]] = 2012
x_numhold[[724]] = 2004
x_numhold[[934]] = 2006
x_numhold[[975]] = 2001
x_numhold[[1059]] = 2013
x_numhold[[1102]] = 2013
x_numhold[[1103]] = 2013
x_numhold[[1104]] = 2013
x_numhold[[1162]] = 2013
x_numhold[[1163]] = 2013
x_numhold[[1164]] = 2013
x_numhold[[1165]] = 2013
x_numhold[[1339]] = 2014
x_numhold[[1422]] = 2016
x_numhold[[1518]] = 2015
x_numhold[[1519]] = 2014
x_numhold[[1520]] = 2014
x_numhold[[1534]] = 2013
x_numhold[[1572]] = 2017

fulldat2$Year = as.numeric(unlist(x_numhold))

##time-lag bias model
tlbias <- rma.mv(yi, vi2, mods = ~ Year,
                  random = list(~1|Citation.number, ~1|ind_id),
                  data = fulldat2)
summary(tlbias)
```

```
## 
## Multivariate Meta-Analysis Model (k = 1832; method: REML)
## 
##     logLik    Deviance         AIC         BIC        AICc 
## -2805.0796   5610.1592   5618.1592   5640.2075   5618.1811   
## 
## Variance Components:
## 
##             estim    sqrt  nlvls  fixed           factor 
## sigma^2.1  0.7114  0.8435    579     no  Citation.number 
## sigma^2.2  0.3194  0.5651   1832     no           ind_id 
## 
## Test for Residual Heterogeneity:
## QE(df = 1830) = 1452368.1452, p-val < .0001
## 
## Test of Moderators (coefficient 2):
## QM(df = 1) = 21.9951, p-val < .0001
## 
## Model Results:
## 
##          estimate       se     zval    pval    ci.lb    ci.ub 
## intrcpt   70.5183  14.9891   4.7046  <.0001  41.1402  99.8964  *** 
## Year      -0.0350   0.0075  -4.6899  <.0001  -0.0496  -0.0203  *** 
## 
## ---
## Signif. codes:  0 '***' 0.001 '**' 0.01 '*' 0.05 '.' 0.1 ' ' 1
```

```
##significant effect of Year value indicates that as time goes on,
##less strong positive effects are published
##

##However, visualizing the data suggests that the effect may not be analytically
##important; a single study from 1960s had relatively strong positive effect;
##additionally, handful of strongly negative effect sizes since 2010 seems to
##be driving effect as well; may be an indication of publication bias against
##reduction in disease early in dataset; however, time lag bias is not strength
##of effect, but rather driven by first 20 years of dataset being positive 
##effect sizes. Effect is weak (-0.03 per year), which by the end of the dataset
##suggests an effect size that is not different from zero, so if there is a bias
##it is a bias towards positive effects early on in dataset, not later. This is
##not an enormous concern, because 96% of the dataset was published since 2000. 
##So, shift since 2000 is -0.54, while shift from 1968 is -1.5. 

data.frame(predict(tlbias, matrix(seq(min(fulldat2$Year), max(fulldat2$Year), 1)))) %>%
  mutate(Year = seq(min(fulldat2$Year), max(fulldat2$Year), 1)) %>%
  ggplot(aes(x = Year, y = pred))+
  geom_hline(yintercept = 0, size = 1, linetype = "dashed")+
  geom_point(data = fulldat2, aes(x = Year, y = yi), shape = 21)+
  geom_ribbon(aes(ymin = ci.lb, ymax = ci.ub), alpha = 0.3, fill = "grey43")+
  geom_line(color = "purple", size = 1)+
  scale_x_continuous(breaks = seq(1970,2010,10))+
  labs(y = "Hedge's G")+
  theme_JR()
```

```
ggsave("C:/Users/mikem/Documents/Research/GCD Analyses/TimeLag.tiff",
       dpi = 300,
       width = 5,
       height = 4.75,
       units = "in")
```

Leave one out analysis to test for robustness of our analyses. For
loop is not run due to time constraints, but code is presented. Leave
one out analyses suggests that our results are robust to removal of
individual studies.

```
# 
subdat = subgranmod = NULL
hld = data.frame(beta = 0,
                 se = 0)

# for(i in 1:length(unique(fulldat2$Citation.number))) {
#   subdat = subgranmod = NULL
#   subdat <- fulldat2 %>%
#     filter(Citation.number != unique(fulldat2$Citation.number)[i])
#   subgranmod <- rma.mv(yi, vi2,
#                        random = list(~1|Citation.number, ~1|ind_id),
#                        data = subdat)
#   hld[i,1] = subgranmod$beta[1]
#   hld[i,2] = subgranmod$se
# }
# saveRDS(hld, "G:/Shared drives/GCD of Disease/Final_dat/Leave1Outdat.RDS")

hld <- readRDS("G:/Shared drives/GCD of Disease/Final_dat/Leave1Outdat.RDS")

ggplot(hld, aes(x = 1:nrow(hld), y = beta))+
  geom_errorbar(aes(ymin = beta - se, ymax = beta + se), width = 0)+
  geom_point(color = "light grey", size = 2)+
  geom_hline(yintercept = 0.222, color = "#EA2B1F", size = 1)+
  geom_hline(yintercept = 0.222+0.0442, color = "#EA2B1F", size = 1)+
  geom_hline(yintercept = 0.222-0.0442, color = "#EA2B1F", size = 1)+
  scale_x_continuous(limits = c(0,580), expand = c(0,0))+
  labs(x = "Study",
       y = "Hedge's G")+
  theme_JR()+
  theme(axis.text.x = element_blank(),
        axis.ticks.x = element_blank())
```

```
ggsave("C:/Users/mikem/Documents/Research/GCD Analyses/Leave1out.tiff",
       dpi = 300,
       width = 5,
       height = 4.65,
       units = "in")
```

Forest plots to show distribution of effect sizes and variances among
global change drivers.

### Analyses

#### Step 1: Test for effects of Global Change Drivers (GCDs)

Generally, we find that biodiversity change, introduced species, and
climate change significantly increase disease. Habitat loss/change
significantly decreases disease. Chemical pollution showed a
non-significant effect on disease.

```
options(contrasts = c("contr.treatment","contr.poly"))

m2 <- rma.mv(yi, vi2,
             mods = ~  Global.Change.Driver - 1,
             random = list(~1|Citation.number, ~1|ind_id),
             data = fulldat2)
summary(m2)
```

```
## 
## Multivariate Meta-Analysis Model (k = 1832; method: REML)
## 
##     logLik    Deviance         AIC         BIC        AICc 
## -2786.9159   5573.8318   5587.8318   5626.4048   5587.8934   
## 
## Variance Components:
## 
##             estim    sqrt  nlvls  fixed           factor 
## sigma^2.1  0.6443  0.8027    579     no  Citation.number 
## sigma^2.2  0.3199  0.5656   1832     no           ind_id 
## 
## Test for Residual Heterogeneity:
## QE(df = 1827) = 1430143.7691, p-val < .0001
## 
## Test of Moderators (coefficients 1:5):
## QM(df = 5) = 79.3854, p-val < .0001
## 
## Model Results:
## 
##                          estimate      se     zval    pval    ci.lb   ci.ub 
## Global.Change.DriverBC     0.6735  0.1062   6.3428  <.0001   0.4654  0.8816 
## Global.Change.DriverCC     0.3225  0.0777   4.1486  <.0001   0.1702  0.4749 
## Global.Change.DriverCP     0.0576  0.1100   0.5241  0.6002  -0.1579  0.2732 
## Global.Change.DriverHLC   -0.1343  0.0748  -1.7946  0.0727  -0.2809  0.0124 
## Global.Change.DriverIS     0.5269  0.1224   4.3065  <.0001   0.2871  0.7667 
##  
## Global.Change.DriverBC   *** 
## Global.Change.DriverCC   *** 
## Global.Change.DriverCP 
## Global.Change.DriverHLC    . 
## Global.Change.DriverIS   *** 
## 
## ---
## Signif. codes:  0 '***' 0.001 '**' 0.01 '*' 0.05 '.' 0.1 ' ' 1
```

```
##Set up emmeans reference grid
m2g <- qdrg(object = m2, data = fulldat2)


m2em <- emmeans(m2g, ~ Global.Change.Driver)
CLD(m2em)
```

```
##  Global.Change.Driver  emmean     SE  df asymp.LCL asymp.UCL .group
##  HLC                  -0.1343 0.0748 Inf    -0.281    0.0124  1    
##  CP                    0.0576 0.1100 Inf    -0.158    0.2732  12   
##  CC                    0.3225 0.0777 Inf     0.170    0.4749   23  
##  IS                    0.5269 0.1224 Inf     0.287    0.7667    3  
##  BC                    0.6735 0.1062 Inf     0.465    0.8816    3  
## 
## Confidence level used: 0.95 
## P value adjustment: tukey method for comparing a family of 5 estimates 
## significance level used: alpha = 0.05
```

.

#### Step 2: Test for Subfactors: i.e., subgroupings of ALL GCDs

##### Information on Step 2

- In Introduced Species:
  - Spillover is own subfactor
  - Split between native to invasive and invasive to native
  - Spillback is own subfactor

We see that only a small handful of subfactors actually significantly
affect disease - Biodiversity loss, mean temperature, CO_2_,
and enemy release increase disease - Urbanization reduces disease -
Fungicides marginally increase disease (p = 0.0676) - All other
subfactors have non-significant effects on disease

```
mGCDriver_sub <- rma.mv(yi, vi2,
             mods = ~  Effect_Reduced_Update - 1,
             random = list(~1|Citation.number, ~1|ind_id),
             data = fulldat2)
summary(mGCDriver_sub)
```

```
## 
## Multivariate Meta-Analysis Model (k = 1832; method: REML)
## 
##     logLik    Deviance         AIC         BIC        AICc 
## -2753.5735   5507.1471   5553.1471   5679.6847   5553.7649   
## 
## Variance Components:
## 
##             estim    sqrt  nlvls  fixed           factor 
## sigma^2.1  0.6264  0.7915    579     no  Citation.number 
## sigma^2.2  0.3212  0.5667   1832     no           ind_id 
## 
## Test for Residual Heterogeneity:
## QE(df = 1811) = 1251142.9619, p-val < .0001
## 
## Test of Moderators (coefficients 1:21):
## QM(df = 21) = 106.8349, p-val < .0001
## 
## Model Results:
## 
##                                                       estimate      se     zval 
## Effect_Reduced_UpdateBiodiversity Gradient              0.1548  0.2156   0.7181 
## Effect_Reduced_UpdateBiodiversity Loss                  0.8180  0.1173   6.9732 
## Effect_Reduced_UpdateClimate                            0.2517  0.1813   1.3880 
## Effect_Reduced_UpdateCO2                                0.4360  0.1693   2.5753 
## Effect_Reduced_UpdateDeforestation                      0.2145  0.2220   0.9665 
## Effect_Reduced_UpdateEnemy Release                      0.6673  0.1399   4.7711 
## Effect_Reduced_UpdateFertilizer                         0.0579  0.1974   0.2934 
## Effect_Reduced_UpdateForest Fragmentation               0.1759  0.2468   0.7127 
## Effect_Reduced_UpdateFungicide                          1.1570  0.6337   1.8259 
## Effect_Reduced_UpdateHerbicide                          0.0055  0.1892   0.0292 
## Effect_Reduced_UpdateInsecticide                       -0.0300  0.1747  -0.1720 
## Effect_Reduced_UpdateMean Temperature                   0.3643  0.0995   3.6600 
## Effect_Reduced_UpdateMetal                              0.2765  0.2561   1.0797 
## Effect_Reduced_UpdatePolycyclic Aromatic Hydrocarbon   -0.2670  0.5129  -0.5205 
## Effect_Reduced_UpdatePrecipitation                      0.0150  0.1829   0.0820 
## Effect_Reduced_UpdateSpillback                          0.0308  0.4387   0.0702 
## Effect_Reduced_UpdateSpillover                          0.1277  0.2717   0.4699 
## Effect_Reduced_UpdateSulfur Containing Compound        -0.4564  0.7976  -0.5722 
## Effect_Reduced_UpdateTemperature Variability or ENSO    0.3158  0.2875   1.0986 
## Effect_Reduced_UpdateUrbanization                      -0.2087  0.0813  -2.5668 
## Effect_Reduced_UpdateUVB                                0.5394  0.4616   1.1685 
##                                                         pval    ci.lb    ci.ub 
## Effect_Reduced_UpdateBiodiversity Gradient            0.4727  -0.2677   0.5773 
## Effect_Reduced_UpdateBiodiversity Loss                <.0001   0.5881   1.0479 
## Effect_Reduced_UpdateClimate                          0.1651  -0.1037   0.6071 
## Effect_Reduced_UpdateCO2                              0.0100   0.1042   0.7678 
## Effect_Reduced_UpdateDeforestation                    0.3338  -0.2205   0.6496 
## Effect_Reduced_UpdateEnemy Release                    <.0001   0.3932   0.9414 
## Effect_Reduced_UpdateFertilizer                       0.7692  -0.3290   0.4448 
## Effect_Reduced_UpdateForest Fragmentation             0.4760  -0.3079   0.6597 
## Effect_Reduced_UpdateFungicide                        0.0679  -0.0849   2.3989 
## Effect_Reduced_UpdateHerbicide                        0.9767  -0.3654   0.3764 
## Effect_Reduced_UpdateInsecticide                      0.8634  -0.3724   0.3123 
## Effect_Reduced_UpdateMean Temperature                 0.0003   0.1692   0.5593 
## Effect_Reduced_UpdateMetal                            0.2803  -0.2254   0.7785 
## Effect_Reduced_UpdatePolycyclic Aromatic Hydrocarbon  0.6027  -1.2722   0.7383 
## Effect_Reduced_UpdatePrecipitation                    0.9346  -0.3435   0.3735 
## Effect_Reduced_UpdateSpillback                        0.9440  -0.8290   0.8906 
## Effect_Reduced_UpdateSpillover                        0.6384  -0.4048   0.6601 
## Effect_Reduced_UpdateSulfur Containing Compound       0.5672  -2.0195   1.1068 
## Effect_Reduced_UpdateTemperature Variability or ENSO  0.2719  -0.2476   0.8793 
## Effect_Reduced_UpdateUrbanization                     0.0103  -0.3681  -0.0493 
## Effect_Reduced_UpdateUVB                              0.2426  -0.3653   1.4440 
##  
## Effect_Reduced_UpdateBiodiversity Gradient 
## Effect_Reduced_UpdateBiodiversity Loss                *** 
## Effect_Reduced_UpdateClimate 
## Effect_Reduced_UpdateCO2                                * 
## Effect_Reduced_UpdateDeforestation 
## Effect_Reduced_UpdateEnemy Release                    *** 
## Effect_Reduced_UpdateFertilizer 
## Effect_Reduced_UpdateForest Fragmentation 
## Effect_Reduced_UpdateFungicide                          . 
## Effect_Reduced_UpdateHerbicide 
## Effect_Reduced_UpdateInsecticide 
## Effect_Reduced_UpdateMean Temperature                 *** 
## Effect_Reduced_UpdateMetal 
## Effect_Reduced_UpdatePolycyclic Aromatic Hydrocarbon 
## Effect_Reduced_UpdatePrecipitation 
## Effect_Reduced_UpdateSpillback 
## Effect_Reduced_UpdateSpillover 
## Effect_Reduced_UpdateSulfur Containing Compound 
## Effect_Reduced_UpdateTemperature Variability or ENSO 
## Effect_Reduced_UpdateUrbanization                       * 
## Effect_Reduced_UpdateUVB 
## 
## ---
## Signif. codes:  0 '***' 0.001 '**' 0.01 '*' 0.05 '.' 0.1 ' ' 1
```

#### Step 3: Test for Interactions between GCDs and other factors:

##### Checking for generalities of the patterns

Checking for main and interactive effects between study, host, and
parasite factors and the main global change drivers.

##### Host/Parasite Endpoint

There are differences between host and parasite endpoints in response
to biodiversity change, introduced species, and chemical pollution.
Habitat loss and climate change indicate no differences in response
according to host/parasite endpoint. For BC, IS, and CP, all show that
parasite endpoints increase more in response to GCD than host
endpoints.

```
##Number of replicates
nrow(fulldat2)
```

```
## [1] 1832
```

```
##Main effect of endpoint
mEndpoint <- rma.mv(yi, vi2,
                  mods = ~ Endpoint_Host_Parasite + Global.Change.Driver,
                  random = list(~1|Citation.number, ~1|ind_id),
                  data=fulldat2, method = "ML")
mEndpointIntML <- rma.mv(yi, vi2,
                  mods = ~ Endpoint_Host_Parasite * Global.Change.Driver,
                  random = list(~1|Citation.number, ~1|ind_id),
                  data=fulldat2, method = "ML")

##different; interaction is significant
anova(mEndpoint, mEndpointIntML)
```

```
## 
##         df       AIC       BIC      AICc     logLik     LRT   pval         QE 
## Full    12 5587.4287 5653.5867 5587.6002 -2781.7144                81730.4066 
## Reduced  8 5601.1304 5645.2357 5601.2094 -2792.5652 21.7017 0.0002 92546.8452
```

```
mEndpointInt <- rma.mv(yi, vi2,
       mods = ~ Endpoint_Host_Parasite * Global.Change.Driver,
       random = list(~1|Citation.number, ~1|ind_id),
       data=fulldat2)


##Currently, testing whether the effect of GCDs are similar across endpoints
##Set up emmeans reference grid
mEndg <- qdrg(object = mEndpointInt, data = fulldat2)

mEndgem <- emmeans(mEndg, ~ Endpoint_Host_Parasite|Global.Change.Driver)

##Host and parasite are different in BC, CP, and IS
pairs(mEndgem)
```

```
## Global.Change.Driver = BC:
##  contrast        estimate    SE  df z.ratio p.value
##  Host - Parasite   -0.845 0.324 Inf -2.605  0.0092 
## 
## Global.Change.Driver = CC:
##  contrast        estimate    SE  df z.ratio p.value
##  Host - Parasite    0.197 0.160 Inf  1.233  0.2177 
## 
## Global.Change.Driver = CP:
##  contrast        estimate    SE  df z.ratio p.value
##  Host - Parasite   -0.374 0.162 Inf -2.311  0.0208 
## 
## Global.Change.Driver = HLC:
##  contrast        estimate    SE  df z.ratio p.value
##  Host - Parasite    0.776 0.487 Inf  1.592  0.1113 
## 
## Global.Change.Driver = IS:
##  contrast        estimate    SE  df z.ratio p.value
##  Host - Parasite   -1.124 0.365 Inf -3.077  0.0021
```

##### Host Taxon

Model indicates a significant interaction, but post host test
indicates that only amphibian/reptile is different from arthropods and
plants in response to climate change. No other differences among host
taxa.

```
##Number of replicates
nrow(dathost)
```

```
## [1] 1499
```

```
mHost_Taxon <- rma.mv(yi, vi2,
                  mods = ~ Native.host.type_reduced_Final + Global.Change.Driver,
                  random = list(~1|Citation.number, ~1|ind_id),
                  data=dathost, method = "ML")
mHost_TaxonIntML <- rma.mv(yi, vi2,
                  mods = ~ Native.host.type_reduced_Final * Global.Change.Driver,
                  random = list(~1|Citation.number, ~1|ind_id),
                  data=dathost, method = "ML")

##different
anova(mHost_Taxon, mHost_TaxonIntML)
```

```
## 
##         df       AIC       BIC      AICc     logLik     LRT   pval           QE 
## Full    22 4506.8081 4623.6843 4507.4938 -2231.4041                1347516.2259 
## Reduced 10 4504.5368 4557.6623 4504.6846 -2242.2684 21.7287 0.0407 1348532.4477
```

```
mHost_TaxonInt <- rma.mv(yi, vi2,
       mods = ~ Native.host.type_reduced_Final * Global.Change.Driver,
       random = list(~1|Citation.number, ~1|ind_id),
       data=dathost)


##Currently, testing whether the effect of GCDs are similar across endpoints
##Set up emmeans reference grid
mHost_Taxong <- qdrg(object = mHost_TaxonInt, data = dathost)

mHost_Taxongem <- emmeans(mHost_Taxong, ~ Native.host.type_reduced_Final|Global.Change.Driver)

##Only different in CC
pairs(mHost_Taxongem)
```

```
## Global.Change.Driver = BC:
##  contrast                        estimate    SE  df z.ratio p.value
##  (Amphibian/Reptile) - Arthropod  0.20530 0.611 Inf  0.336  0.9869 
##  (Amphibian/Reptile) - Mammal     0.63427 0.405 Inf  1.567  0.3978 
##  (Amphibian/Reptile) - Plant      0.21861 0.396 Inf  0.553  0.9459 
##  Arthropod - Mammal               0.42897 0.511 Inf  0.840  0.8355 
##  Arthropod - Plant                0.01331 0.512 Inf  0.026  1.0000 
##  Mammal - Plant                  -0.41566 0.230 Inf -1.804  0.2712 
## 
## Global.Change.Driver = CC:
##  contrast                        estimate    SE  df z.ratio p.value
##  (Amphibian/Reptile) - Arthropod  1.08925 0.421 Inf  2.590  0.0473 
##  (Amphibian/Reptile) - Mammal     0.28416 0.293 Inf  0.971  0.7661 
##  (Amphibian/Reptile) - Plant      0.74025 0.270 Inf  2.743  0.0310 
##  Arthropod - Mammal              -0.80509 0.380 Inf -2.120  0.1467 
##  Arthropod - Plant               -0.34900 0.363 Inf -0.963  0.7706 
##  Mammal - Plant                   0.45609 0.201 Inf  2.274  0.1040 
## 
## Global.Change.Driver = CP:
##  contrast                        estimate    SE  df z.ratio p.value
##  (Amphibian/Reptile) - Arthropod  0.00451 0.303 Inf  0.015  1.0000 
##  (Amphibian/Reptile) - Mammal    -0.88655 0.532 Inf -1.668  0.3409 
##  (Amphibian/Reptile) - Plant      0.04252 0.273 Inf  0.156  0.9987 
##  Arthropod - Mammal              -0.89106 0.550 Inf -1.620  0.3674 
##  Arthropod - Plant                0.03801 0.308 Inf  0.124  0.9993 
##  Mammal - Plant                   0.92906 0.534 Inf  1.740  0.3029 
## 
## Global.Change.Driver = HLC:
##  contrast                        estimate    SE  df z.ratio p.value
##  (Amphibian/Reptile) - Arthropod -0.05760 0.431 Inf -0.134  0.9991 
##  (Amphibian/Reptile) - Mammal     0.00621 0.369 Inf  0.017  1.0000 
##  (Amphibian/Reptile) - Plant      0.06949 0.684 Inf  0.102  0.9996 
##  Arthropod - Mammal               0.06381 0.250 Inf  0.255  0.9942 
##  Arthropod - Plant                0.12708 0.628 Inf  0.202  0.9971 
##  Mammal - Plant                   0.06327 0.587 Inf  0.108  0.9996 
## 
## Global.Change.Driver = IS:
##  contrast                        estimate    SE  df z.ratio p.value
##  (Amphibian/Reptile) - Arthropod -0.48259 0.678 Inf -0.712  0.8925 
##  (Amphibian/Reptile) - Mammal    -0.61050 0.791 Inf -0.772  0.8670 
##  (Amphibian/Reptile) - Plant     -0.78958 0.558 Inf -1.415  0.4901 
##  Arthropod - Mammal              -0.12791 0.709 Inf -0.180  0.9979 
##  Arthropod - Plant               -0.30699 0.435 Inf -0.706  0.8947 
##  Mammal - Plant                  -0.17908 0.595 Inf -0.301  0.9905 
## 
## P value adjustment: tukey method for comparing a family of 4 estimates
```

##### Parasite Taxon

Some differences among parasite taxon. Interaction model suggests
only differences in responses among taxa are within biodiversity change,
introduced species, and Habitat loss and change - Helminths respond more
positively than bacteria, fungi, and viruses to biodiversity change -
Fungi respond more positively than viruses to introduced species

```
##Number of replicates
nrow(datenemy)
```

```
## [1] 1596
```

```
mEnemy_Taxon <- rma.mv(yi, vi2,
                  mods = ~ Enemy.type_further_reduced_Final + Global.Change.Driver,
                  random = list(~1|Citation.number, ~1|ind_id),
                  data=datenemy, method = "ML")
mEnemy_TaxonIntML <- rma.mv(yi, vi2,
                  mods = ~ Enemy.type_further_reduced_Final * Global.Change.Driver,
                  random = list(~1|Citation.number, ~1|ind_id),
                  data=datenemy, method = "ML")

##very different
anova(mEnemy_Taxon, mEnemy_TaxonIntML)
```

```
## 
##         df       AIC       BIC      AICc     logLik     LRT   pval           QE 
## Full    27 4923.0839 5068.2158 4924.0482 -2434.5419                1340381.6851 
## Reduced 11 4945.5902 5004.7180 4945.7569 -2461.7951 54.5063 <.0001 1342144.7469
```

```
mEnemy_TaxonInt <- rma.mv(yi, vi2,
                      mods = ~ Enemy.type_further_reduced_Final * Global.Change.Driver,
                      random = list(~1|Citation.number, ~1|ind_id), data=datenemy)

##Set up emmeans reference grid
mEnemyg <- qdrg(object = mEnemy_TaxonInt, data = datenemy)

mEnemygem <- emmeans(mEnemyg, ~ Enemy.type_further_reduced_Final|Global.Change.Driver)

#
pairs((mEnemygem))
```

```
## Global.Change.Driver = BC:
##  contrast            estimate    SE  df z.ratio p.value
##  Bacteria - Fungi     -0.0770 0.312 Inf -0.247  0.9992 
##  Bacteria - Helminth  -0.8450 0.287 Inf -2.949  0.0265 
##  Bacteria - Protist   -0.3516 0.303 Inf -1.162  0.7731 
##  Bacteria - Virus     -0.0957 0.315 Inf -0.303  0.9982 
##  Fungi - Helminth     -0.7680 0.270 Inf -2.847  0.0357 
##  Fungi - Protist      -0.2746 0.333 Inf -0.823  0.9235 
##  Fungi - Virus        -0.0187 0.290 Inf -0.064  1.0000 
##  Helminth - Protist    0.4934 0.282 Inf  1.752  0.4020 
##  Helminth - Virus      0.7493 0.263 Inf  2.853  0.0351 
##  Protist - Virus       0.2559 0.321 Inf  0.798  0.9312 
## 
## Global.Change.Driver = CC:
##  contrast            estimate    SE  df z.ratio p.value
##  Bacteria - Fungi      0.4091 0.249 Inf  1.643  0.4698 
##  Bacteria - Helminth   0.4718 0.296 Inf  1.593  0.5017 
##  Bacteria - Protist    0.2241 0.293 Inf  0.765  0.9405 
##  Bacteria - Virus      0.6059 0.378 Inf  1.603  0.4954 
##  Fungi - Helminth      0.0626 0.233 Inf  0.269  0.9989 
##  Fungi - Protist      -0.1850 0.235 Inf -0.786  0.9348 
##  Fungi - Virus         0.1967 0.334 Inf  0.589  0.9768 
##  Helminth - Protist   -0.2477 0.282 Inf -0.878  0.9051 
##  Helminth - Virus      0.1341 0.368 Inf  0.364  0.9963 
##  Protist - Virus       0.3817 0.369 Inf  1.033  0.8400 
## 
## Global.Change.Driver = CP:
##  contrast            estimate    SE  df z.ratio p.value
##  Bacteria - Fungi      0.4897 0.310 Inf  1.578  0.5114 
##  Bacteria - Helminth   0.3815 0.367 Inf  1.040  0.8369 
##  Bacteria - Protist    0.6006 0.343 Inf  1.750  0.4031 
##  Bacteria - Virus      0.5417 0.355 Inf  1.525  0.5461 
##  Fungi - Helminth     -0.1082 0.318 Inf -0.340  0.9971 
##  Fungi - Protist       0.1109 0.306 Inf  0.362  0.9963 
##  Fungi - Virus         0.0520 0.299 Inf  0.174  0.9998 
##  Helminth - Protist    0.2191 0.375 Inf  0.585  0.9774 
##  Helminth - Virus      0.1602 0.385 Inf  0.416  0.9937 
##  Protist - Virus      -0.0589 0.353 Inf -0.167  0.9998 
## 
## Global.Change.Driver = HLC:
##  contrast            estimate    SE  df z.ratio p.value
##  Bacteria - Fungi      0.3436 0.270 Inf  1.274  0.7070 
##  Bacteria - Helminth   0.3629 0.190 Inf  1.908  0.3130 
##  Bacteria - Protist    0.1026 0.155 Inf  0.660  0.9647 
##  Bacteria - Virus     -0.2033 0.175 Inf -1.161  0.7738 
##  Fungi - Helminth      0.0192 0.289 Inf  0.067  1.0000 
##  Fungi - Protist      -0.2410 0.266 Inf -0.905  0.8949 
##  Fungi - Virus        -0.5469 0.273 Inf -2.004  0.2641 
##  Helminth - Protist   -0.2602 0.153 Inf -1.703  0.4318 
##  Helminth - Virus     -0.5662 0.213 Inf -2.655  0.0609 
##  Protist - Virus      -0.3059 0.184 Inf -1.661  0.4583 
## 
## Global.Change.Driver = IS:
##  contrast            estimate    SE  df z.ratio p.value
##  Bacteria - Fungi     -1.2217 0.504 Inf -2.425  0.1085 
##  Bacteria - Helminth  -1.0827 0.598 Inf -1.810  0.3677 
##  Bacteria - Protist    0.5311 0.846 Inf  0.628  0.9707 
##  Bacteria - Virus     -0.4735 0.534 Inf -0.886  0.9021 
##  Fungi - Helminth      0.1389 0.350 Inf  0.397  0.9948 
##  Fungi - Protist       1.7527 0.693 Inf  2.528  0.0845 
##  Fungi - Virus         0.7482 0.193 Inf  3.883  0.0010 
##  Helminth - Protist    1.6138 0.748 Inf  2.158  0.1959 
##  Helminth - Virus      0.6092 0.383 Inf  1.589  0.5046 
##  Protist - Virus      -1.0046 0.711 Inf -1.413  0.6190 
## 
## P value adjustment: tukey method for comparing a family of 5 estimates
```

##### Vector

Significant difference between vector and non-vector parasites in
Habitat Loss/Change and Climate Change. In Habitat Loss/Change, vector
parasites decrease, while non vectors do not change (CIs overlap 0). In
Climate Change, vectors are predicted to increase more than non
vectors.

```
##Number of replicates
nrow(vecdat)
```

```
## [1] 1142
```

```
mVector <- rma.mv(yi, vi2,
                  mods = ~ Vector + Global.Change.Driver,
                  random = list(~1|Citation.number, ~1|ind_id),
                  data=vecdat, method = "ML")
mVectorInt <- rma.mv(yi, vi2,
                  mods = ~ Vector * Global.Change.Driver,
                  random = list(~1|Citation.number, ~1|ind_id),
                  data=vecdat, method = "ML")
## different
anova(mVector, mVectorInt)
```

```
## 
##         df       AIC       BIC      AICc     logLik    LRT   pval           QE 
## Full     8 3614.1941 3654.5184 3614.3212 -1799.0971               1407984.5332 
## Reduced  6 3619.3500 3649.5932 3619.4240 -1803.6750 9.1559 0.0103 1412932.2521
```

```
mVectorInt <- rma.mv(yi, vi2,
                      mods = ~ Vector * Global.Change.Driver,
                      random = list(~1|Citation.number, ~1|ind_id), data=vecdat)

##Set up emmeans reference grid
mVectorg <- qdrg(object = mVectorInt, data = vecdat)

mVectorgem <- emmeans(mVectorg, ~ Vector|Global.Change.Driver)

#
pairs((mVectorgem))
```

```
## Global.Change.Driver = CC:
##  contrast estimate    SE  df z.ratio p.value
##  No - Yes   -0.534 0.265 Inf -2.013  0.0442 
## 
## Global.Change.Driver = HLC:
##  contrast estimate    SE  df z.ratio p.value
##  No - Yes    0.433 0.181 Inf  2.398  0.0165 
## 
## Global.Change.Driver = IS:
##  contrast estimate    SE  df z.ratio p.value
##  No - Yes    0.343 0.684 Inf  0.501  0.6161
```

##### Vector-borne

No generalizable differences between vector-borne and
non-vector-borne diseases, but within biodiversity change, there are
differences. In biodiversity change, vector-borne diseases respond
significantly stronger (positive) to biodiversity change than
non-vector-borne diseases. This is consistent with previous work from
Dave Civitello, suggesting that dilution effect is more prominent in
vector-borne diseases.

```
##Number of replicates
nrow(vecbordat)
```

```
## [1] 1753
```

```
mVectorBorne <- rma.mv(yi, vi2,
                  mods = ~ Vector.borne + Global.Change.Driver,
                  random = list(~1|Citation.number, ~1|ind_id),
                  data=vecbordat, method = "ML")
mVectorBorneIntML <- rma.mv(yi, vi2,
                  mods = ~ Vector.borne * Global.Change.Driver,
                  random = list(~1|Citation.number, ~1|ind_id),
                  data=vecbordat, method = "ML")

anova(mVectorBorne, mVectorBorneIntML)
```

```
## 
##         df       AIC       BIC      AICc     logLik     LRT   pval           QE 
## Full    12 5393.8477 5459.4767 5394.0270 -2684.9238                1420554.5059 
## Reduced  8 5397.5505 5441.3032 5397.6331 -2690.7752 11.7028 0.0197 1421677.7435
```

```
mVectorBorneInt <- rma.mv(yi, vi2,
                      mods = ~ Vector.borne * Global.Change.Driver,
                      random = list(~1|Citation.number, ~1|ind_id), data=vecbordat)

##Set up emmeans reference grid
mVectorBg <- qdrg(object = mVectorBorneInt, data = vecbordat)

mVectorBgem <- emmeans(mVectorBg, ~ Vector.borne|Global.Change.Driver)

#Only difference is in BC
pairs((mVectorBgem))
```

```
## Global.Change.Driver = BC:
##  contrast estimate    SE  df z.ratio p.value
##  No - Yes   -0.427 0.170 Inf -2.510  0.0121 
## 
## Global.Change.Driver = CC:
##  contrast estimate    SE  df z.ratio p.value
##  No - Yes    0.205 0.172 Inf  1.190  0.2341 
## 
## Global.Change.Driver = CP:
##  contrast estimate    SE  df z.ratio p.value
##  No - Yes   -0.143 0.265 Inf -0.540  0.5889 
## 
## Global.Change.Driver = HLC:
##  contrast estimate    SE  df z.ratio p.value
##  No - Yes    0.132 0.117 Inf  1.133  0.2574 
## 
## Global.Change.Driver = IS:
##  contrast estimate    SE  df z.ratio p.value
##  No - Yes    0.505 0.320 Inf  1.582  0.1138
```

##### Ecto- v. Endoparasites

No differences between ecto- and edo- parasites in response to
biodiversity change, chemical pollution, and climate change. For
Introduced Species, ecto-parasites respond positively, while
endo-parasites do not change. Conversely, for habitat loss,
ecto-parasites respond negatively, while endo-parasites do not
change.

```
##Number of replicates
nrow(ectodat)
```

```
## [1] 1808
```

```
mEctoparasite <- rma.mv(yi, vi2,
                  mods = ~ Ectoparasite + Global.Change.Driver,
                  random = list(~1|Citation.number, ~1|ind_id),
                  data=ectodat, method = "ML")
mEctoparasiteIntML <- rma.mv(yi, vi2,
                  mods = ~ Ectoparasite * Global.Change.Driver,
                  random = list(~1|Citation.number, ~1|ind_id),
                  data=ectodat, method = "ML")

anova(mEctoparasite, mEctoparasiteIntML)
```

```
## 
##         df       AIC       BIC      AICc     logLik     LRT   pval           QE 
## Full    12 5523.6426 5589.6423 5523.8164 -2749.8213                1363468.1603 
## Reduced  8 5547.4134 5591.4132 5547.4934 -2765.7067 31.7708 <.0001 1381743.0860
```

```
mEctoparasiteInt <- rma.mv(yi, vi2,
                      mods = ~ Ectoparasite * Global.Change.Driver,
                      random = list(~1|Citation.number, ~1|ind_id), data=ectodat)

##Set up emmeans reference grid
mEctoparasiteBg <- qdrg(object = mEctoparasiteInt, data = ectodat)

mEctoparasiteBgem <- emmeans(mEctoparasiteBg, ~ Ectoparasite|Global.Change.Driver)

#Only difference is in IS
pairs((mEctoparasiteBgem))
```

```
## Global.Change.Driver = BC:
##  contrast estimate    SE  df z.ratio p.value
##  No - Yes   0.3657 0.222 Inf  1.644  0.1002 
## 
## Global.Change.Driver = CC:
##  contrast estimate    SE  df z.ratio p.value
##  No - Yes  -0.0484 0.156 Inf -0.310  0.7565 
## 
## Global.Change.Driver = CP:
##  contrast estimate    SE  df z.ratio p.value
##  No - Yes   0.3419 0.222 Inf  1.542  0.1231 
## 
## Global.Change.Driver = HLC:
##  contrast estimate    SE  df z.ratio p.value
##  No - Yes   0.3271 0.145 Inf  2.259  0.0239 
## 
## Global.Change.Driver = IS:
##  contrast estimate    SE  df z.ratio p.value
##  No - Yes  -0.7305 0.156 Inf -4.683  <.0001
```

##### Human v. Non-human Parasites

There are no differences in human v. non-human parasites in their
response to global change drivers. Overall, probably a good thing, but
not too exciting.

```
###### Human Parasites
##Number of replicates
nrow(humdat)
```

```
## [1] 1788
```

```
mHuman <- rma.mv(yi, vi2,
                  mods = ~ Human.ParasiteBoth.NA + Global.Change.Driver,
                  random = list(~1|Citation.number, ~1|ind_id),
                  data=humdat, method = "ML")
mHumanInt <- rma.mv(yi, vi2,
                  mods = ~ Human.ParasiteBoth.NA * Global.Change.Driver,
                  random = list(~1|Citation.number, ~1|ind_id),
                  data=humdat, method = "ML")

#Not significant
anova(mHuman, mHumanInt)
```

```
## 
##         df       AIC       BIC      AICc     logLik    LRT   pval           QE 
## Full    12 5494.7852 5560.6514 5494.9609 -2735.3926               1421190.1133 
## Reduced  8 5494.5981 5538.5090 5494.6791 -2739.2991 7.8130 0.0987 1428206.6534
```

```
anova(mHuman, btt = 2)
```

```
## 
## Test of Moderators (coefficient 2):
## QM(df = 1) = 0.0003, p-val = 0.9855
```

```
mHuman <- rma.mv(yi, vi2,
                  mods = ~ Human.ParasiteBoth.NA + Global.Change.Driver,
                  random = list(~1|Citation.number, ~1|ind_id),
                  data=humdat, method = "REML")
mHumang <- qdrg(object = mHuman, data = humdat)
```

##### Venue

Lab studies tend to overestimate the effect of changes in
biodiversity on disease. For all other GCDs tested (introduced species,
climate change, and chemical pollution), lab studies were no different
from field studies of these drivers on disease.

```
##Number of replicates
nrow(vendat)
```

```
## [1] 1328
```

```
mVenue <- rma.mv(yi, vi2,
                  mods = ~ VenueNoModel + Global.Change.Driver,
                  random = list(~1|Citation.number, ~1|ind_id),
                  data=vendat, method = "ML")
mVenueIntML <- rma.mv(yi, vi2,
                  mods = ~ VenueNoModel * Global.Change.Driver,
                  random = list(~1|Citation.number, ~1|ind_id),
                  data=vendat, method = "ML")

#Very diff
anova(mVenue, mVenueIntML)
```

```
## 
##         df       AIC       BIC      AICc     logLik     LRT   pval           QE 
## Full    10 4116.3024 4168.2167 4116.4695 -2048.1512                1388297.5489 
## Reduced  7 4134.0635 4170.4035 4134.1483 -2060.0317 23.7610 <.0001 1389095.6276
```

```
mVenueInt <- rma.mv(yi, vi2,
                      mods = ~ VenueNoModel * Global.Change.Driver,
                      random = list(~1|Citation.number, ~1|ind_id), data=vendat)

##Set up emmeans reference grid
mVenueg <- qdrg(object = mVenueInt, data = vendat)

mVenuegem <- emmeans(mVenueg, ~ VenueNoModel|Global.Change.Driver)

#Only difference is in BC
pairs((mVenuegem))
```

```
## Global.Change.Driver = BC:
##  contrast    estimate    SE  df z.ratio p.value
##  Field - Lab   -1.415 0.304 Inf -4.660  <.0001 
## 
## Global.Change.Driver = CC:
##  contrast    estimate    SE  df z.ratio p.value
##  Field - Lab    0.152 0.161 Inf  0.944  0.3450 
## 
## Global.Change.Driver = CP:
##  contrast    estimate    SE  df z.ratio p.value
##  Field - Lab    0.283 0.253 Inf  1.119  0.2632 
## 
## Global.Change.Driver = IS:
##  contrast    estimate    SE  df z.ratio p.value
##  Field - Lab    0.291 0.547 Inf  0.532  0.5950
```

##### Habitat

There is a difference in how different habitats respond to
biodiversity change. Specifically, effects of biodiversity change on
disease is expected to be strongest in marine systems, followed by
freshwater, with terrestrial systems responding relatively weakly.

```
##Number of replicates
nrow(habdat)
```

```
## [1] 1832
```

```
mHabitat <- rma.mv(yi, vi2,
                  mods = ~ Habitat + Global.Change.Driver,
                  random = list(~1|Citation.number, ~1|ind_id),
                  data=habdat, method = "ML")
mHabitatIntML <- rma.mv(yi, vi2,
                  mods = ~ Habitat * Global.Change.Driver,
                  random = list(~1|Citation.number, ~1|ind_id),
                  data=habdat, method = "ML")

#No diff
anova(mHabitat, mHabitatIntML)
```

```
## 
##         df       AIC       BIC      AICc     logLik     LRT   pval           QE 
## Full    17 5580.8652 5674.5890 5581.2026 -2773.4326                1428731.0518 
## Reduced  9 5599.1859 5648.8043 5599.2847 -2790.5929 34.3206 <.0001 1429669.6393
```

```
mHabitatInt <- rma.mv(yi, vi2,
                      mods = ~ Habitat * Global.Change.Driver,
                      random = list(~1|Citation.number, ~1|ind_id), data=habdat)

##Set up emmeans reference grid
mHabitatg <- qdrg(object = mHabitatInt, data = habdat)

mHabitatgem <- emmeans(mHabitatg, ~ Habitat | Global.Change.Driver)

#Only difference is in BC
pairs((mHabitatgem))
```

```
## Global.Change.Driver = BC:
##  contrast                 estimate    SE  df z.ratio p.value
##  Freshwater - Marine        -2.145 0.550 Inf -3.903  0.0003 
##  Freshwater - Terrestrial    0.641 0.257 Inf  2.495  0.0337 
##  Marine - Terrestrial        2.786 0.513 Inf  5.434  <.0001 
## 
## Global.Change.Driver = CC:
##  contrast                 estimate    SE  df z.ratio p.value
##  Freshwater - Marine         0.220 0.268 Inf  0.821  0.6903 
##  Freshwater - Terrestrial    0.112 0.185 Inf  0.606  0.8170 
##  Marine - Terrestrial       -0.108 0.234 Inf -0.464  0.8880 
## 
## Global.Change.Driver = CP:
##  contrast                 estimate    SE  df z.ratio p.value
##  Freshwater - Marine         0.524 0.412 Inf  1.272  0.4111 
##  Freshwater - Terrestrial    0.380 0.220 Inf  1.725  0.1958 
##  Marine - Terrestrial       -0.144 0.419 Inf -0.344  0.9369 
## 
## Global.Change.Driver = HLC:
##  contrast                 estimate    SE  df z.ratio p.value
##  Freshwater - Marine         0.317 1.020 Inf  0.311  0.9482 
##  Freshwater - Terrestrial    0.643 0.303 Inf  2.122  0.0853 
##  Marine - Terrestrial        0.326 0.979 Inf  0.333  0.9405 
## 
## Global.Change.Driver = IS:
##  contrast                 estimate    SE  df z.ratio p.value
##  Freshwater - Marine        -0.147 0.505 Inf -0.292  0.9542 
##  Freshwater - Terrestrial   -0.358 0.363 Inf -0.986  0.5860 
##  Marine - Terrestrial       -0.211 0.397 Inf -0.531  0.8563 
## 
## P value adjustment: tukey method for comparing a family of 3 estimates
```

##### Route

Route (complex or direct) was significantly different for
biodiversity change and habitat loss/change. Very similar patterns to
the vector-borne status results, so likely very similar drivers
there.

```
##Number of replicates
nrow(datRoute)
```

```
## [1] 1746
```

```
mRoute <- rma.mv(yi, vi2,
                  mods = ~ RouteBoth.NA + Global.Change.Driver,
                  random = list(~1|Citation.number, ~1|ind_id),
                  data=datRoute, method = "ML")
mRouteIntML <- rma.mv(yi, vi2,
                  mods = ~ RouteBoth.NA * Global.Change.Driver,
                  random = list(~1|Citation.number, ~1|ind_id),
                  data=datRoute, method = "ML")

#Difference
anova(mRoute, mRouteIntML)
```

```
## 
##         df       AIC       BIC      AICc     logLik     LRT   pval           QE 
## Full    12 5346.9769 5412.5579 5347.1569 -2661.4885                1418609.3451 
## Reduced  8 5354.4335 5398.1542 5354.5164 -2669.2168 15.4566 0.0038 1420568.3432
```

```
mRouteInt <- rma.mv(yi, vi2,
                      mods = ~ RouteBoth.NA * Global.Change.Driver,
                      random = list(~1|Citation.number, ~1|ind_id), data=datRoute)

##Set up emmeans reference grid
mRouteIntg <- qdrg(object = mRouteInt, data = datRoute)

mRouteIntgem <- emmeans(mRouteIntg, ~ RouteBoth.NA | Global.Change.Driver)

#Differences in BC and HLC
pairs((mRouteIntgem))
```

```
## Global.Change.Driver = BC:
##  contrast         estimate    SE  df z.ratio p.value
##  Complex - Direct   0.5600 0.191 Inf  2.930  0.0034 
## 
## Global.Change.Driver = CC:
##  contrast         estimate    SE  df z.ratio p.value
##  Complex - Direct  -0.2250 0.231 Inf -0.975  0.3296 
## 
## Global.Change.Driver = CP:
##  contrast         estimate    SE  df z.ratio p.value
##  Complex - Direct  -0.0364 0.236 Inf -0.154  0.8775 
## 
## Global.Change.Driver = HLC:
##  contrast         estimate    SE  df z.ratio p.value
##  Complex - Direct  -0.2508 0.112 Inf -2.241  0.0250 
## 
## Global.Change.Driver = IS:
##  contrast         estimate    SE  df z.ratio p.value
##  Complex - Direct  -0.4815 0.339 Inf -1.418  0.1560
```

##### Free Living Stages

There was no difference among different types of free living stages
in their response to global change drivers.

```
##Number of replicates
nrow(datFLstage)
```

```
## [1] 1758
```

```
mStage <- rma.mv(yi, vi2,
                  mods = ~ Free.living.stages + Global.Change.Driver,
                  random = list(~1|Citation.number, ~1|ind_id),
                  data=datFLstage, method = "ML")
mStageIntML <- rma.mv(yi, vi2,
                  mods = ~ Free.living.stages * Global.Change.Driver,
                  random = list(~1|Citation.number, ~1|ind_id),
                  data=datFLstage, method = "ML")

#No Difference
anova(mStage, mStageIntML)
```

```
## 
##         df       AIC       BIC      AICc     logLik    LRT   pval           QE 
## Full    12 5409.0930 5474.7562 5409.2718 -2692.5465               1421041.3421 
## Reduced  8 5408.0034 5451.7788 5408.0857 -2696.0017 6.9103 0.1407 1421460.2248
```

```
anova(mStage, btt = 2)
```

```
## 
## Test of Moderators (coefficient 2):
## QM(df = 1) = 0.8073, p-val = 0.3689
```

```
##No difference between free-living and non-free living parasites

mStage <- rma.mv(yi, vi2,
                  mods = ~ Free.living.stages + Global.Change.Driver,
                  random = list(~1|Citation.number, ~1|ind_id),
                  data=datFLstage, method = "REML")
mStageg <- qdrg(object = mStage, data = datFLstage)
```

##### Macroparasites

Parasite size was different for biodiversity change (macroparasites
responded more positively to biodiversity loss) and for Habitat
Loss/Change (macroparasites responded more negatively to habitat
loss/change).

```
##Number of replicates
nrow(datMacroparasite)
```

```
## [1] 1809
```

```
mMacro <- rma.mv(yi, vi2,
                  mods = ~ Macroparasite + Global.Change.Driver,
                  random = list(~1|Citation.number, ~1|ind_id),
                  data=datMacroparasite, method = "ML")
mMacroIntML <- rma.mv(yi, vi2,
                  mods = ~ Macroparasite * Global.Change.Driver,
                  random = list(~1|Citation.number, ~1|ind_id),
                  data=datMacroparasite, method = "ML")

#Difference
anova(mMacro, mMacroIntML)
```

```
## 
##         df       AIC       BIC      AICc     logLik     LRT   pval           QE 
## Full    12 5537.0476 5603.0539 5537.2213 -2756.5238                1420524.4466 
## Reduced  8 5546.7569 5590.7611 5546.8369 -2765.3784 17.7093 0.0014 1422096.5916
```

```
mMacroInt <- rma.mv(yi, vi2,
                  mods = ~ Macroparasite * Global.Change.Driver,
                  random = list(~1|Citation.number, ~1|ind_id),
                  data=datMacroparasite)

##Set up emmeans reference grid
mMacroIntg <- qdrg(object = mMacroInt, data = datMacroparasite)

mMacroIntgem <- emmeans(mMacroIntg, ~ Macroparasite | Global.Change.Driver)

#Differences in BC (IS p = 0.064)
pairs((mMacroIntgem))
```

```
## Global.Change.Driver = BC:
##  contrast estimate    SE  df z.ratio p.value
##  No - Yes  -0.6112 0.175 Inf -3.485  0.0005 
## 
## Global.Change.Driver = CC:
##  contrast estimate    SE  df z.ratio p.value
##  No - Yes  -0.0571 0.176 Inf -0.325  0.7450 
## 
## Global.Change.Driver = CP:
##  contrast estimate    SE  df z.ratio p.value
##  No - Yes   0.0459 0.258 Inf  0.178  0.8585 
## 
## Global.Change.Driver = HLC:
##  contrast estimate    SE  df z.ratio p.value
##  No - Yes   0.2684 0.119 Inf  2.254  0.0242 
## 
## Global.Change.Driver = IS:
##  contrast estimate    SE  df z.ratio p.value
##  No - Yes  -0.1864 0.290 Inf -0.644  0.5198
```

##### Host Thermy

Interestingly, we see a slight difference in disease response between
ecto- and endo- thermic hosts for biodiversity change. For biodiversity
change, this suggests that changes in biodiversity will increase disease
more in ectothermic hosts than in endothermic hosts.

```
##Number of replicates
nrow(datHostThermy)
```

```
## [1] 1832
```

```
mThermy <- rma.mv(yi, vi2,
                  mods = ~ Ectothermic.host + Global.Change.Driver,
                  random = list(~1|Citation.number, ~1|ind_id),
                  data=datHostThermy, method = "ML")
mThermyIntML <- rma.mv(yi, vi2,
                  mods = ~ Ectothermic.host * Global.Change.Driver,
                  random = list(~1|Citation.number, ~1|ind_id),
                  data=datHostThermy, method = "ML")

#Difference!
anova(mThermy, mThermyIntML)
```

```
## 
##         df       AIC       BIC      AICc     logLik     LRT   pval           QE 
## Full    12 5600.8108 5666.9687 5600.9823 -2788.4054                1421492.3174 
## Reduced  8 5603.8897 5647.9950 5603.9687 -2793.9448 11.0789 0.0257 1422926.8905
```

```
mThermyInt <- rma.mv(yi, vi2,
                  mods = ~ Ectothermic.host * Global.Change.Driver,
                  random = list(~1|Citation.number, ~1|ind_id),
                  data=datHostThermy)

##Set up emmeans reference grid
mThermyIntg <- qdrg(object = mThermyInt, data = datHostThermy)

mThermyIntgem <- emmeans(mThermyIntg, ~ Ectothermic.host | Global.Change.Driver)

#Differences in BC and HLC
pairs((mThermyIntgem))
```

```
## Global.Change.Driver = BC:
##  contrast estimate    SE  df z.ratio p.value
##  No - Yes   -0.464 0.203 Inf -2.289  0.0221 
## 
## Global.Change.Driver = CC:
##  contrast estimate    SE  df z.ratio p.value
##  No - Yes    0.264 0.184 Inf  1.433  0.1520 
## 
## Global.Change.Driver = CP:
##  contrast estimate    SE  df z.ratio p.value
##  No - Yes    0.565 0.475 Inf  1.189  0.2343 
## 
## Global.Change.Driver = HLC:
##  contrast estimate    SE  df z.ratio p.value
##  No - Yes   -0.180 0.181 Inf -0.997  0.3190 
## 
## Global.Change.Driver = IS:
##  contrast estimate    SE  df z.ratio p.value
##  No - Yes   -0.624 0.405 Inf -1.541  0.1233
```

#### Step 4: Model Selection:

##### What factors best explain disease response to specific global change drivers?

- Model selection process for each individual GCD
- Only using citation here to stay within the bounds of the
  meta-analysis framework

###### Data prep

- For each individual global change driver, removed all NA values for
  predictors. \*Cchecked for missing cells (i.e. Marine habitats and
  Habitat Loss).
- As such, replicates for each GCD are below.

```
dataBL2 = subset(fulldat2, Global.Change.Driver == "BC")

##Remove all rows (probably 3-5) with NA values in ANY of the categories
dataBL2 = dataBL2 %>%
  filter(complete.cases(Habitat) &
           complete.cases(Enemy.type_further_reduced_Final) &
           complete.cases(Vector.borne) &
           complete.cases(Free.living.stages) &
           complete.cases(Effect_Reduced_Update))

##Drop Birds and Fish (2 reps of each)
dataBL2 = subset(dataBL2, Native.host.type_reduced_Final != "Bird" &
                   Native.host.type_reduced_Final != "Fish")

##Replicates for biodiversity
nrow(dataBL2)
```

```
## [1] 401
```

```
##Climate change
dataCC2 = subset(fulldat2, Global.Change.Driver == "CC")


##Remove all rows with NA values in ANY of the categories
dataCC2 = dataCC2 %>%
  filter(complete.cases(Habitat) &
           complete.cases(Enemy.type_further_reduced_Final) &
           complete.cases(Vector.borne) &
           complete.cases(Native.host.type_reduced_Final) & 
           complete.cases(Ectoparasite) &
           complete.cases(Vector.borne) &
           complete.cases(Free.living.stages) &
           complete.cases(RouteBoth.NA) &
           complete.cases(Human.ParasiteBoth.NA) &
           complete.cases(MacroparasiteBoth.NA) &
           complete.cases(Effect_Reduced_Update))

##Replicates for climate change
nrow(dataCC2)
```

```
## [1] 284
```

```
#Habitat Loss/Change
dataHLC2 = subset(fulldat2, Global.Change.Driver == "HLC")

##Remove all rows with NA values in ANY of the categories
dataHLC2 = dataHLC2 %>%
  filter(complete.cases(Habitat) &
           complete.cases(Enemy.type_further_reduced_Final) &
           complete.cases(Vector.borne) &
           complete.cases(Native.host.type_reduced_Final) & 
           complete.cases(Ectoparasite) &
           complete.cases(Vector.borne) &
           complete.cases(Free.living.stages) &
           complete.cases(RouteBoth.NA) &
           complete.cases(Human.ParasiteBoth.NA) &
           complete.cases(MacroparasiteBoth.NA) &
           complete.cases(Effect_Reduced_Update))

dataHLC2 = subset(dataHLC2, Native.host.type_reduced_Final != "Mollusk" &
                    Native.host.type_reduced_Final != "Fish" &
                    Habitat != "Marine")

##Replicates for habitat loss/change
nrow(dataHLC2)
```

```
## [1] 400
```

```
##Introduced species
dataIS2 = subset(fulldat2, Global.Change.Driver == "IS")

##Remove all rows with NA values in ANY of the categories
dataIS2 = dataIS2 %>%
  filter(complete.cases(Habitat) &
           complete.cases(Enemy.type_further_reduced_Final) &
           complete.cases(Vector.borne) &
           complete.cases(Native.host.type_reduced_Final) & 
           complete.cases(Ectoparasite) &
           complete.cases(Vector.borne) &
           complete.cases(Free.living.stages) &
           complete.cases(RouteBoth.NA) &
           complete.cases(Human.ParasiteBoth.NA) &
           complete.cases(MacroparasiteBoth.NA) &
           complete.cases(Effect_Reduced_Update))

dataIS2 = subset(dataIS2, 
                   Enemy.type_further_reduced_Final != "Bacteria")

##Replicates for introduced species
nrow(dataIS2)
```

```
## [1] 260
```

```
##Chemical pollution
dataCP2 = subset(fulldat2, Global.Change.Driver == "CP")

##Remove all rows with NA values in ANY of the categories
dataCP2 = dataCP2 %>%
  filter(complete.cases(Habitat) &
           complete.cases(Enemy.type_further_reduced_Final) &
           complete.cases(Vector.borne) &
           complete.cases(Native.host.type_reduced_Final) & 
           complete.cases(Ectoparasite) &
           complete.cases(Vector.borne) &
           complete.cases(Free.living.stages) &
           complete.cases(RouteBoth.NA) &
           complete.cases(Human.ParasiteBoth.NA) &
           complete.cases(MacroparasiteBoth.NA) &
           complete.cases(Effect_Reduced_Update))

##Replicates for chemical pollution
nrow(dataCP2)
```

```
## [1] 253
```

###### Data Analysis

- Model selection output is for \(\Delta\) AIC < 2.
- Best models are AICc indicated best models, not including parsimony
  in best model.

```
##rma() wrapper for dredge
makeArgs.rma <- function(obj, termNames, comb, opt, 
                         ...) {
  ret <- MuMIn:::makeArgs.default(obj, termNames, comb, opt)
  names(ret)[1L] <- "mods"
  ret
}

##rma() wrapper for dredge
coefTable.rma <- function(model, ...) {
  MuMIn:::.makeCoefTable(model$b, model$se, coefNames = rownames(model$b))
}
```

##### Biodiversity Change

```
## Fixed term is "(Intercept)"
```

```
## 
## Call:
## model.avg(object = BioDiv_Dredge, subset = delta < 2)
## 
## Component model call: 
## rma.mv(yi = yi, V = vi2, mods = ~<38 unique rhs>, random = list(~1 | 
##      Citation.number, ~1 | ind_id), data = dataBL2, method = ML)
## 
## Component models: 
##                      df  logLik   AICc delta weight
## 3/4/6/9/10/12        13 -441.94 910.83  0.00   0.05
## 2/3/4/6/9/10/12      13 -441.94 910.83  0.00   0.05
## 3/4/6/8/9/10/12      14 -440.88 910.85  0.03   0.05
## 2/3/4/6/8/9/10/12    14 -440.88 910.85  0.03   0.05
## 4/6/8/9/10/12        13 -441.98 910.90  0.07   0.04
## 2/4/6/8/9/10/12      13 -441.98 910.90  0.07   0.04
## 3/4/6/8/9/11/12      14 -441.09 911.26  0.43   0.04
## 2/3/4/6/8/9/11/12    14 -441.09 911.26  0.43   0.04
## 4/6/9/10/12          12 -443.23 911.27  0.44   0.04
## 2/4/6/9/10/12        12 -443.23 911.27  0.44   0.04
## 4/6/8/9/11/12        13 -442.25 911.45  0.62   0.03
## 2/4/6/8/9/11/12      13 -442.25 911.45  0.62   0.03
## 1/3/4/6/9/10/12      14 -441.57 912.22  1.39   0.02
## 1/2/3/4/6/9/10/12    14 -441.57 912.22  1.39   0.02
## 2/3/4/6/9/10         12 -443.72 912.24  1.41   0.02
## 3/4/6/9/10           12 -443.72 912.24  1.41   0.02
## 4/6/8/9/12           12 -443.82 912.43  1.61   0.02
## 2/4/6/8/9/12         12 -443.82 912.43  1.61   0.02
## 4/5/6/8/9/12         13 -442.76 912.46  1.63   0.02
## 2/4/5/6/8/9/12       13 -442.76 912.46  1.63   0.02
## 2/3/4/5/6/8/9/12     14 -441.71 912.50  1.68   0.02
## 3/4/5/6/8/9/12       14 -441.71 912.50  1.68   0.02
## 3/4/5/6/9/10/12      14 -441.73 912.54  1.71   0.02
## 2/3/4/5/6/9/10/12    14 -441.73 912.54  1.71   0.02
## 3/4/6/8/9/10         13 -442.80 912.54  1.72   0.02
## 2/3/4/6/8/9/10       13 -442.80 912.54  1.72   0.02
## 3/4/6/8/9/12         13 -442.82 912.58  1.75   0.02
## 2/3/4/6/8/9/12       13 -442.82 912.58  1.75   0.02
## 3/4/6/8/9/10/11/12   15 -440.70 912.65  1.82   0.02
## 2/3/4/6/8/9/10/11/12 15 -440.70 912.65  1.82   0.02
## 2/3/4/6/7/8/9/10/12  15 -440.70 912.65  1.83   0.02
## 3/4/6/7/8/9/10/12    15 -440.70 912.65  1.83   0.02
## 4/6/7/8/9/10/12      14 -441.79 912.67  1.84   0.02
## 2/4/6/7/8/9/10/12    14 -441.79 912.67  1.84   0.02
## 4/6/8/9/10/11/12     14 -441.84 912.77  1.94   0.02
## 2/4/6/8/9/10/11/12   14 -441.84 912.77  1.94   0.02
## 1/2/3/4/6/8/9/10/12  15 -440.79 912.82  1.99   0.02
## 1/3/4/6/8/9/10/12    15 -440.79 912.82  1.99   0.02
## 
## Term codes: 
##                   Ectoparasite               Ectothermic.host 
##                              1                              2 
##          Effect_Reduced_Update         Endpoint_Host_Parasite 
##                              3                              4 
##             Free.living.stages                        Habitat 
##                              5                              6 
##          Human.ParasiteBoth.NA           MacroparasiteBoth.NA 
##                              7                              8 
## Native.host.type_reduced_Final                   RouteBoth.NA 
##                              9                             10 
##                   Vector.borne                   VenueNoModel 
##                             11                             12 
## 
## Model-averaged coefficients:  
## (full average) 
##                                         Estimate Std. Error z value Pr(>|z|)
## intrcpt                                 -0.09914    0.66510   0.149 0.881511
## Effect_Reduced_UpdateBiodiversity Loss   0.17764    0.19901   0.893 0.372055
## Endpoint_Host_ParasiteParasite           0.62299    0.22806   2.732 0.006300
## HabitatMarine                            1.65747    0.43007   3.854 0.000116
## HabitatTerrestrial                      -0.75147    0.47167   1.593 0.111111
## Native.host.type_reduced_FinalArthropod  0.85702    0.51285   1.671 0.094708
## Native.host.type_reduced_FinalMammal     0.42868    0.53718   0.798 0.424861
## Native.host.type_reduced_FinalMollusk    0.21247    0.34709   0.612 0.540441
## Native.host.type_reduced_FinalPlant      1.38927    0.48837   2.845 0.004445
## RouteBoth.NADirect                      -0.22214    0.20919   1.062 0.288280
## VenueNoModelLab                          0.76582    0.44606   1.717 0.086009
## Ectothermic.hostYes                     -0.42868    0.53718   0.798 0.424861
## MacroparasiteBoth.NAYes                  0.18273    0.17549   1.041 0.297754
## Vector.borneYes                          0.04277    0.11374   0.376 0.706927
## EctoparasiteYes                         -0.01224    0.07664   0.160 0.873131
## Free.living.stagesYes                   -0.01225    0.08301   0.148 0.882642
## Human.ParasiteBoth.NAYes                 0.01156    0.08114   0.142 0.886759
##                                            
## intrcpt                                    
## Effect_Reduced_UpdateBiodiversity Loss     
## Endpoint_Host_ParasiteParasite          ** 
## HabitatMarine                           ***
## HabitatTerrestrial                         
## Native.host.type_reduced_FinalArthropod .  
## Native.host.type_reduced_FinalMammal       
## Native.host.type_reduced_FinalMollusk      
## Native.host.type_reduced_FinalPlant     ** 
## RouteBoth.NADirect                         
## VenueNoModelLab                         .  
## Ectothermic.hostYes                        
## MacroparasiteBoth.NAYes                    
## Vector.borneYes                            
## EctoparasiteYes                            
## Free.living.stagesYes                      
## Human.ParasiteBoth.NAYes                   
##  
## (conditional average) 
##                                         Estimate Std. Error z value Pr(>|z|)
## intrcpt                                 -0.09914    0.66510   0.149 0.881511
## Effect_Reduced_UpdateBiodiversity Loss   0.28887    0.17965   1.608 0.107844
## Endpoint_Host_ParasiteParasite           0.62299    0.22806   2.732 0.006300
## HabitatMarine                            1.65747    0.43007   3.854 0.000116
## HabitatTerrestrial                      -0.75147    0.47167   1.593 0.111111
## Native.host.type_reduced_FinalArthropod  0.85702    0.51285   1.671 0.094708
## Native.host.type_reduced_FinalMammal     0.85736    0.45782   1.873 0.061107
## Native.host.type_reduced_FinalMollusk    0.21247    0.34709   0.612 0.540441
## Native.host.type_reduced_FinalPlant      1.38927    0.48837   2.845 0.004445
## RouteBoth.NADirect                      -0.31872    0.17890   1.782 0.074817
## VenueNoModelLab                          0.83673    0.39757   2.105 0.035323
## Ectothermic.hostYes                     -0.85736    0.45782   1.873 0.061107
## MacroparasiteBoth.NAYes                  0.26003    0.15402   1.688 0.091345
## Vector.borneYes                          0.19951    0.17054   1.170 0.242059
## EctoparasiteYes                         -0.15264    0.22765   0.670 0.502558
## Free.living.stagesYes                   -0.10200    0.21954   0.465 0.642224
## Human.ParasiteBoth.NAYes                 0.15650    0.25786   0.607 0.543894
##                                            
## intrcpt                                    
## Effect_Reduced_UpdateBiodiversity Loss     
## Endpoint_Host_ParasiteParasite          ** 
## HabitatMarine                           ***
## HabitatTerrestrial                         
## Native.host.type_reduced_FinalArthropod .  
## Native.host.type_reduced_FinalMammal    .  
## Native.host.type_reduced_FinalMollusk      
## Native.host.type_reduced_FinalPlant     ** 
## RouteBoth.NADirect                      .  
## VenueNoModelLab                         *  
## Ectothermic.hostYes                     .  
## MacroparasiteBoth.NAYes                 .  
## Vector.borneYes                            
## EctoparasiteYes                            
## Free.living.stagesYes                      
## Human.ParasiteBoth.NAYes                   
## ---
## Signif. codes:  0 '***' 0.001 '**' 0.01 '*' 0.05 '.' 0.1 ' ' 1
```

```
##                      Endpoint_Host_Parasite Habitat
## Sum of weights:      1.00                   1.00   
## N containing models:  167                    167   
##                      Native.host.type_reduced_Final VenueNoModel
## Sum of weights:      0.96                           0.85        
## N containing models:  154                            132        
##                      Effect_Reduced_Update MacroparasiteBoth.NA RouteBoth.NA
## Sum of weights:      0.64                  0.61                 0.59        
## N containing models:  111                   100                   94        
##                      Ectothermic.host Vector.borne Free.living.stages
## Sum of weights:      0.50             0.32         0.24              
## N containing models:   84               58           52              
##                      Ectoparasite Human.ParasiteBoth.NA
## Sum of weights:      0.20         0.18                 
## N containing models:   42           39
```

```
##  Native.host.type_reduced_Final emmean    SE  df asymp.LCL asymp.UCL
##  Amphibian/Reptile               0.469 0.383 Inf    -0.283      1.22
##  Arthropod                       1.080 0.434 Inf     0.230      1.93
##  Mammal                          1.299 0.299 Inf     0.713      1.88
##  Mollusk                         0.825 0.243 Inf     0.348      1.30
##  Plant                           1.820 0.322 Inf     1.190      2.45
## 
## Results are averaged over the levels of: Effect_Reduced_Update, Endpoint_Host_Parasite, Habitat, RouteBoth.NA 
## Confidence level used: 0.95
```

```
##  Endpoint_Host_Parasite emmean    SE  df asymp.LCL asymp.UCL
##  Host                    0.782 0.292 Inf      0.21      1.35
##  Parasite                1.415 0.180 Inf      1.06      1.77
## 
## Results are averaged over the levels of: Effect_Reduced_Update, Habitat, Native.host.type_reduced_Final, RouteBoth.NA 
## Confidence level used: 0.95
```

```
##  Effect_Reduced_Update emmean    SE  df asymp.LCL asymp.UCL
##  Biodiversity Gradient  0.909 0.245 Inf     0.429      1.39
##  Biodiversity Loss      1.288 0.208 Inf     0.880      1.70
## 
## Results are averaged over the levels of: Endpoint_Host_Parasite, Habitat, Native.host.type_reduced_Final, RouteBoth.NA 
## Confidence level used: 0.95
```

```
##  Habitat     emmean    SE  df asymp.LCL asymp.UCL
##  Freshwater   0.905 0.270 Inf     0.377     1.433
##  Marine       2.691 0.462 Inf     1.786     3.596
##  Terrestrial -0.301 0.244 Inf    -0.779     0.177
## 
## Results are averaged over the levels of: Effect_Reduced_Update, Endpoint_Host_Parasite, Native.host.type_reduced_Final, RouteBoth.NA 
## Confidence level used: 0.95
```

##### Climate Change

Cannot have “Enemy.type\_further\_reduced\_Final” and
“MacroparasiteBoth.NA” in the model; not uniquely determined. So,
dropped “macroparasite”.

```
## Fixed term is "(Intercept)"
```

```
## 
## Call:
## model.avg(object = ClimChange_Dredge, subset = delta < 2)
## 
## Component model call: 
## rma.mv(yi = yi, V = vi2, mods = ~<16 unique rhs>, random = list(~1 | 
##      Citation.number, ~1 | ind_id), data = dataCC2, method = ML)
## 
## Component models: 
##        df  logLik    AICc delta weight
## 3       4 -529.60 1067.35  0.00   0.11
## 23      5 -528.67 1067.56  0.20   0.10
## (Null)  3 -530.84 1067.76  0.41   0.09
## 234     6 -527.93 1068.15  0.80   0.07
## 38      5 -529.08 1068.37  1.02   0.07
## 237     6 -528.06 1068.42  1.07   0.07
## 36      5 -529.26 1068.75  1.39   0.06
## 2       4 -530.32 1068.77  1.42   0.05
## 35      5 -529.33 1068.88  1.53   0.05
## 236     6 -528.32 1068.94  1.59   0.05
## 37      5 -529.40 1069.01  1.66   0.05
## 2346    7 -527.30 1069.01  1.66   0.05
## 123     6 -528.39 1069.09  1.74   0.05
## 8       4 -530.53 1069.20  1.85   0.04
## 6       4 -530.54 1069.21  1.86   0.04
## 34      5 -529.54 1069.29  1.94   0.04
## 
## Term codes: 
##           Ectoparasite       Ectothermic.host Endpoint_Host_Parasite 
##                      1                      2                      3 
##     Free.living.stages  Human.ParasiteBoth.NA           RouteBoth.NA 
##                      4                      5                      6 
##           Vector.borne           VenueNoModel 
##                      7                      8 
## 
## Model-averaged coefficients:  
## (full average) 
##                                 Estimate Std. Error z value Pr(>|z|)
## intrcpt                         0.585068   0.392287   1.491    0.136
## Endpoint_Host_ParasiteParasite -0.306025   0.263878   1.160    0.246
## Ectothermic.hostYes            -0.199863   0.305539   0.654    0.513
## Free.living.stagesYes           0.065371   0.217031   0.301    0.763
## VenueNoModelLab                -0.024438   0.104592   0.234    0.815
## Vector.borneYes                -0.025825   0.111898   0.231    0.817
## RouteBoth.NADirect              0.057730   0.186647   0.309    0.757
## Human.ParasiteBoth.NAYes        0.013915   0.101722   0.137    0.891
## EctoparasiteYes                 0.008485   0.064993   0.131    0.896
##  
## (conditional average) 
##                                Estimate Std. Error z value Pr(>|z|)  
## intrcpt                          0.5851     0.3923   1.491    0.136  
## Endpoint_Host_ParasiteParasite  -0.3997     0.2313   1.728    0.084 .
## Ectothermic.hostYes             -0.4518     0.3118   1.449    0.147  
## Free.living.stagesYes            0.3936     0.3930   1.002    0.316  
## VenueNoModelLab                 -0.2188     0.2354   0.930    0.353  
## Vector.borneYes                 -0.2259     0.2536   0.891    0.373  
## RouteBoth.NADirect               0.2901     0.3281   0.884    0.377  
## Human.ParasiteBoth.NAYes         0.2671     0.3620   0.738    0.460  
## EctoparasiteYes                  0.1812     0.2427   0.747    0.455  
## ---
## Signif. codes:  0 '***' 0.001 '**' 0.01 '*' 0.05 '.' 0.1 ' ' 1
```

```
##                      Endpoint_Host_Parasite Ectothermic.host RouteBoth.NA
## Sum of weights:       0.69                   0.44             0.26       
## N containing models:    51                     34               25       
##                      Free.living.stages VenueNoModel Vector.borne
## Sum of weights:       0.24               0.22         0.21       
## N containing models:    22                 21           19       
##                      Human.ParasiteBoth.NA Ectoparasite Habitat
## Sum of weights:       0.19                  0.16         0.02  
## N containing models:    19                    17            3  
##                      Enemy.type_further_reduced_Final
## Sum of weights:      <0.01                           
## N containing models:     1
```

```
## 
## Multivariate Meta-Analysis Model (k = 284; method: REML)
## 
##    logLik   Deviance        AIC        BIC       AICc 
## -520.8188  1041.6376  1057.6376  1086.6585  1058.1729   
## 
## Variance Components:
## 
##             estim    sqrt  nlvls  fixed           factor 
## sigma^2.1  1.2614  1.1231    141     no  Citation.number 
## sigma^2.2  0.7102  0.8427    284     no           ind_id 
## 
## Test for Residual Heterogeneity:
## QE(df = 278) = 1053713.9798, p-val < .0001
## 
## Test of Moderators (coefficients 2:6):
## QM(df = 5) = 2.4429, p-val = 0.7851
## 
## Model Results:
## 
##                                                       estimate      se     zval 
## intrcpt                                                 0.1513  0.2732   0.5539 
## Effect_Reduced_UpdateCO2                                0.2540  0.3558   0.7138 
## Effect_Reduced_UpdateMean Temperature                   0.1174  0.3023   0.3884 
## Effect_Reduced_UpdatePrecipitation                     -0.1240  0.3900  -0.3179 
## Effect_Reduced_UpdateTemperature Variability or ENSO    0.5336  0.5831   0.9152 
## Effect_Reduced_UpdateUVB                                0.5904  0.7718   0.7650 
##                                                         pval    ci.lb   ci.ub 
## intrcpt                                               0.5797  -0.3841  0.6868 
## Effect_Reduced_UpdateCO2                              0.4753  -0.4434  0.9514 
## Effect_Reduced_UpdateMean Temperature                 0.6977  -0.4751  0.7099 
## Effect_Reduced_UpdatePrecipitation                    0.7505  -0.8883  0.6403 
## Effect_Reduced_UpdateTemperature Variability or ENSO  0.3601  -0.6092  1.6764 
## Effect_Reduced_UpdateUVB                              0.4443  -0.9223  2.1031 
##  
## intrcpt 
## Effect_Reduced_UpdateCO2 
## Effect_Reduced_UpdateMean Temperature 
## Effect_Reduced_UpdatePrecipitation 
## Effect_Reduced_UpdateTemperature Variability or ENSO 
## Effect_Reduced_UpdateUVB 
## 
## ---
## Signif. codes:  0 '***' 0.001 '**' 0.01 '*' 0.05 '.' 0.1 ' ' 1
```

##### Habitat Loss and Change

```
## Fixed term is "(Intercept)"
```

```
## 
## Call:
## model.avg(object = HabLoss_Dredge, subset = delta < 2)
## 
## Component model call: 
## rma.mv(yi = yi, V = vi2, mods = ~<44 unique rhs>, random = list(~1 | 
##      Citation.number, ~1 | ind_id), data = dataHLC2, method = ML)
## 
## Component models: 
##        df  logLik    AICc delta weight
## 2789    7 -585.81 1185.91  0.00   0.04
## 1238   12 -580.60 1186.01  0.10   0.04
## 789     6 -586.94 1186.10  0.20   0.04
## 138    11 -581.72 1186.13  0.22   0.04
## 289     6 -586.98 1186.17  0.26   0.04
## 12789   9 -583.90 1186.26  0.36   0.03
## 1789    8 -585.07 1186.52  0.61   0.03
## 89      5 -588.23 1186.61  0.70   0.03
## 1278    8 -585.15 1186.67  0.76   0.03
## 12389  13 -579.98 1186.91  1.00   0.03
## 278     6 -587.36 1186.93  1.02   0.02
## 178     7 -586.34 1186.96  1.05   0.02
## 248     6 -587.40 1187.02  1.11   0.02
## 1389   12 -581.12 1187.04  1.13   0.02
## 5789    7 -586.41 1187.11  1.21   0.02
## 78      5 -588.50 1187.16  1.25   0.02
## 1289    8 -585.41 1187.19  1.28   0.02
## 589     6 -587.50 1187.22  1.31   0.02
## 1368   12 -581.22 1187.25  1.35   0.02
## 12368  13 -580.17 1187.29  1.38   0.02
## 12348  13 -580.19 1187.33  1.42   0.02
## 2478    7 -586.55 1187.38  1.47   0.02
## 1358   12 -581.29 1187.39  1.48   0.02
## 15789   9 -584.47 1187.39  1.49   0.02
## 12478   9 -584.47 1187.40  1.50   0.02
## 48      5 -588.63 1187.42  1.51   0.02
## 1348   12 -581.32 1187.45  1.55   0.02
## 2589    7 -586.60 1187.48  1.58   0.02
## 25789   8 -585.56 1187.48  1.58   0.02
## 12678   9 -584.54 1187.54  1.64   0.02
## 478     6 -587.68 1187.57  1.67   0.02
## 126789 10 -583.52 1187.60  1.69   0.02
## 1678    8 -585.63 1187.63  1.72   0.02
## 1478    8 -585.65 1187.66  1.76   0.02
## 1248    8 -585.65 1187.67  1.76   0.02
## 16789   9 -584.61 1187.68  1.78   0.02
## 189     7 -586.70 1187.69  1.78   0.02
## 2489    7 -586.71 1187.71  1.81   0.02
## 26789   8 -585.69 1187.75  1.84   0.02
## 125789 10 -583.60 1187.76  1.85   0.02
## 12358  13 -580.41 1187.76  1.86   0.02
## 12378  13 -580.44 1187.83  1.92   0.02
## 6789    7 -586.78 1187.84  1.94   0.02
## 24789   8 -585.76 1187.88  1.98   0.02
## 
## Term codes: 
##            Effect_Reduced_Update           Endpoint_Host_Parasite 
##                                1                                2 
## Enemy.type_further_reduced_Final               Free.living.stages 
##                                3                                4 
##                          Habitat            Human.ParasiteBoth.NA 
##                                5                                6 
##             MacroparasiteBoth.NA                     RouteBoth.NA 
##                                7                                8 
##                     Vector.borne 
##                                9 
## 
## Model-averaged coefficients:  
## (full average) 
##                                           Estimate Std. Error z value Pr(>|z|)
## intrcpt                                    0.20807    0.66836   0.311   0.7556
## Endpoint_Host_ParasiteParasite            -0.46159    0.59176   0.780   0.4354
## MacroparasiteBoth.NAYes                   -0.11722    0.16497   0.711   0.4774
## RouteBoth.NADirect                         0.42256    0.20051   2.107   0.0351
## Vector.borneYes                            0.16784    0.21609   0.777   0.4373
## Effect_Reduced_UpdateForest Fragmentation -0.01863    0.22871   0.081   0.9351
## Effect_Reduced_UpdateUrbanization         -0.21940    0.25302   0.867   0.3859
## Enemy.type_further_reduced_FinalBacteria   0.07228    0.18587   0.389   0.6974
## Enemy.type_further_reduced_FinalFungi     -0.08370    0.22900   0.366   0.7147
## Enemy.type_further_reduced_FinalHelminth   0.01925    0.12687   0.152   0.8794
## Enemy.type_further_reduced_FinalProtist    0.05553    0.16864   0.329   0.7419
## Enemy.type_further_reduced_FinalVirus      0.13955    0.27885   0.500   0.6167
## Free.living.stagesYes                     -0.04429    0.12301   0.360   0.7188
## HabitatTerrestrial                        -0.04601    0.16784   0.274   0.7840
## Human.ParasiteBoth.NAYes                   0.01635    0.06263   0.261   0.7941
##                                            
## intrcpt                                    
## Endpoint_Host_ParasiteParasite             
## MacroparasiteBoth.NAYes                    
## RouteBoth.NADirect                        *
## Vector.borneYes                            
## Effect_Reduced_UpdateForest Fragmentation  
## Effect_Reduced_UpdateUrbanization          
## Enemy.type_further_reduced_FinalBacteria   
## Enemy.type_further_reduced_FinalFungi      
## Enemy.type_further_reduced_FinalHelminth   
## Enemy.type_further_reduced_FinalProtist    
## Enemy.type_further_reduced_FinalVirus      
## Free.living.stagesYes                      
## HabitatTerrestrial                         
## Human.ParasiteBoth.NAYes                   
##  
## (conditional average) 
##                                           Estimate Std. Error z value Pr(>|z|)
## intrcpt                                    0.20807    0.66836   0.311   0.7556
## Endpoint_Host_ParasiteParasite            -0.84339    0.56376   1.496   0.1347
## MacroparasiteBoth.NAYes                   -0.21985    0.16877   1.303   0.1927
## RouteBoth.NADirect                         0.42256    0.20051   2.107   0.0351
## Vector.borneYes                            0.32466    0.19852   1.635   0.1020
## Effect_Reduced_UpdateForest Fragmentation -0.03218    0.29987   0.107   0.9145
## Effect_Reduced_UpdateUrbanization         -0.37897    0.22385   1.693   0.0905
## Enemy.type_further_reduced_FinalBacteria   0.27818    0.27513   1.011   0.3120
## Enemy.type_further_reduced_FinalFungi     -0.32216    0.35358   0.911   0.3622
## Enemy.type_further_reduced_FinalHelminth   0.07408    0.24060   0.308   0.7582
## Enemy.type_further_reduced_FinalProtist    0.21373    0.27504   0.777   0.4371
## Enemy.type_further_reduced_FinalVirus      0.53713    0.29279   1.835   0.0666
## Free.living.stagesYes                     -0.21313    0.19190   1.111   0.2667
## HabitatTerrestrial                        -0.29703    0.32755   0.907   0.3645
## Human.ParasiteBoth.NAYes                   0.11235    0.12716   0.884   0.3770
##                                            
## intrcpt                                    
## Endpoint_Host_ParasiteParasite             
## MacroparasiteBoth.NAYes                    
## RouteBoth.NADirect                        *
## Vector.borneYes                            
## Effect_Reduced_UpdateForest Fragmentation  
## Effect_Reduced_UpdateUrbanization         .
## Enemy.type_further_reduced_FinalBacteria   
## Enemy.type_further_reduced_FinalFungi      
## Enemy.type_further_reduced_FinalHelminth   
## Enemy.type_further_reduced_FinalProtist    
## Enemy.type_further_reduced_FinalVirus     .
## Free.living.stagesYes                      
## HabitatTerrestrial                         
## Human.ParasiteBoth.NAYes                   
## ---
## Signif. codes:  0 '***' 0.001 '**' 0.01 '*' 0.05 '.' 0.1 ' ' 1
```

```
##                      RouteBoth.NA Effect_Reduced_Update Endpoint_Host_Parasite
## Sum of weights:      0.97         0.61                  0.50                  
## N containing models:  201          131                   105                  
##                      Vector.borne MacroparasiteBoth.NA Free.living.stages
## Sum of weights:      0.48         0.47                 0.31              
## N containing models:   99           96                   75              
##                      Enemy.type_further_reduced_Final Habitat
## Sum of weights:      0.28                             0.28   
## N containing models:   61                               72   
##                      Human.ParasiteBoth.NA Ectothermic.host
## Sum of weights:      0.27                  0.16            
## N containing models:   72                    49            
##                      Native.host.type_reduced_Final
## Sum of weights:      0.01                          
## N containing models:    4
```

```
##  contrast estimate    SE  df z.ratio p.value
##  No - Yes   -0.334 0.191 Inf -1.743  0.0814 
## 
## Results are averaged over the levels of: Endpoint_Host_Parasite, RouteBoth.NA, MacroparasiteBoth.NA
```

```
##  contrast        estimate    SE  df z.ratio p.value
##  Host - Parasite     0.85 0.569 Inf 1.494   0.1352 
## 
## Results are averaged over the levels of: Vector.borne, RouteBoth.NA, MacroparasiteBoth.NA
```

```
##  contrast         estimate    SE  df z.ratio p.value
##  Complex - Direct   -0.537 0.184 Inf -2.921  0.0035 
## 
## Results are averaged over the levels of: Endpoint_Host_Parasite, Vector.borne, MacroparasiteBoth.NA
```

```
##  contrast estimate   SE  df z.ratio p.value
##  No - Yes    0.198 0.13 Inf 1.529   0.1262 
## 
## Results are averaged over the levels of: Endpoint_Host_Parasite, Vector.borne, RouteBoth.NA
```

##### Introduced Species

```
## Fixed term is "(Intercept)"
```

```
## 
## Call:
## model.avg(object = IntroSpec_Dredge, subset = delta < 2)
## 
## Component model call: 
## rma.mv(yi = yi, V = vi2, mods = ~<9 unique rhs>, random = list(~1 | 
##      Citation.number, ~1 | ind_id), data = dataIS2, method = ML)
## 
## Component models: 
##                  df  logLik   AICc delta weight
## 4/6/7/8           8 -353.96 724.50  0.00   0.23
## 4/6/7/8/11        9 -353.62 725.95  1.45   0.11
## 4/5/6/7/8/9/11   19 -342.45 726.07  1.57   0.10
## 2/4/5/6/7/8/9/11 19 -342.45 726.07  1.57   0.10
## 4/6/7/8/10        9 -353.73 726.19  1.68   0.10
## 1/4/6/8           7 -355.92 726.29  1.78   0.09
## 3/4/5/7/8/9/10   20 -341.43 726.38  1.88   0.09
## 2/3/4/5/7/8/9/10 20 -341.43 726.38  1.88   0.09
## 4/6/7/8/10/11    10 -352.80 726.49  1.99   0.08
## 
## Term codes: 
##                     Ectoparasite                 Ectothermic.host 
##                                1                                2 
##            Effect_Reduced_Update           Endpoint_Host_Parasite 
##                                3                                4 
## Enemy.type_further_reduced_Final               Free.living.stages 
##                                5                                6 
##                          Habitat            Human.ParasiteBoth.NA 
##                                7                                8 
##   Native.host.type_reduced_Final                     RouteBoth.NA 
##                                9                               10 
##                     Vector.borne 
##                               11 
## 
## Model-averaged coefficients:  
## (full average) 
##                                           Estimate Std. Error z value Pr(>|z|)
## intrcpt                                  -1.552218   3.512712   0.442   0.6586
## Endpoint_Host_ParasiteParasite            1.559590   0.314694   4.956    7e-07
## Free.living.stagesYes                     1.608947   1.314854   1.224   0.2211
## HabitatMarine                             1.955138   2.290846   0.853   0.3934
## HabitatTerrestrial                       -0.112979   1.309367   0.086   0.9312
## Human.ParasiteBoth.NAYes                 -3.202654   1.707674   1.875   0.0607
## Vector.borneYes                           0.481459   0.817120   0.589   0.5557
## Enemy.type_further_reduced_FinalFungi    -0.820298   1.133066   0.724   0.4691
## Enemy.type_further_reduced_FinalHelminth -0.640203   1.080893   0.592   0.5537
## Enemy.type_further_reduced_FinalProtist  -1.654719   2.759751   0.600   0.5488
## Enemy.type_further_reduced_FinalVirus    -0.678450   1.407048   0.482   0.6297
## Native.host.type_reduced_FinalArthropod   0.436324   1.316011   0.332   0.7402
## Native.host.type_reduced_FinalBird        0.759340   1.784216   0.426   0.6704
## Native.host.type_reduced_FinalFish       -0.008497   1.064225   0.008   0.9936
## Native.host.type_reduced_FinalMammal      0.651945   1.502779   0.434   0.6644
## Native.host.type_reduced_FinalMollusk    -1.940185   2.594877   0.748   0.4546
## Native.host.type_reduced_FinalPlant       1.219329   2.090511   0.583   0.5597
## Ectothermic.hostYes                      -0.651945   1.502779   0.434   0.6644
## RouteBoth.NADirect                       -0.193147   0.732881   0.264   0.7921
## EctoparasiteYes                           0.037685   0.137040   0.275   0.7833
## Effect_Reduced_UpdateSpillback           -0.224674   0.513075   0.438   0.6615
## Effect_Reduced_UpdateSpillover           -0.163093   0.380554   0.429   0.6682
##                                             
## intrcpt                                     
## Endpoint_Host_ParasiteParasite           ***
## Free.living.stagesYes                       
## HabitatMarine                               
## HabitatTerrestrial                          
## Human.ParasiteBoth.NAYes                 .  
## Vector.borneYes                             
## Enemy.type_further_reduced_FinalFungi       
## Enemy.type_further_reduced_FinalHelminth    
## Enemy.type_further_reduced_FinalProtist     
## Enemy.type_further_reduced_FinalVirus       
## Native.host.type_reduced_FinalArthropod     
## Native.host.type_reduced_FinalBird          
## Native.host.type_reduced_FinalFish          
## Native.host.type_reduced_FinalMammal        
## Native.host.type_reduced_FinalMollusk       
## Native.host.type_reduced_FinalPlant         
## Ectothermic.hostYes                         
## RouteBoth.NADirect                          
## EctoparasiteYes                             
## Effect_Reduced_UpdateSpillback              
## Effect_Reduced_UpdateSpillover              
##  
## (conditional average) 
##                                          Estimate Std. Error z value Pr(>|z|)
## intrcpt                                  -1.55222    3.51271   0.442 0.658572
## Endpoint_Host_ParasiteParasite            1.55959    0.31469   4.956    7e-07
## Free.living.stagesYes                     1.95785    1.19191   1.643 0.100461
## HabitatMarine                             2.15640    2.31391   0.932 0.351374
## HabitatTerrestrial                       -0.12461    1.37458   0.091 0.927769
## Human.ParasiteBoth.NAYes                 -3.20265    1.70767   1.875 0.060731
## Vector.borneYes                           1.19606    0.89665   1.334 0.182229
## Enemy.type_further_reduced_FinalFungi    -2.12471    0.74421   2.855 0.004304
## Enemy.type_further_reduced_FinalHelminth -1.65824    1.15673   1.434 0.151700
## Enemy.type_further_reduced_FinalProtist  -4.28601    2.90682   1.474 0.140356
## Enemy.type_further_reduced_FinalVirus    -1.75730    1.79781   0.977 0.328337
## Native.host.type_reduced_FinalArthropod   1.13016    1.92399   0.587 0.556934
## Native.host.type_reduced_FinalBird        1.96682    2.42296   0.812 0.416938
## Native.host.type_reduced_FinalFish       -0.02201    1.71268   0.013 0.989747
## Native.host.type_reduced_FinalMammal      3.37731    1.57944   2.138 0.032493
## Native.host.type_reduced_FinalMollusk    -5.02542    1.39143   3.612 0.000304
## Native.host.type_reduced_FinalPlant       3.15828    2.27946   1.386 0.165889
## Ectothermic.hostYes                      -3.37731    1.57944   2.138 0.032493
## RouteBoth.NADirect                       -0.53559    1.14280   0.469 0.639310
## EctoparasiteYes                           0.40377    0.23109   1.747 0.080591
## Effect_Reduced_UpdateSpillback           -1.26073    0.41350   3.049 0.002297
## Effect_Reduced_UpdateSpillover           -0.91518    0.35265   2.595 0.009455
##                                             
## intrcpt                                     
## Endpoint_Host_ParasiteParasite           ***
## Free.living.stagesYes                       
## HabitatMarine                               
## HabitatTerrestrial                          
## Human.ParasiteBoth.NAYes                 .  
## Vector.borneYes                             
## Enemy.type_further_reduced_FinalFungi    ** 
## Enemy.type_further_reduced_FinalHelminth    
## Enemy.type_further_reduced_FinalProtist     
## Enemy.type_further_reduced_FinalVirus       
## Native.host.type_reduced_FinalArthropod     
## Native.host.type_reduced_FinalBird          
## Native.host.type_reduced_FinalFish          
## Native.host.type_reduced_FinalMammal     *  
## Native.host.type_reduced_FinalMollusk    ***
## Native.host.type_reduced_FinalPlant         
## Ectothermic.hostYes                      *  
## RouteBoth.NADirect                          
## EctoparasiteYes                          .  
## Effect_Reduced_UpdateSpillback           ** 
## Effect_Reduced_UpdateSpillover           ** 
## ---
## Signif. codes:  0 '***' 0.001 '**' 0.01 '*' 0.05 '.' 0.1 ' ' 1
```

```
##                      Endpoint_Host_Parasite Human.ParasiteBoth.NA
## Sum of weights:      1.00                   1.00                 
## N containing models:   53                     53                 
##                      Free.living.stages Habitat
## Sum of weights:      0.75               0.62   
## N containing models:   40                 30   
##                      Enemy.type_further_reduced_Final Effect_Reduced_Update
## Sum of weights:      0.56                             0.47                 
## N containing models:   32                               28                 
##                      RouteBoth.NA Native.host.type_reduced_Final
## Sum of weights:      0.42         0.41                          
## N containing models:   24           22                          
##                      Ectothermic.host Vector.borne Ectoparasite
## Sum of weights:      0.38             0.29         0.21        
## N containing models:   22               16           14
```

```
##  contrast        estimate    SE  df z.ratio p.value
##  Host - Parasite    -1.55 0.306 Inf -5.045  <.0001 
## 
## Results are averaged over the levels of: Free.living.stages, Habitat, Human.ParasiteBoth.NA
```

```
##  contrast estimate    SE  df z.ratio p.value
##  No - Yes     -1.4 0.387 Inf -3.628  0.0003 
## 
## Results are averaged over the levels of: Endpoint_Host_Parasite, Habitat, Human.ParasiteBoth.NA
```

```
##  contrast                 estimate    SE  df z.ratio p.value
##  Freshwater - Marine        -0.691 0.474 Inf -1.456  0.3122 
##  Freshwater - Terrestrial   -0.850 0.331 Inf -2.567  0.0277 
##  Marine - Terrestrial       -0.160 0.390 Inf -0.410  0.9114 
## 
## Results are averaged over the levels of: Endpoint_Host_Parasite, Free.living.stages, Human.ParasiteBoth.NA 
## P value adjustment: tukey method for comparing a family of 3 estimates
```

```
##  contrast estimate    SE  df z.ratio p.value
##  No - Yes     2.31 0.871 Inf 2.657   0.0079 
## 
## Results are averaged over the levels of: Endpoint_Host_Parasite, Free.living.stages, Habitat
```

##### Chemical Pollution

```
## Fixed term is "(Intercept)"
```

```
## 
## Call:
## model.avg(object = ChemPoll_Dredge, subset = delta < 2)
## 
## Component model call: 
## rma.mv(yi = yi, V = vi2, mods = ~<24 unique rhs>, random = list(~1 | 
##      Citation.number, ~1 | ind_id), data = dataCP2, method = ML)
## 
## Component models: 
##        df  logLik   AICc delta weight
## 1       4 -436.90 881.95  0.00   0.08
## 8       4 -437.11 882.39  0.43   0.06
## 5       4 -437.12 882.40  0.45   0.06
## 58      5 -436.08 882.41  0.46   0.06
## 45      6 -435.17 882.68  0.73   0.05
## (Null)  3 -438.34 882.77  0.82   0.05
## 258     6 -435.29 882.92  0.97   0.05
## 15      5 -436.37 882.98  1.02   0.05
## 3       4 -437.44 883.05  1.09   0.04
## 35      5 -436.41 883.07  1.12   0.04
## 38      5 -436.52 883.29  1.34   0.04
## 458     7 -434.43 883.31  1.36   0.04
## 18      5 -436.54 883.32  1.37   0.04
## 13      5 -436.55 883.34  1.39   0.04
## 358     6 -435.54 883.42  1.46   0.04
## 56      5 -436.71 883.66  1.71   0.03
## 17      5 -436.73 883.69  1.74   0.03
## 6       4 -437.81 883.77  1.82   0.03
## 16      5 -436.78 883.79  1.84   0.03
## 457     7 -434.71 883.87  1.92   0.03
## 2358    7 -434.73 883.91  1.96   0.03
## 12      5 -436.85 883.94  1.98   0.03
## 25      5 -436.85 883.94  1.99   0.03
## 28      5 -436.85 883.95  1.99   0.03
## 
## Term codes: 
##           Ectoparasite       Ectothermic.host Endpoint_Host_Parasite 
##                      1                      2                      3 
##                Habitat  Human.ParasiteBoth.NA   MacroparasiteBoth.NA 
##                      4                      5                      6 
##           RouteBoth.NA           Vector.borne 
##                      7                      8 
## 
## Model-averaged coefficients:  
## (full average) 
##                                 Estimate Std. Error z value Pr(>|z|)
## intrcpt                        -0.021522   0.364646   0.059    0.953
## EctoparasiteYes                -0.064581   0.128759   0.502    0.616
## Vector.borneYes                 0.100503   0.167064   0.602    0.547
## Human.ParasiteBoth.NAYes        0.407690   0.575410   0.709    0.479
## HabitatMarine                   0.021872   0.139178   0.157    0.875
## HabitatTerrestrial             -0.034042   0.107854   0.316    0.752
## Ectothermic.hostYes             0.065223   0.316945   0.206    0.837
## Endpoint_Host_ParasiteParasite  0.042585   0.107750   0.395    0.693
## MacroparasiteBoth.NAYes         0.015872   0.075077   0.211    0.833
## RouteBoth.NADirect             -0.007248   0.044682   0.162    0.871
##  
## (conditional average) 
##                                Estimate Std. Error z value Pr(>|z|)  
## intrcpt                        -0.02152    0.36465   0.059    0.953  
## EctoparasiteYes                -0.22387    0.14769   1.516    0.130  
## Vector.borneYes                 0.26586    0.17283   1.538    0.124  
## Human.ParasiteBoth.NAYes        0.80948    0.57633   1.405    0.160  
## HabitatMarine                   0.18143    0.36295   0.500    0.617  
## HabitatTerrestrial             -0.28239    0.16237   1.739    0.082 .
## Ectothermic.hostYes             0.40768    0.69877   0.583    0.560  
## Endpoint_Host_ParasiteParasite  0.18537    0.15514   1.195    0.232  
## MacroparasiteBoth.NAYes         0.17014    0.18487   0.920    0.357  
## RouteBoth.NADirect             -0.11874    0.13953   0.851    0.395  
## ---
## Signif. codes:  0 '***' 0.001 '**' 0.01 '*' 0.05 '.' 0.1 ' ' 1
```

```
##                      Human.ParasiteBoth.NA Vector.borne Ectoparasite
## Sum of weights:      0.54                  0.41         0.28        
## N containing models:   74                    57           40        
##                      Ectothermic.host Endpoint_Host_Parasite Habitat
## Sum of weights:      0.28             0.27                   0.27   
## N containing models:   46               40                     44   
##                      RouteBoth.NA MacroparasiteBoth.NA Free.living.stages
## Sum of weights:      0.20         0.19                 0.14              
## N containing models:   37           33                   25              
##                      Native.host.type_reduced_Final Effect_Reduced_Update
## Sum of weights:      0.06                           0.02                 
## N containing models:   12                              4
```

```
## 
## Multivariate Meta-Analysis Model (k = 253; method: REML)
## 
##    logLik   Deviance        AIC        BIC       AICc 
## -434.8114   869.6228   877.6228   891.7246   877.7854   
## 
## Variance Components:
## 
##             estim    sqrt  nlvls  fixed           factor 
## sigma^2.1  0.1159  0.3405     67     no  Citation.number 
## sigma^2.2  0.6377  0.7986    253     no           ind_id 
## 
## Test for Residual Heterogeneity:
## QE(df = 251) = 1176.6115, p-val < .0001
## 
## Test of Moderators (coefficient 2):
## QM(df = 1) = 2.3149, p-val = 0.1281
## 
## Model Results:
## 
##                  estimate      se     zval    pval    ci.lb   ci.ub 
## intrcpt            0.1772  0.1117   1.5864  0.1126  -0.0417  0.3961    
## EctoparasiteYes   -0.2515  0.1653  -1.5215  0.1281  -0.5755  0.0725    
## 
## ---
## Signif. codes:  0 '***' 0.001 '**' 0.01 '*' 0.05 '.' 0.1 ' ' 1
```
